# A cytochrome P450 is essential for cardenolide biosynthesis in *Chrysolina* beetles

**DOI:** 10.64898/2026.11.19.767920

**Authors:** Houchao Xu, Angeliki Stathaki, Matilde Florean, Song Wu, Katrin Luck, Heiko Vogel, Maike L. Fischer, Yoko Nakamura, Martin Kaltenpoth, Tobias Köllner, Sarah E. O’Connor

## Abstract

Cardenolides are naturally occurring steroidal derivatives that function as chemical defense compounds. These natural products are also widely used in medicine, as exemplified by the use of digitoxin in treating cardiac disease. Although cardenolides are most widely known as plant-derived natural products, these compounds are also produced in certain beetle species. However, while the biosynthesis of cardenolides has been well-studied in plants, it is not known how insects produce these compounds. Here we report the identification of a cardenolide biosynthetic enzyme from the leaf beetle *Chrysolina coerulans*. This cytochrome P450, *Cc*CYP9Z273, catalyzes an array of steroid oxidations, including the conversion of pregnenolone into caraumbellogenin through C14β-hydroxylation. Gene silencing suggests that *Cc*CYP9Z273 is essential for beetle cardenolide formation and isotope tracing experiments support caraumbellogenin as an on-pathway biosynthetic intermediate. Mutagenesis of *Cc*CYP9Z273 identifies an active-site residue that controls the enzyme product selectivity between C14- and C15-hydroxylation. These findings establish an early step in beetle cardenolide biosynthesis and highlight how distinct enzyme families have been recruited in insects and plants to generate a shared intermediate in the biosynthesis of these important metabolites.

## Introduction

Cardenolides are steroidal natural products bearing a five-membered α,β-unsaturated lactone at C17 and, typically, a 14β-hydroxyl group.^1^ The glycosylated derivatives of cardenolides, also categorized as cardiac glycosides, are best known as inhibitors of Na+/K+-ATPase enzymes. This inhibitory activity underlies the mechanistic basis by which these compounds function as treatments in cardiac disorders.^2^ Cardenolides occur predominantly in plants and are distributed across distantly related lineages, including *Digitalis* (Plantaginaceae), *Calotropis* and *Periploca* (Apocynaceae), and *Erysimum* (Brassicaceae).^3–5^ Representative compounds include digitoxin (**1**) from *Digitalis purpurea*, which is used in the treatment of heart failure,^6^ and erysimoside (**2**) from *Erysimum cheiranthoides* (Fig. 1A), which confers to the plant resistance against herbivores.^4^

**Fig. 1.**
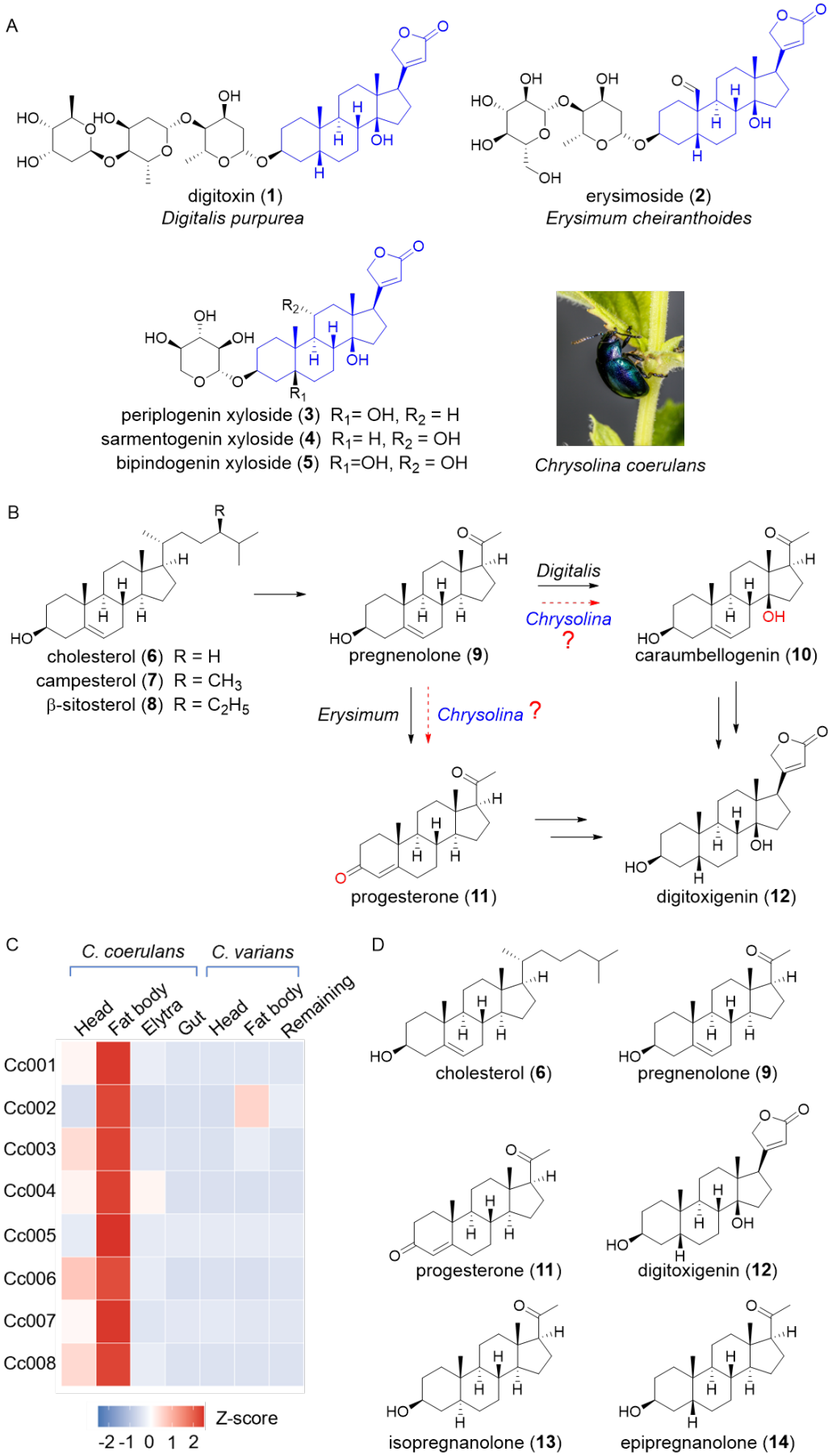
(A) Representative cardiac glycosides from plants and *Chrysolina* beetles. The steroidal aglycone moieties are highlighted in blue. (B) Simplified biosynthetic pathways to the cardenolide aglycone digitoxigenin. (C) Tissue-specific expression profiles of the top eight candidate P450 genes in *C. coerulans*. Expression profiles of the nearest homologs from the no-producer, *C. varians*, are included for comparison. The heatmap shows normalized transcript abundance (Z-scores) across different tissues and body samples. Red and blue indicate relatively high and low expression levels, respectively. Cc001 represents *Cc*CYP9Z273. (D) Putative intermediates in cardenolide biosynthesis tested with heterologously expressed enzymes.

Recent studies have elucidated the genetic and enzymatic basis of cardenolide biosynthesis in plants. The pathway is initiated by oxidative side-chain cleavage of either cholesterol (**6**) or a phytosterol, such as campesterol (**7**) and β-sitosterol (**8**), to yield the C_21_ intermediate pregnenolone (**9**; Fig. 1B).^3–5, 10–11^ After pregnenolone (**9**) formation, the order of steroid modification differs among the characterized plant pathways. In both *Digitalis purpurea* and *Periploca sepium*, a 2-oxoglutarate/Fe(II)-dependent steroid 14β-hydroxylase (S14βH and *Ps*14βPH, respectively) converts **9** directly into caraumbellogenin (**10**), supporting an early C14β-hydroxylation step in both species.^12 5^ In contrast, biochemical and genetic evidence from *Erysimum cheiranthoides* supports conversion of **9** into progesterone (**11**) through 3β-hydroxyl oxidation and Δ^5^-to-Δ^4^ double-bond isomerization, catalyzed by the enzymes *Ec*3βHSD and *Ec*3KSI.^13^ Thus, the known plant pathways differ in the sequence of biosynthetic steps immediately downstream of pregnenolone (**9**) (Fig. 1B).

Notably, cardenolides are also found in insects. Typically, sequestration after feeding on a cardenolide-producing plant is the underlying mechanism by which cardenolides occur in these invertebrates. For example, monarch butterflies (*Danaus plexippus*) sequester cardenolides acquired through larval feeding on milkweeds, whereas the oleander bug *Caenocoris nerii* acquires and sequesters these natural products by feeding on *Nerium oleander* plants.^7,8^ However, in some cases, insects produce cardenolides de novo. Leaf beetles of the genus *Chrysolina* are one such example: these beetles produce cardenolides despite feeding on plants established as non-producers of cardenolides.^9^ In the mint-feeding species *Chrysolina coerulans* (Fig. 1A), the major cardenolides identified in defensive secretions include periplogenin xyloside (**3**), sarmentogenin xyloside (**4**) and bipindogenin xyloside (**5**), together with the corresponding aglycones (Fig. 1A).^9^ Labeling experiments in *C. coerulans* have demonstrated that isotopically labeled cholesterol (**6**) is converted to cardenolides through a C_21_ intermediate.^14^ However, the pathway underlying cardenolide production in *Chrysolina* remains otherwise uncharacterized. One of the key unresolved questions is whether pregnenolone (**9**) undergoes C14β-hydroxylation to caraumbellogenin (**10**), as in *Digitalis* and *Periploca* plants, or is converted into progesterone (**11**), as is observed in *Erysimum* plants. Resolving this branch point would provide a foundation for elucidating the beetle cardenolide pathway and comparing the enzymes recruited by plants and insects for specialized metabolism.

Here we combine isotope-labeled precursor feeding, gene silencing and in vitro assays to identify the enzyme that catalyzes this early step of cardenolide biosynthesis in *Chrysolina* beetles. We identify a cytochrome P450 that catalyzes the C14β-hydroxylation of pregnenolone (**9**) to yield caraumbellogenin (**10**), and we establish the biosynthetic role of this enzymatic product by gene silencing in beetles. Feeding of isotopically labeled caraumbellogenin (**10**) also supports the conclusion that this metabolite is an on-pathway intermediate in cardenolide biosynthesis in leaf beetles. This work demonstrates that the cardenolide biosynthetic pathways found in both plants and insects share a conserved steroid transformation, even though this reaction is catalyzed by unrelated enzyme families.

## Results

### Identification and biochemical characterization of *Cc*CYP9Z273

To identify enzymes involved in cardenolide biosynthesis, we first examined cardenolide distribution across adult tissues and developmental stages in *C. coerulans*, which were collected locally in Jena, Germany. Cardenolides were detected in methanolic extracts of all body parts examined in both male and female adults, as well as in larvae, although the specific profile of individual cardenolides differed among these samples (Fig. S01). This widespread distribution provided no clear indication of a tissue-specific site of biosynthesis. We therefore prioritized genes expressed in the fat body, since this organ has been implicated in production of defensive metabolites in other leaf beetle species.^15,16^ Additionally, in this initial screen, we limited our focus to genes annotated as cytochrome P450s, since we rationalized that conversion of pregnenolone (**9**) would most likely involve an oxidative transformation. Guided by these two assumptions, we selected eight P450 candidates with enriched expression in the fat body of *C. coerulans* for functional screening. We also included tissue-resolved transcriptomic data from the non-cardenolide-producing related species, *Chrysolina varians*, as a comparative reference (Fig. 1C and Fig. S02).^17^

We separately co-expressed each of the eight P450 candidates with a cytochrome P450 reductase (CPR, Table S01) in *Saccharomyces cerevisiae* and prepared the corresponding microsomal fractions for in vitro enzyme assays. Microsomal fractions prepared from yeast expressing CPR alone were used as negative controls. The microsomes were incubated with pregnenolone (**9**), as well as a variety of other available substrates that could likely serve as biosynthetic intermediates: cholesterol (**6**), progesterone (**11**), digitoxigenin (**12**), isopregnanolone (**13**), and epipregnanolone (**14**), which each represent a potential intermediate in the cardenolide pathway (Fig. 1D). From this screen, we identified a P450, designated *Cc*CYP9Z273 (Table S01), that converted multiple steroid substrates into distinct profiles of oxidized products (Fig. S03).

Incubation of pregnenolone (**9**) with microsomes containing *Cc*CYP9Z273 yielded five oxidized compounds (**10, 11** and **15** – **17**), with caraumbellogenin (**10**), and 5β-hydroxypregnenolone (**15**) as the major products (Fig. 2A). The structures of **10, 15**, and **16**, the latter representing a previously undescribed compound, were established by NMR spectroscopy (Fig. S04 – S16, Table S02 – S03), whereas **11** and **17** were identified by comparison with authentic standards.^5,18^ These products revealed that the enzyme is capable of hydroxylating pregnenolone at C3, C7, C14, C15, and C21, highlighting the catalytic promiscuity of *Cc*CYP9Z273. Importantly, the assignment of **10** as the C14β-hydroxylated product caraumbellogenin was supported by comparison with NMR data reported for the cardenolide biosynthetic pathway from the plant *P. sepium*, in which the structure and stereochemistry of the product **10** were confirmed by X-ray crystallography.^5^ *Cc*CYP9Z273 also converted isopregnanolone (**13**) into three products (**18** – **20**), all of which were structurally characterized by NMR spectroscopy (Fig. 2B, Fig. S17 – S36, Table S02, S04 – S05). Compounds **18** and **19** are previously reported steroids,^19,20^ whereas **20** was characterized as a new compound. The C5 epimer of **13**, epipregnanolone (**14**), was also turned over by *Cc*CYP9Z273, yielding **21** and **22**, although levels of product formation were low (Fig. 2C). Progesterone (**11**) was converted into **23**, the structure of which was established by NMR spectroscopy (Fig. 2D, Fig. S37 – S39, Table S02).^21^ Beyond pregnane substrates, *Cc*CYP9Z273 also converted digitoxigenin (**12**) into periplogenin (**24)**, as confirmed by comparison with an authentic standard (Fig. 2E). This C5β-hydroxylation introduces a structural feature that is also present in the aglycones of the beetle metabolites periplogenin xyloside (**3**) and bipindogenin xyloside (**5**). None of the gene candidates yielded detectable products from cholesterol (**6**) under the assay conditions. Phylogenetic analysis with functionally characterized insect P450s showed that *Cc*CYP9Z273 grouped within the CYP3 clan, near *lc*CYP9T1 and *lc*CYP9T2, which catalyze monoterpene hydroxylation, and *Am*CYP9Q3, which is involved in insecticide detoxification (Fig. 3A, Fig. S40).^22,23 24^ The nearest ortholog of *Cc*CYP9Z273 (*Cv*CYP9Z273-like, 65 % amino acid identity, Fig. S41) in the non-producer *C. varians* showed no activity with any of these substrates (Fig. S03).

**Fig. 2.**
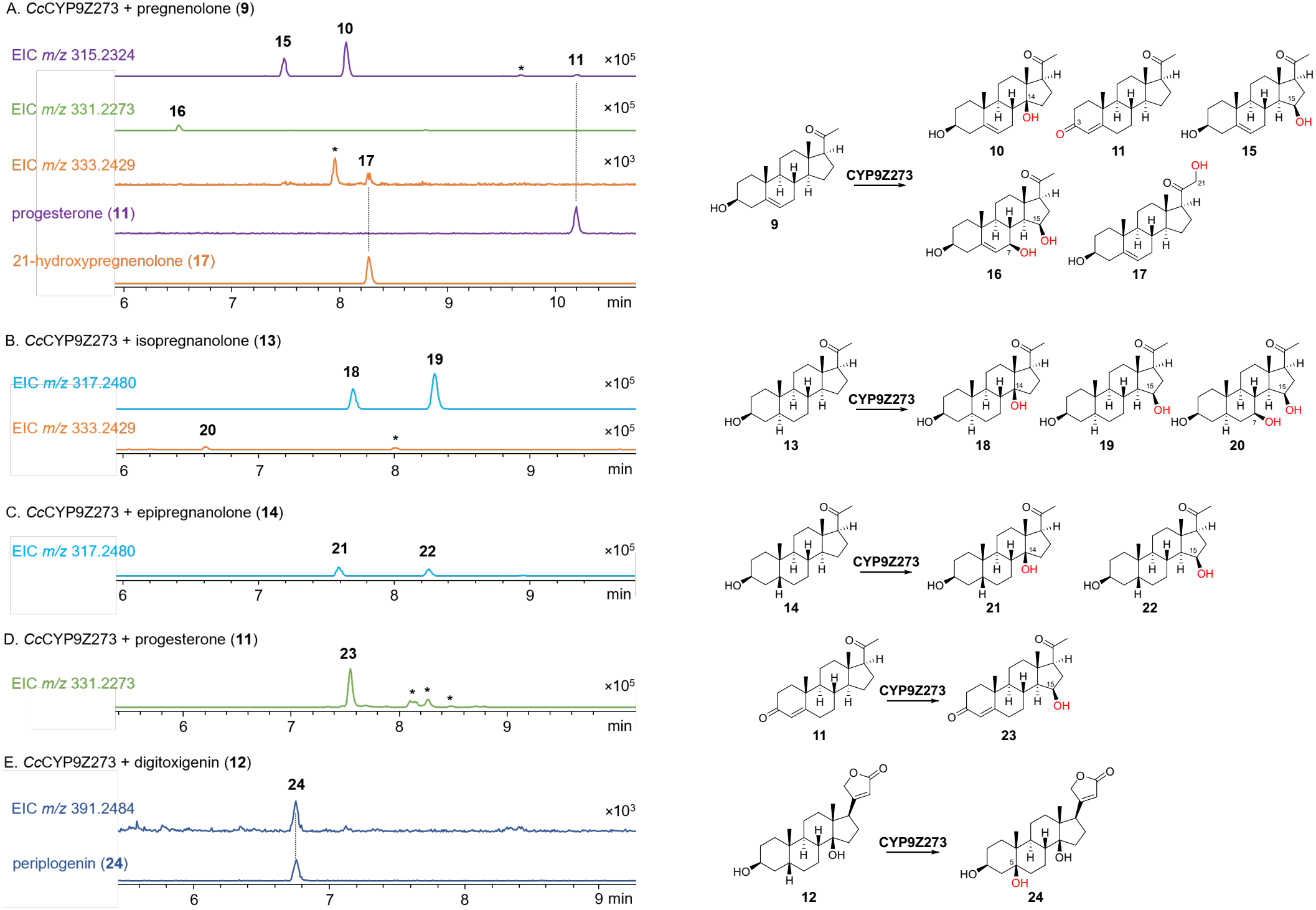
Substrate scope and product profiles of *Cc*CYP9Z273 as assessed with *Saccharomyces cerevisiae* microsomes containing heterologously produced protein. Representative extracted-ion chromatograms (EICs) of products generated by *Cc*CYP9Z273 from **9** and **11** – **14**. Enzyme assays were performed with three biological replicates (n = 3) for each substrate. Representative extracted ion chromatograms (EICs) are shown. Products were assigned as compounds **10, 11**, and **15** – **24**. Corresponding product structures are shown on the right, with sites of oxidation highlighted in red. Asterisks denote additional products detected in the enzymatic reactions but not structurally characterized. EIC intensity scales are indicated on the right. Chromatograms showing products with *m/z* 315 (**10, 11**, and **15**) are shown in purple, *m/z* 331 (**16** and **23**) shown in green, *m/z* 333 shown in orange (**17** and **20**), *m/z* 317 (**18, 19, 21**, and **22**) in light blue, and *m/z* 391 (**24**) shown in dark blue.

**Fig. 3.**
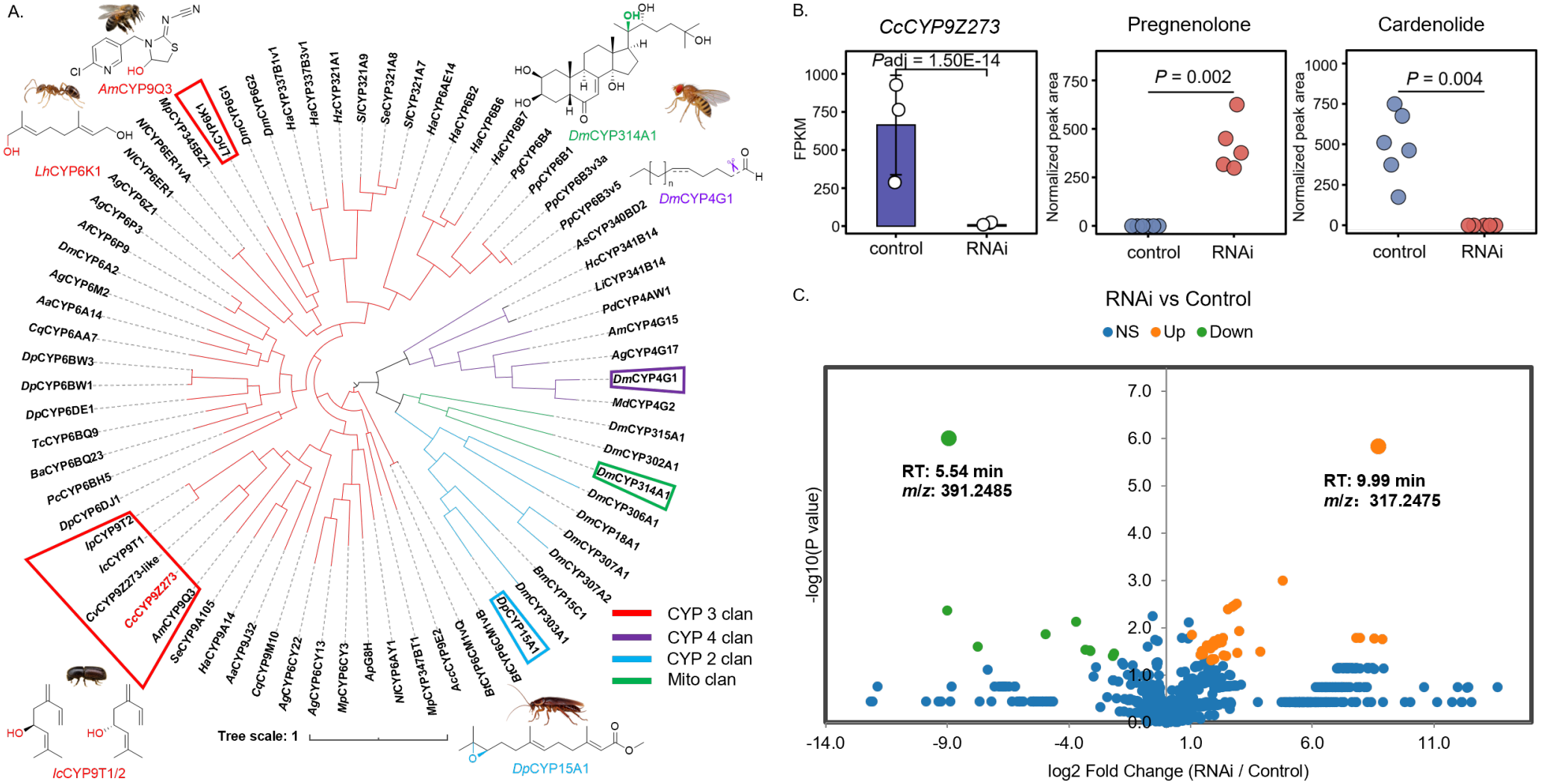
Phylogenetic analysis and gene silencing of *CcCYP9Z273*. (A) Phylogenetic analysis of representative functionally characterized insect cytochrome P450s. Branches are colored according to P450 clan: CYP3 (red), CYP4 (purple), CYP2 (blue), and mitochondrial (green). Representative P450-catalyzed reactions and the corresponding insect species are shown. *Cc*CYP9Z273 is highlighted in red. The tree with bootstrap support values is shown in Fig. S40. (B) *Cc*CYP9Z273 transcript abundance and levels of pregnenolone (**9**) and a cardenolide-associated feature (*m*/*z* 391.2485) in control and RNAi-treated larvae. Transcript abundance is expressed as FPKM, and metabolite levels are normalized LC-MS peak areas. Transcript analysis included three biological replicates per group (n = 3), whereas metabolite analysis included six control samples (n = 6) and five RNAi-treated samples (n = 5). Points represent individual biological replicates, and bars indicate mean ± SD. The adjusted *P* value from the RNA-seq differential expression analysis is shown for transcript abundance; metabolite *P* values were calculated using the Mann-Whitney *U* test. (C) Volcano plot of metabolite features following RNAi. Orange, green, and blue indicate significantly increased, significantly decreased, and non-significantly changed features, respectively. Pregnenolone (**9**, RT 9.99 min, *m*/*z* 317.2475) and the cardenolide-associated feature showing the most pronounced decrease (RT 5.54 min, *m*/*z* 391.2485) are highlighted.

### RNA interference supports a role for *Cc*CYP9Z273 in cardenolide biosynthesis

Since *Cc*CYP9Z273 exhibits relaxed substrate and product specificity in vitro, the role that this enzyme would play in cardenolide biosynthesis was not immediately apparent. Therefore, we performed gene silencing in *C. coerulans* larvae by injecting double-stranded RNA targeting *CcCYP9Z273* to establish whether disruption of the expression of this gene impacted cardenolide levels. Transcriptomic analysis confirmed a significant reduction in its transcript abundance compared with controls (*P* = 0.026; Fig. 3B). Comparative metabolomic profiling revealed changes in the levels of pregnenolone (**9**) and other LC–MS features following *CcCYP9Z273* silencing (Fig. 3B, C, Table S7 – S8). Pregnenolone (**9**; RT 9.99 min, *m/z* 317.2475) exhibited significantly increased levels (*P* = 0.002), consistent with impaired turnover of this substrate, which suggests that **9** is the physiological substrate for *Cc*CYP9Z273. Conversely, the cardenolide-associated feature showing the most pronounced decrease (RT 5.54 min, *m*/*z* 391.2485) was significantly reduced (*P* = 0.004). Additional decreased cardenolide-associated features and other significantly altered LC–MS features are listed in Tables S5 and S6. Together with the biochemical assays, the accumulation of pregnenolone (**9**) and the depletion of cardenolide-associated features support a role for *Cc*CYP9Z273 in pregnenolone hydroxylation during cardenolide biosynthesis.

### Isotope tracing supports caraumbellogenin (10) as a biosynthetic intermediate

The chemotype of the gene silenced larvae clearly suggested that *Cc*CYP9Z273 is involved in cardenolide biosynthesis and that pregnenolone (**9**) is a physiological substrate for this enzyme. However, *Cc*CYP9Z273 can generate multiple oxidized derivatives of **9** when assayed in vitro, and these gene silencing experiments cannot definitively establish which of these enzyme products contributes to the downstream pathway. In vitro, *Cc*CYP9Z273 converted **9** into caraumbellogenin (**10**) and 15β-hydroxypregnenolone (**15**), along with several minor products (**11, 16** and **17**). Consequently, accumulation of **9** following silencing of *Cc*CYP9Z273 is consistent with a biosynthetic hypothesis in which cardenolide biosynthesis proceeds through **10** or **11**, as described in plants, or the alternative major product **15**.

To determine which products of *Cc*CYP9Z273 contribute to cardenolide biosynthesis, we traced the metabolic fate of isotope-labeled precursors in *C. coerulans*. We fed (^13^C_3_)-**6** (cholesterol) and (^13^C_2_, ^2^H_2_)-**9** (pregnenolone) to adults, and (^13^C_5_)-**11** (progesterone) to both adults and larvae. Labeled (^13^C_2_, ^2^H_2_)-**10** (caraumbellogenin) and (^13^C_2_, ^2^H_2_)-**15** (15β-hydroxypregnenolone) were obtained by preparative-scale conversion of labeled (^13^C_2_, ^2^H_2_)-**9** using *Cc*CYP9Z273, followed by isolation and purification. Because only limited amounts were available, these compounds were administered only to larvae by injection (Fig. 4A). LC–MS analysis revealed isotope incorporation into all cardenolides detected in the beetle extracts after feeding with labeled **6** and **9** or injection of labeled **10**. The cardenolide detected at a retention time of 5.90 min and an *m/z* of 537.3063 is shown as a representative example. Feeding labeled cholesterol (**6**) to adults yielded a signal at the expected mass of the triply labeled representative cardenolide (*m/z* 540.3159), with a retention time matching that of the endogenous compound. Feeding labeled pregnenolone (**9**) likewise produced an enriched isotopologue at *m/z* 541.3251, consistent with retention of two ^13^C and two deuterium labels. These results support the incorporation of both **6** and **9** into cardenolides in beetles. In contrast, feeding labeled progesterone (**11**) did not yield detectable incorporation into the cardenolides in either adults or larvae under the conditions tested (Fig. 4B and C). However, administration of labeled caraumbellogenin (**10**) resulted in a detectable signal at the expected mass of the labeled cardenolide (*m/z* 541.3251), with a retention time matching that of the endogenous compound. In contrast, injection of labeled 15β-hydroxypregnenolone (**15**) did not produce any detectable isotopic enrichment of the cardenolides. The products **16** and **17** are produced in trace amounts by *Cc*CYP9Z273 and so we did not attempt to isolate the labeled compounds to test for incorporation. Thus, although **10** and **15** are both major products of pregnenolone conversion in vitro, only **10** was incorporated into the cardenolides (Fig. 4B and C).

**Fig. 4.**
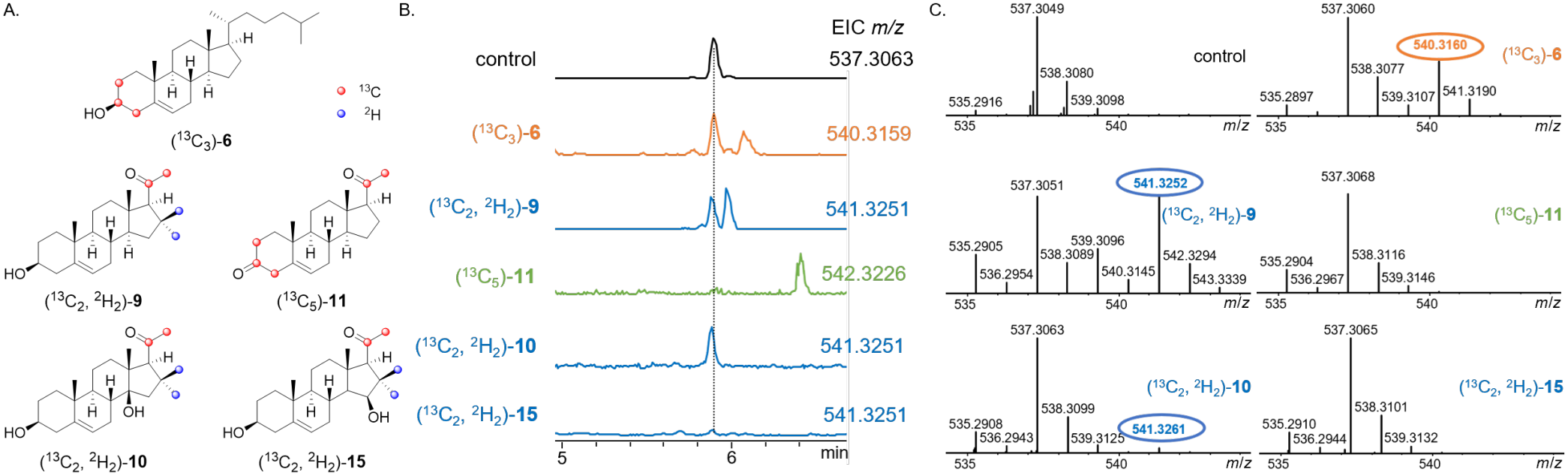
In vivo conversion of isotope-labeled substrates administered to *C. coerulans*. (A) Structures of the isotope-labeled compounds **6, 9**, and **11** were administered by feeding, whereas **10** and **15** were administered by injection. ^13^C and ^2^H labels are indicated in the structures. (B) Extracted ion chromatograms of the corresponding products detected in beetle extracts after feeding or injection of the labeled substrates. The control chromatogram, representing no isotopic label, is shown in black. Products derived from the labeled **9** and **10** exhibited slightly earlier retention times, consistent with the chromatographic isotope effect of deuterium during LC–MS analysis. (C) Mass spectra of the corresponding product peaks used to assess the incorporation of isotope labels. The expected isotope-shifted ions were detected following administration of **6, 9**, and **10**, whereas no corresponding mass shifts were observed for **11** and **15**. Experiments were repeated three times.

### Active-site substitutions alter the regioselectivity of *Cc*CYP9Z273

The broad substrate scope and product profile of *Cc*CYP9Z273 prompted us to investigate the mechanistic basis of this catalytic promiscuity. Using a structural model generated with AlphaFold 3, we selected residues within 3 Å of the heme for site-directed mutagenesis (Fig. 5A). The resulting variants were expressed in yeast and screened using a whole-cell feeding assay with the steroid substrates previously shown to be converted by the wild-type enzyme. Although nearly all mutations abolished or reduced levels of detectable product formation, incubation with the pregnenolone (**9**) substrate revealed several mutants (W126F, V333A, V333G and G337A) that resulted in higher levels of 5β-hydroxypregnenolone (**15**) relative to caraumbellogenin (**10**) when compared to the wild type enzyme (Fig. S42). Assays with proteins isolated as microsomal preparations confirmed this shift in product distribution for W126F, V333A, and V333G mutants, although total levels of product were substantially reduced compared to microsomes containing wild-type protein (Fig. 5B, Fig. S43). Strikingly, assays with G337A showed formation of **15** at levels approximately equivalent to the wild-type protein, while levels of caraumbellogenin (**10**) were reduced to substantially lower levels (Fig. 5B, Fig. S43). G337A therefore appears to serve a switch between these competing reactions. Docking of pregnenolone (**9**) into the wild-type and G337A models using SwissDock suggested that **9** adopted a different orientation relative to the heme in G337A, with C15 positioned closer to the heme iron than in the wild-type protein (Fig. 5C, Fig. S44, Table S09). This predicted difference is consistent with the higher relative levels of C15β-hydroxylation and suggests that the local environment around Gly337 contributes to substrate positioning and regioselectivity.

**Fig. 5.**
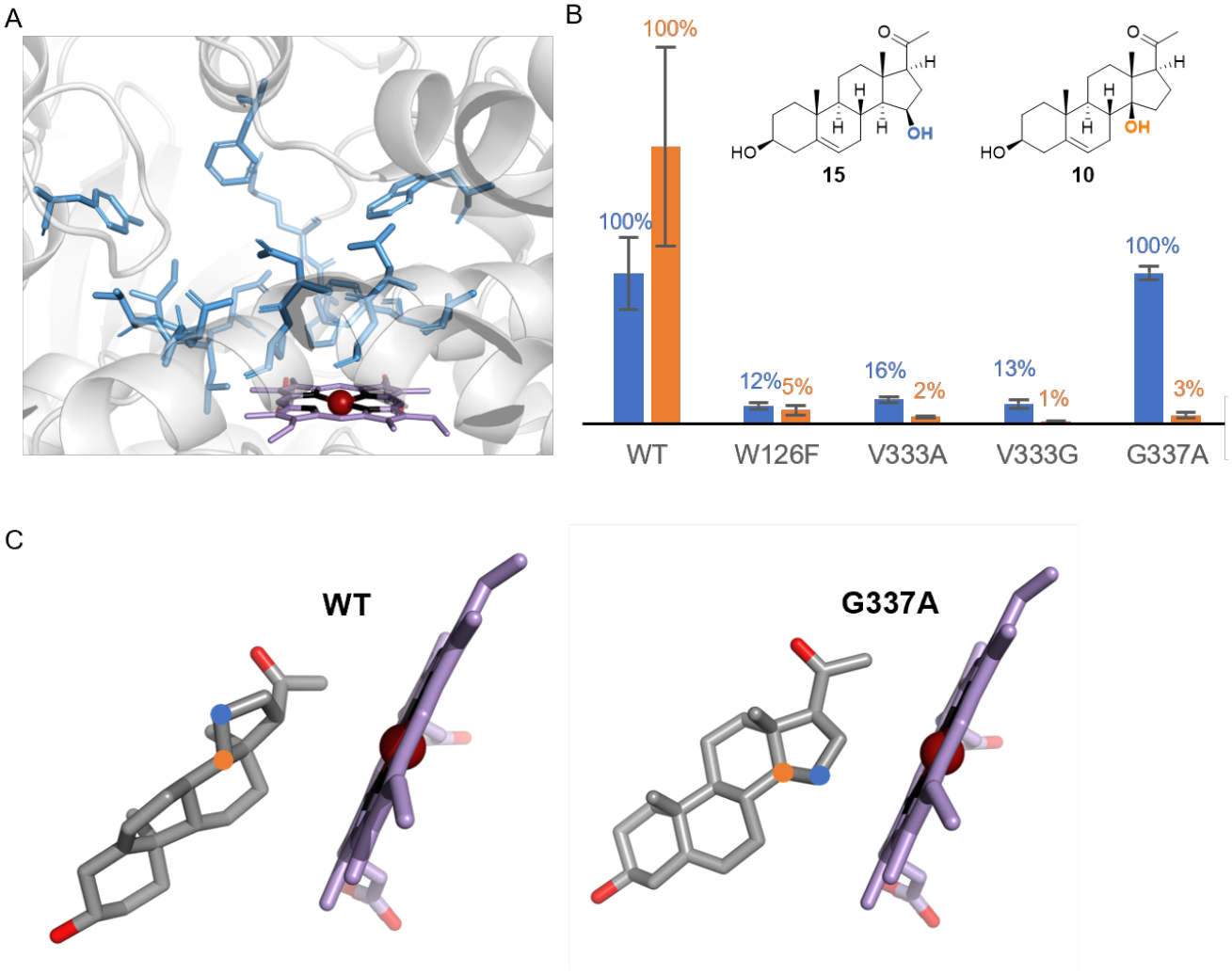
Mutational analysis and molecular docking reveal that G337 impacts the regioselectivity of pregnenolone (**9**) hydroxylation. (A) Predicted substrate-binding pocket of the wild-type (WT) enzyme showing amino acid residues within 3 Å of the heme cofactor. The heme and surrounding residues are shown as sticks. (B) Relative formation of the C15- (blue) and C14-hydroxylated (orange) products by WT and the indicated mutants. Product formation by WT was normalized to 100%. Bars represent mean ± SD (n = 3). Individual data points are shown in Fig. S43. (C) Representative productive-like docking poses of **9** in WT and the G337A mutant. In WT, the distances from C14 (orange) and C15 (blue) to the heme iron are 5.53 and 5.72 Å, respectively, whereas in G337A, the corresponding distances are 5.73 and 4.37 Å, respectively.

**Fig. 6.**
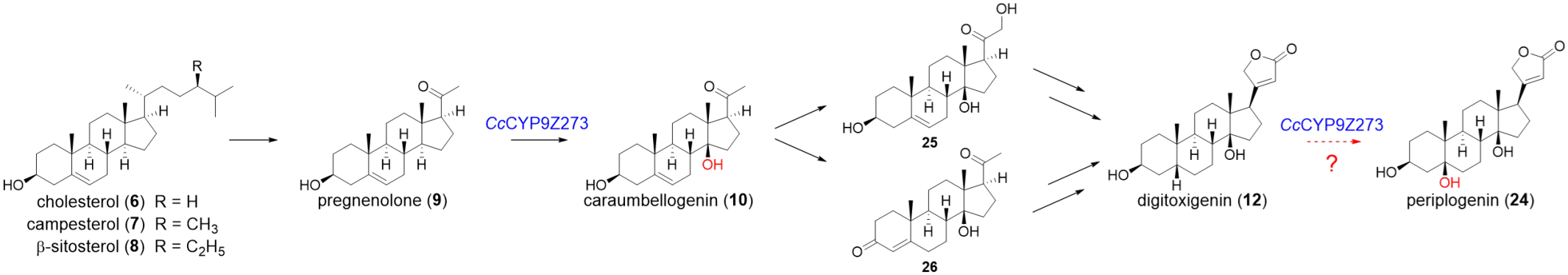
Proposed biosynthetic pathway to periplogenin (**24**) in *Chrysolina* beetles. *Cc*CYP9Z273 catalyzes the C14β- hydroxylation of pregnenolone (**9**) to caraumbellogenin (**10**). Alternative downstream routes through the putative intermediates **25** and **26** that have been proposed to lead to digitoxigenin (**12**) in plants^12^. *Cc*CYP9Z273 also converts **12** to periplogenin (**24**) in vitro, although the contribution of this reaction to biosynthesis in vivo remains unresolved.

## Discussion

Cardenolides are fascinating natural products from chemical, ecological and medicinal perspectives. These compounds are best known as plant-derived natural products, but reports from decades ago have shown that leaf beetles (*Chrysolina*) also produce cardenolides. While the biosynthetic pathways for plant-derived cardenolides have been extensively studied, little is known about the molecular or genetic basis for production of these compounds in insects. Here we identify a cytochrome P450, *Cc*CYP9Z273, that plays an essential role in cardenolide biosynthesis in *C. coerulans*. In vitro biochemical assays with heterologous protein indicate that this enzyme can modify a diverse range of steroid substrates. However, silencing of this gene in *C. coerulans* results in accumulation of pregnenolone (**9**), which strongly suggests that **9** is a physiologically relevant substrate. Although *Cc*CYP9Z273 produces both the C14 and C15 hydroxylated products of pregnenolone (**9**) in vitro, isotope tracing studies indicate that the 14β-hydroxy product, caraumbellogenin (**10**), is the sole on-pathway intermediate to cardenolides (Fig. **6**); incorporation of the alternative major C15-hydroxylated product **15** was not detected under the conditions tested. Additionally, progesterone (**11**), a validated biosynthetic intermediate in some plant cardenolide biosynthetic pathways,^13^ is not incorporated when fed to *C. coerulans*.

Plants from *Digitalis* and *Periploca* genera, like beetles, utilize the conversion of **9** into **10** in the biosynthesis of cardenolides. Notably, *Digitalis* and *Periploca* plants use 2-oxoglutarate-dependent dioxygenases (2OGD) to catalyze this stereoinverted C14β-hydroxylation, while *C. coerulans* employs a cytochrome P450. In the model of *Cc*CYP9Z273, the C14 position of the substrate is oriented relative to the heme in a geometry compatible with abstraction of the C14α-H by the P450 oxo species (Fig. 5C). Formation of the 14β-hydroxylated product could involve stereochemical inversion after hydrogen abstraction, potentially through reorientation of the substrate radical followed by hydroxyl rebound from the opposite face, analogous to the mechanisms proposed for the corresponding 2OGD-catalyzed reactions in *Digitalis* and *Periploca*.^5,12^

Although we identified G337 as a residue important for controlling the distribution between C14 and C15 hydroxylation, it remains unresolved why wild-type *Cc*CYP9Z273 yields two major products from pregnenolone. The C15-hydroxylated product may serve another function in *C. coerulans* metabolism, or perhaps, in the context of the native cellular environment, the product specificity of *Cc*CYP9Z273 is more selective for on-pathway C14 hydroxylation. However, these findings clearly establish that *Cc*CYP9Z273 catalyzes an early, physiologically relevant C14β-hydroxylation step in beetle cardenolide biosynthesis. We additionally note that *Cc*CYP9Z273 can also catalyze C5β-hydroxylation of digitoxigenin (**12**) in vitro, which is a structural feature that present in the aglycones of the beetle metabolites **3** and **5**. This suggests that *Cc*CYP9Z273 could potentially catalyze a second reaction in cardenolide diversification (Fig. **6**), but an additional, downstream catalytic function cannot be established by gene silencing. Nevertheless, despite these remaining ambiguities, identification of *Cc*CYP9Z273 establishes an entry point into investigating how insects assemble defensive cardenolides and the extent to which plant and insect pathways share intermediates while relying on independently evolved catalytic machinery.

## Materials and Methods

### Beetles

Adult *Chrysolina coerulans* and *Chrysolina varians* were collected from gardens and other outdoor sites in and around Jena, Germany. Field-collected *C. coerulans* adults were kept at 22–23 °C, fed commercially purchased organic mint leaves, and allowed to lay eggs. The eggs were kept at the same temperature until they hatched, and the resulting larvae were used in experiments.

### Chemicals

Solvents were purchased from Fisher Scientific. HPLC-grade solvents were used for extraction and semi-preparative HPLC, whereas MS-grade solvents were used for LC–MS analyses. Cholesterol (**6**), b-sitosterol (**8**), 21-hydroxypregnenolone (**17**) and periplogenin (**24**) were purchased from Sigma-Aldrich. Campesterol (**7**) was obtained from PhytoLab. Pregnenolone (**9**) and digitoxigenin (**12**) were purchased from TCI, and progesterone (**11**) was obtained from EMD Millipore. Isopregnanolone (**13**) and epipregnanolone (**14**) were purchased from Steraloids. All isotopically labeled substrates used in this study were purchased from Sigma-Aldrich.

### Beetle dissection and RNA extraction

The head, fat body, elytra, and gut of adult *C. coerulans*, as well as the head, fat body, and remaining body tissues of adult *C. varians*, were dissected in 1 × phosphate-buffered saline (PBS) and immediately transferred to ceramic bead tubes containing ice-cold TRIzol reagent (Direct-zol). Tissues from five specimens per life stage were pooled and homogenized using a TissueLyser LT (Qiagen). Total RNA was extracted using the Direct-zol RNA Miniprep kit (Zymo Research) with DNase treatment. RNA concentration was measured using an N60 nanophotometer (Implen), and RNA integrity was assessed using a TapeStation System (Agilent).

### RNAseq and transcriptome assembly

The total RNA was further processed by the Max Planck Genome Center (Cologne, Germany), performing poly(A)+ mRNA enrichment using oligo-dT magnetic beads and library preparation using the TruSeq Sample Prep approach. Sequencing of one replicate pool per tissue was carried out on an Illumina NextSeq 2000 Genome Analyser platform using paired-end (2 × 150 bp) read technology resulting in 25-30 Mio reads per sample. The obtained reads were trimmed using Trim Galore with a Phred quality score of 30 and a length cutoff of 50 bp. Sequences were further processed by removal of reads containing primer/adaptor sequences and ribosomal RNA using OmicsBox v3.4.1 (https://www.biobam.com/omicsbox). Transcriptome *de novo* assemblies were created using CLC Genomics Workbench v20.1, combining all sequence reads obtained from the different tissue samples and developmental stages of *C. coerulans*. The *de novo* reference transcriptome assembly of *C. coerulans* contained 48,950 contigs (minimum contig size = 250 bp; minimum contig coverage cut-off criteria = 3x) with an N50 contig length of 1,738 bp and a maximum contig length of 22,036 bp. Transcriptome annotation using BLAST, Gene Ontology mapping and InterProScan implemented in OmicsBox (https://www.biobam.com/omicsbox/) was carried out as previously described.^25^ The quality and completeness of the assembly was assessed by BUSCO v.3 with the Insecta BUSCO set from the OrthoDB v9.1 database. This resulted in 90.7% complete, 4.4% fragmented and 3.5% missing BUSCO genes for the *C. coerulans* transcriptome assembly. Digital gene expression analysis was implemented in CLC Genomics Workbench v20.1 by generating BAM files and using sequence counts to estimate expression levels. Mapped read values were normalized as implemented in CLC Genomics Workbench v20.1 and used to calculate log_2_-transformed transcripts per million (TPM+1) values.

### Identification and selection of candidate cytochrome P450 genes

Cytochrome P450 genes were identified from the *C. coerulans* transcriptome based on functional annotations and Gene Ontology (GO) terms. Transcript abundance was assessed using reads per kilobase per million mapped reads (RPKM) values across different adult and larval tissues. P450 genes were ranked according to their expression levels in the adult fat body of *C. coerulans*, and the eight most highly expressed candidates were selected for functional characterization. Their expression profiles were subsequently compared with those of *C. varians*, a closely related species that does not produce cardenolides.

### Molecular Biology

Molecular biology kits were used according to the manufacturers’ instructions. Total RNA was extracted using the Direct-zol RNA Miniprep kit (Zymo Research), and reverse-transcribed into cDNA using iScript reverse transcriptase (Bio-Rad). Genes and gene fragments were amplified with SuperFi DNA polymerase (Invitrogen).

PCR products purified using the Zymoclean Gel DNA Recovery Kit (Zymo Research). Double-stranded RNA (dsRNA) for RNAi was prepared using the MEGAscript T7 kit (Thermo Fischer Scientific). All oligonucleotide primers were synthesized by Sigma-Aldrich. Plasmid construction was performed using the In-Fusion cloning kit (Takara). Plasmid DNA was purified from bacterial cultures using the Wizard Plus SV Minipreps DNA Purification System (Promega).

### Gene cloning and expression in yeast

A putative NADPH-cytochrome P450 reductase (CPR) was identified from the *C. coerulans* transcriptome based on functional annotation. The corresponding coding sequence was amplified from *C. coerulans* cDNA using SuperFi DNA polymerase. Purified PCR products were cloned into the second multiple cloning site (MCS2) of the pESC-LEU vector, pre-linearized with *Bam*HI and *Hind*III, using In-Fusion cloning (Takara). The resulting constructs were introduced into *Escherichia coli* TOP10 competent cells by heat-shock transformation.

Recombinant colonies were selected on LB agar plates containing carbenicillin (100 μg/mL) and cultured in LB liquid medium supplemented with the same antibiotic. Plasmids were isolated using the Wizard Plus SV Minipreps DNA Purification Kit (Promega) and confirmed by Sanger sequencing (Azenta Life Sciences). The P450 candidate genes were then cloned into MCS1 of the reductase-containing pESC-LEU vector, pre-linearized with *Sac*I and *Not*I, using the same workflow described above.

Sequence-verified constructs were transformed into *S. cerevisiae* INVSc1 using the Frozen-EZ Yeast Transformation II Kit (Zymo Research). Transformants were selected on SD-Leu medium (6.7 g/L yeast nitrogen base without amino acids and 2 g/L dropout mix lacking leucine) supplemented with 2% (w/v) glucose and incubated at 30 °C for 2 days. Positive colonies were identified by colony PCR and inoculated into 10 mL of SD-Leu medium containing 2% (w/v) glucose, then cultured overnight at 30 °C with shaking at 220 rpm. The resulting cultures were preserved as glycerol stocks at a final glycerol concentration of 25% (v/v) and stored at −80 °C for subsequent use.

### Site-directed mutagenesis

Site-directed mutagenesis of *Cc*CYP9Z273 was performed by overlap-based PCR amplification followed by In-Fusion cloning. For each mutant, the *Cc*CYP9Z273 coding sequence was amplified as two separate fragments, with the desired mutation introduced at the junction between the fragments using mutagenic primers. The resulting PCR fragments were assembled into the CPR-containing pESC-LEU vector, pre-linearized with *Sac*I and *Not*I, using In-Fusion cloning (Takara). The resulting constructs were transformed into *E. coli* TOP10 competent cells, and the desired mutations were confirmed by Sanger sequencing. Sequence-verified mutant constructs were subsequently transformed into *S. cerevisiae* INVSc1 as described above.

### Isolation of microsomes from yeast

A single colony of transformed *S. cerevisiae* INVSc1 was inoculated into 10 mL of SD-Leu medium containing 2% glucose and cultured overnight at 30 °C with shaking at 200 rpm. A 1-mL aliquot of the overnight culture was transferred to 100 mL of fresh SD-Leu medium containing 2% glucose and cultured for 24 h at 30 °C with shaking at 200 rpm. Cells were collected by centrifugation at 4,000 × *g* for 5 min at room temperature and resuspended in 100 mL of SD-Leu containing 2% galactose. Expression was induced for more than 24 h at 30 °C with shaking at 200 rpm.

Cells were harvested by centrifugation at 4,000 × *g* for 5 min at 4 °C, washed with 30 mL of TEK buffer (50 mM Tris-HCl, pH 7.5, 1 mM EDTA, and 100 mM KCl), and centrifuged again under the same conditions. Cell pellets were processed immediately or stored at −80 °C. Each pellet was resuspended in 2 mL of TES buffer (50 mM Tris-HCl, pH 7.5, 1 mM EDTA, and 600 mM sorbitol), freshly supplemented with 1% (w/v) serum albumin and 0.012% (v/v) β-mercaptoethanol. Glass beads (500 μm in diameter) were added to the surface of the cell suspension, and cells were disrupted by vigorous shaking for five 1-min cycles, with 1-min intervals on ice. The lysate was recovered, and the beads were rinsed repeatedly with 5-mL portions of supplemented TES buffer to obtain a combined lysate volume of approximately 20 mL.

The combined lysate was centrifuged at 7,500 × *g* for 10 min at 4 °C to remove cell debris. The supernatant was transferred to ultracentrifuge tubes and centrifuged at 100,000 × *g* for 90 min at 4 °C. The microsomal pellet was gently rinsed sequentially with 2 mL of TES buffer and 2 mL of TEG buffer (50 mM Tris-HCl, pH 7.5, 1 mM EDTA, and 30% glycerol). The pellet was resuspended in TEG buffer using two successive 250-μL to 1-mL portions, depending on pellet size, and homogenized using a Potter homogenizer. Microsomal suspensions were aliquoted and stored at −20 °C until use.

### Analytical chromatography

Analytical LC–MS was performed using an UltiMate 3000 UHPLC system (Thermo Fisher Scientific) coupled to an Impact II ultra-high-resolution quadrupole time-of-flight mass spectrometer (UHR-Q-ToF; Bruker Daltonics). Chromatographic separation was achieved on a Kinetex XB-C18 column (100 × 2.1 mm, 2.6 μm, 100 Å; Phenomenex). The mobile phases consisted of water containing 0.1% formic acid (A) and acetonitrile (B). The following gradient was applied: 5% B for 1 min, 5–98% B from 1 to 12.25 min, 98% B until 13.9 min, followed by a return to 5% B at 14 min and re-equilibration until 17 min. The flow rate was 0.3 mL min^−1^, and the column temperature was maintained at 35 °C, and the injection volume was 3 µL. Mass spectrometric analysis was performed in positive electrospray ionization (ESI) mode with a capillary voltage of 4,000 V, an end-plate offset of 500 V, a nebulizer pressure of 2.2 bar, and a drying gas flow of 10 L min^−1^ at 250 °C. Mass spectra were acquired over an m/z range of 100–1,200 at a spectral acquisition rate of 12 Hz using data-dependent Auto MS/MS. Auto MS/MS acquisition was performed with a cycle time of 0.5 s and up to five precursor ions per cycle. Active exclusion was disabled, and collision energy was set to 20 eV for singly charged precursor ions at the specified reference masses. Sodium formate was used as the mass calibration reference.

### Enzymatic assays

Enzymatic assays were performed in a total volume of 100 mL containing 30 µL of yeast microsomes (6 mg mL^−1^ final protein concentration), 65 mL of Tris–HCl buffer (10 mM, pH 7.5), 2 mL of NADPH stock solution (50 mM;) and 3 μL of substrate stock solution in MeOH (2 mg mL^−1^). Reactions were incubated at 30 °C for 2 h and quenched by adding 150 μL of methanol. Samples were filtered and analyzed directly by LC–MS.

### Large-scale biotransformation and compound isolation

For large-scale biotransformation of **9, 11** and **13**, *S. cerevisiae* expressing *Cc*CYP9Z273 was cultured in 2 L of medium as described above. Cells were harvested by centrifugation, the supernatant was discarded, and the cell pellet was resuspended in 200 mL of Tris buffer (50 mM, pH 7.5). Substrate (20 mg dissolved in 2 mL of methanol) was added, and the suspension was incubated overnight at 30 °C with shaking. The reaction mixture was extracted with ethyl acetate (200 mL x 3). The combined organic phases were concentrated under reduced pressure, and the crude extract was fractionated by silica gel column chromatography using ethyl acetate/hexane (2:1 to 1:1, v/v). Product-containing fractions were further purified by HPLC using the system and column described below.

Compound purification was performed using an Agilent 1260 Infinity HPLC system (Agilent Technologies) equipped with an autosampler, quaternary pump, column oven, diode array detector (DAD), and fraction collector. Separations were carried out on a Phenomenex Kinetex XB-C18 column (4.6 × 100 mm, 5 μm, 100 Å) maintained at 35 °C. The mobile phases consisted of water containing 0.1% formic acid (A) and acetonitrile (B). The following gradient was applied: 5% B for 1 min, a linear increase from 5% to 95% B over 12 min, an increase to 100% B within 0.1 min, followed by 100% B for 1.8 min. The mobile phase was then returned to 5% B within 0.1 min, and the column was re-equilibrated at 5% B for 3 min, resulting in a total run time of 18 min. The flow rate was 1.0 mL min^−1^. UV detection was performed at 195, 214, 254, 260, and 360 nm.

### NMR measurements

NMR measurements were carried out on a 700 MHz Bruker Avance III HD spectrometer (Bruker Biospin GmbH, Rheinstetten, Germany) and a 500 MHz Bruker Avance III HD spectrometer (Bruker Biospin GmbH, Rheinstetten, Germany) using standard pulse sequences as implemented in Bruker Topspin ver. 3.6.1. 700 MHz and 500 MHz NMR are equipped with a TCI cryoprobe. Chemical shifts were referenced to the residual solvent signals of CDCl_3_ (*δ*_H_ 7.26/*δ*_C_ 77.16), MeOH-*d*_*3*_ (*δ*_H_ 3.31/*δ*_C_ 49.0), DMSO-*d*_6_ (*δ*_H_ 2.50/*δ*_C_ 39.52). All spectra were recorded at 298 K.

### Phylogenetic Analysis

Amino acid sequences were aligned using ClustalW in MEGA12.^26,27^ A maximum-likelihood phylogenetic tree was inferred using the Poisson substitution model with uniform rates across sites. All alignment positions were retained, and tree searching was performed using nearest-neighbor interchange (NNI). Branch support was evaluated using 1,000 standard bootstrap replicates. The tree was visualized layout using iTOL.^28^

### Isotope-labeled precursor feeding and injection experiments

For feeding experiments, commercially available isotope-labeled cholesterol (**6**), pregnenolone (**9**), and progesterone (**11**) were dissolved individually in ethanol at 5 mg mL^−1^. Each stock solution was diluted 100-fold with water to a final concentration of 50 mg mL^−1^ and applied to mint leaves supplied to *C. coerulans*. Adults were used for experiments with **6** and **9**, whereas both adults and larvae were tested with **11**. Control beetles received mint leaves treated with water. After 5 days, beetles were collected and extracted with 200 mL of methanol for LC– MS analysis.

Because only limited amounts of labeled caraumbellogenin (**10**) and 15β-hydroxypregnenolone (**15**) were available, these compounds were administered to larvae by injection. Labeled **10** and **15** were generated by *Cc*CYP9Z273-catalyzed conversion of labeled **9** and purified before use. Each compound was dissolved in DMSO at 50 mM, and 10 nL was injected per larva, corresponding to 0.5 nmol per individual. Control larvae received 10 nL of DMSO. Larvae were collected 48 h after injection and extracted with 200 mL of methanol for LC–MS analysis. Each treatment included three to five biological replicates.

### RNAi-mediated gene silencing

Double-stranded RNA (dsRNA) targeting *CcCYP9Z273* was prepared using the MEGAscript T7 kit (Thermo Fischer Scientific) according to the manufacturer’s protocol. Potential off-target effects were predicted by searching all possible 21 mers derived from each strand of dsRNA against the transcriptomes of *C. coerulans* and a 350 bp template for dsRNA was generated by PCR using gene-specific primers containing the T7 RNA polymerase promoter sequence. The resulting dsRNA was diluted to a final concentration of 2 μg/μL. Larvae were injected with dsRNA (500 nL) and water was injected as a control (500 nL). All injections were performed using a Nanoject III™ (Drummond Scientific Company). To confirm gene silencing, individuals from control and dsRNA-treated groups were collected 48 h post-injection and subjected to RNA sequencing. For metabolite analysis, samples were collected 5 days post-injection.

### Structural modelling and molecular docking

Structural models of *Cc*CYP9Z273 WT and the G337A variant were generated independently using AlphaFold3, with heme included in each model. Pregnenolone was docked into the active-site cavity using SwissDock with the Attracting Cavities option; all other docking parameters were left at the default settings. Docking outputs were converted into multistate PDB files and inspected in PyMOL. The positions of C14 and C15 of pregnenolone relative to the heme iron were evaluated for candidate binding modes. Docking score was not used as the sole criterion for selecting representative poses because highly ranked poses could adopt geometries in which the candidate oxidation sites were too distant from the heme center to be catalytically plausible. Representative productive-like poses were therefore selected by considering substrate-heme geometry together with cluster representation, docking score and steric plausibility.

The docking analysis was performed to investigate the structural basis for the altered regioselectivity observed for G337A. Static docking does not explicitly model the reactive Compound I state or protein dynamics; the selected structures are therefore interpreted as catalytically plausible binding modes rather than direct representations of the reactive state or transition state.

### Comparison of pregnenolone (9) binding geometries in WT and G337A

Docking of pregnenolone to WT *Cc*CYP9Z273 generated 239 poses. The top-ranked WT solution, pose1 (cluster 0, member 1; MEMBER_SCORE = -2.3943; SP-dG = -8.0817 kcal mol^-1), positioned C14 and C15 9.14 and 10.53 Å from the heme iron, respectively, indicating a non-productive geometry for direct oxidation at either site. A representative productive-like pose, pose190 (cluster 37, member 1), positioned C14 and C15 at 5.53 and 5.72 Å from the heme iron, respectively, placing the two candidate oxidation sites at similar distances from the catalytic center.

Docking of pregnenolone to G337A generated 186 poses. The top-ranked G337A solution, pose1 (cluster 0, member 1; MEMBER_SCORE = -3.7134; SP-dG = -8.0040 kcal mol^-1), positioned C14 and C15 10.71 and 11.54 Å from the heme iron, respectively, and was likewise considered non-productive. In contrast, pose9 (cluster 1, member 1; MEMBER_SCORE = -1.1620; SP-dG = -7.7934 kcal mol^-1) positioned C14 and C15 5.73 and 4.37 Å from the heme iron, respectively, revealing a stronger geometric bias toward C15 than observed in the representative WT pose.

The comparison suggests that substitution of Gly337 by Ala can alter substrate positioning within the active site, favoring a binding geometry in which C15 is brought closer to the heme center while C14 is comparatively displaced. This provides a structural rationale consistent with the experimentally observed strong reduction of C14 oxidation while C15 oxidation is retained in G337A. The docking analysis is interpreted as supportive rather than definitive evidence for the origin of the altered regioselectivity.

## Acknowledgements

We thank Prof. David R. Nelson (University of Tennessee Health Science Center, USA) for assigning the P450 name CYP9Z273. We thank Dr. Maritta Kunert and Sarah Heinicke for assistance with mass spectrometry data acquisition, Dr. Maritta Kunert, Dr. Rayko Halitschke and Anja David for collecting beetles, and Dr. Danny Kessler for providing beetle photograph.

## Author Contributions

S. E. O, T. G. K., and H. X. designed the project and the experiments; H. X., A. S., M. F., K. L., and M. L. F. performed the experiments; H. X., A. S., M. F., S. W., H. V., Y. N., and M. K. analyzed the data; H. X. and S. E. O wrote the manuscript.

## Competing Interest Statement

The authors declare no competing interest.

## Additional information

Supplementary information is available for this paper at XXX

## Notes

### Competing Interest Statement

The authors have declared no competing interest.

### Funding

- Max Planck Society

