## Supporting Figures for "A cytochrome P450 is essential for cardenolide biosynthesis in *Chrysolina* beetles"

### Table of Contents

**Fig. S01**. Cardenolide profiles in adult body parts and larvae of *C. coerulans*. 3
**Fig. S02**. Tissue-specific expression profiles of the remaining P450 candidates. 4
**Fig. S03**. Functional screening of candidate P450s for steroid-modifying activity. 5
**Fig. S04 – Fig. S07**. NMR spectra of compound **10**. 6 – 9
**Fig. S08 – Fig. S09**. NMR spectra of compound **15**. 10 – 11
**Fig. S10**. Structure elucidation of 7β,15β-dihydroxypregnenolone (**16**). 12
**Fig. S11 – Fig. S16**. NMR spectra of compound **16**. 13 – 18
**Fig. S17**. Structure elucidation of 14β-hydroxyisopregnanolone (**18**). 19
**Fig. S18 – Fig. S25**. NMR spectra of compound **18**. 20 – 27
**Fig. S26 – Fig. S29**. NMR spectra of compound **19**. 28 – 31
**Fig. S30**. Structure elucidation of 7β,15β-dihydroxyisopregnanolone (**20**). 32
**Fig. S31 – Fig. S36**. NMR spectra of compound **20**. 33 – 38
**Fig. S37 – Fig. S39**. NMR spectra of compound **23**. 39 – 41
**Fig. S40**. Phylogenetic relationships of *Cc*CYP9Z273 and representative insect P450s. 42
**Fig. S41**. Pairwise amino acid sequence alignment of *Cc*CYP9Z273 and its closest P450 homolog from *Chrysolina varians*. 43
**Fig. S42**. Screening of *Cc*CYP9Z273 variants in yeast fed pregnenolone (**9**). 44
**Fig. S43**. Individual replicate data for pregnenolone (**9**) hydroxylation by *Cc*CYP9Z273 and its mutants. 45
**Fig. S44**. Detailed comparison of representative productive-like docking poses of pregnenolone in WT *Cc*CYP9Z273 and the G337A variant. 46
**Table S01**. Nucleotide sequences of *Cc*CYP9Z273 and the P450 reductase 47
**Table S02**. Isolation yields and NMR spectroscopic data of compounds **10**, **15**, **16**, **18**, **19**, **20** and **23**. 48
**Table S03**. NMR data of 7β,15β-dihydroxypregnenolone (**16**) in MeOH-$d_3$ recorded at 298 K. 49
**Table S04**. NMR data of 14β-hydroxyisopregnanolone (**18**) in MeOH-$d_3$ recorded at 298 K. 50
**Table S05**. NMR data of 7β,15β-dihydroxypregnenolone (**20**) in MeOH-$d_3$ recorded at 298 K. 51
**Table S06**. Nucleotide sequences used for RNAi-mediated silencing of *Cc*CYP9Z273 52
**Table S07**. LC–MS features increased following *Cc*CYP9Z273 RNAi. 53
**Table S08**. LC–MS features decreased following *Cc*CYP9Z273 RNAi. 54
**Table S09**. Selected docking poses of pregnenolone in WT and G337A *Cc*CYP9Z273 55

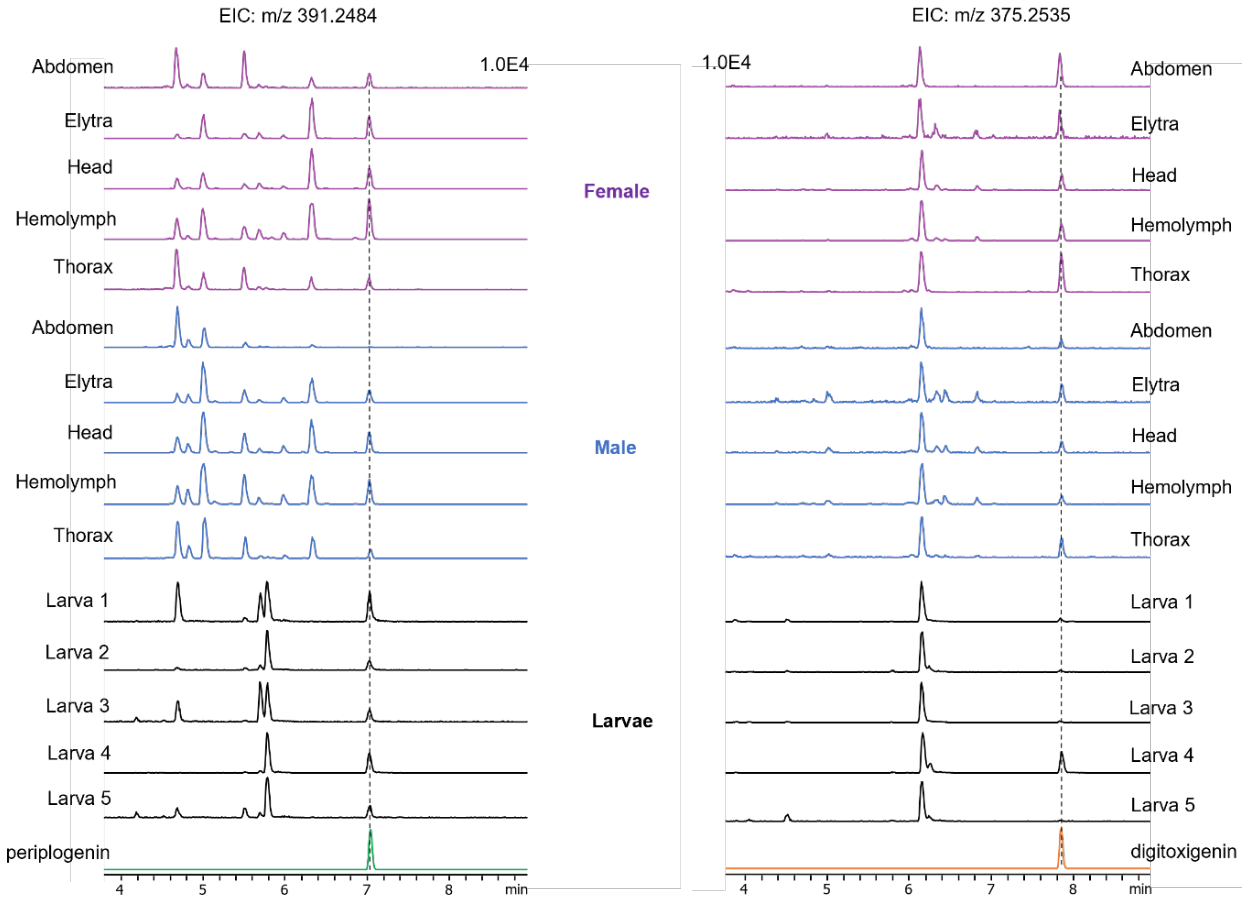


**Fig. S01**. Cardenolide profiles in adult body parts and larvae of *C. coerulans*. Extracted ion chromatograms (EICs) at *m*/*z* 391.2484 (left) and 375.2535 (right) of extracts from the abdomen, elytra, head, hemolymph and thorax of adult females (purple) and males (blue), and five individual larvae (black). Representative chromatograms are shown from five biological replicates (n = 5) for each sample type. The monitored ions include protonated free aglycones and aglycone fragment ions derived from cardiac glycosides.

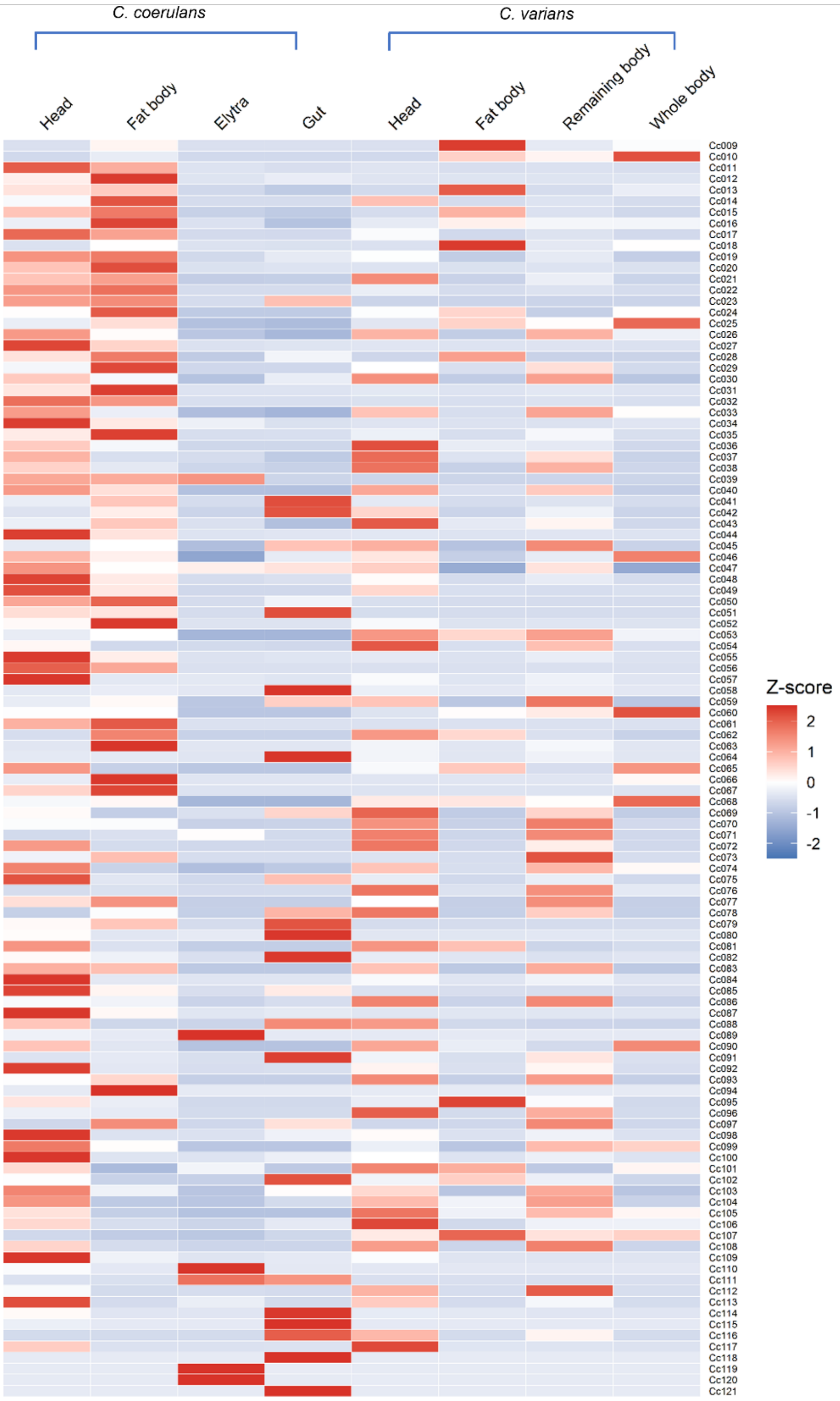


**Fig. S02**. Tissue-specific expression profiles of the remaining P450 candidates. Heatmap showing the relative expression patterns of P450 candidates not included among the eight candidates highlighted in the main text. Candidate selection and expression assessment were primarily based on *C. coerulans*, with *C. varians* samples included for comparison. FPKM values were standardized within each candidate across samples and are displayed as row Z-scores. Red and blue indicate relatively high and low expression, respectively.

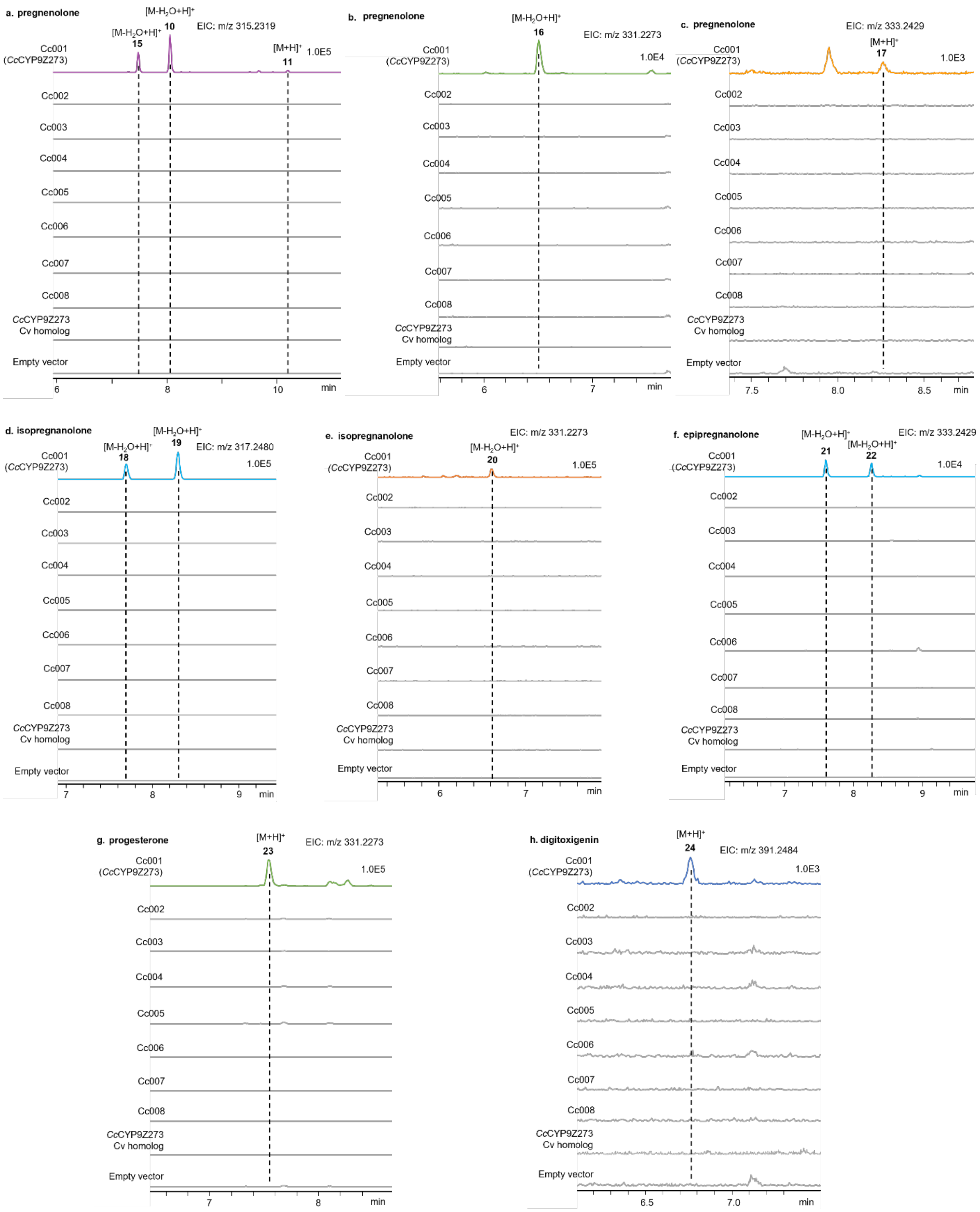


**Fig. S03**. Functional screening of candidate P450s for steroid-modifying activity. Extracted ion chromatograms of reaction mixtures containing yeast microsomes expressing individual *C coerulans* P450 candidates, *Cc*CYP9Z273 homolog from *C varians* and pregnenolone (a–c), isopregnanolone (d,e), epipregnanolone (f), progesterone (g) or digitoxigenin (h). Microsomes from yeast expressing CPR alone served as the negative control. Representative extracted ion chromatograms are shown from three technical replicate enzyme assays (n = 3).

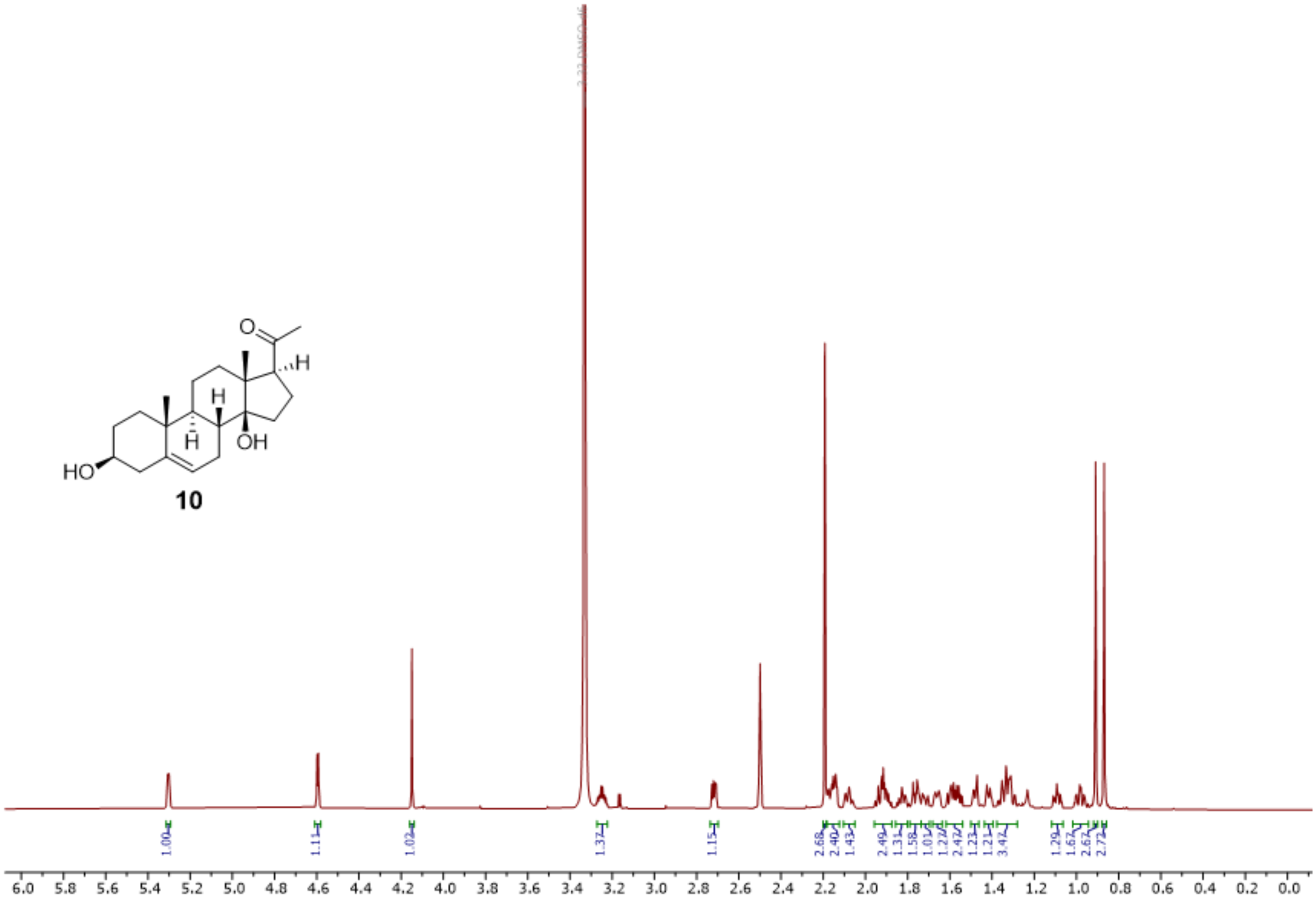


**Fig. S04**. $^{1}$H-NMR spectrum of **10** (700 MHz, DMSO-$d_6$).

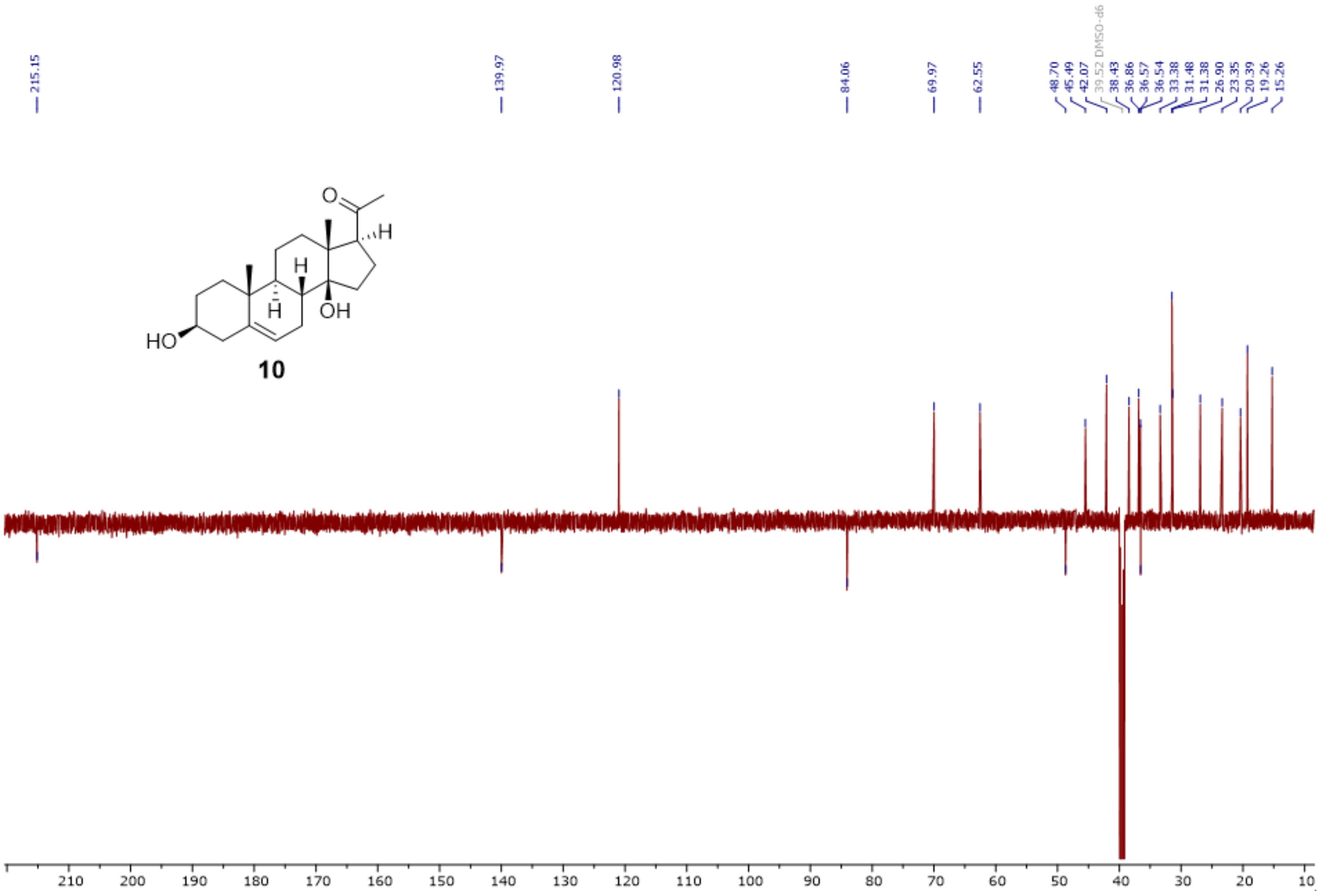


**Fig. S05**. $^{13}$C-DEPTQ spectrum of **10** (176 MHz, DMSO-$d_6$).

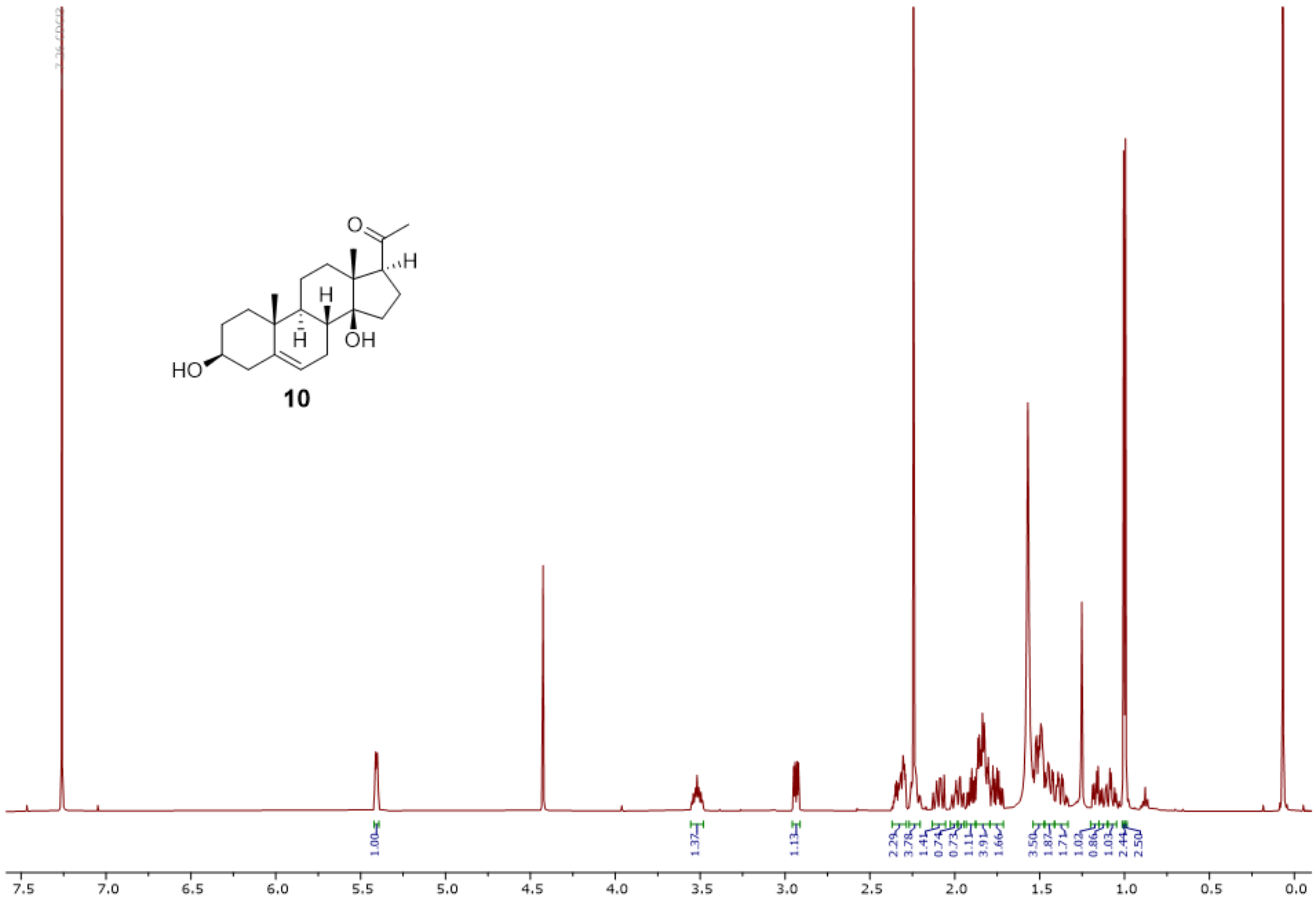

**Fig. S06**. $^1$H-NMR spectrum of **10** (500 MHz, $CDCl_3$).

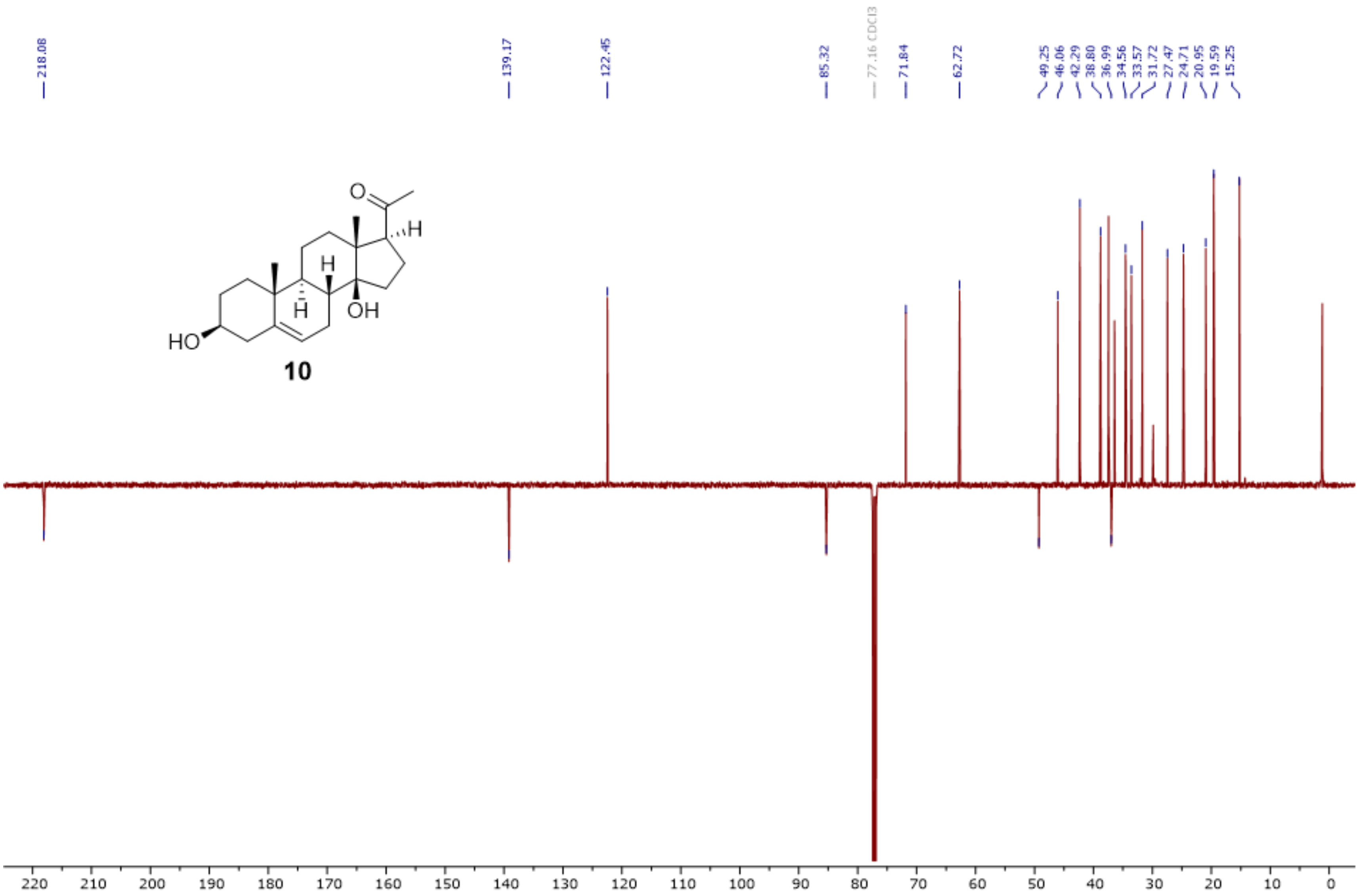


**Fig. S07**. $^{13}$C-DEPTQ spectrum of **10** (126 MHz, $CDCl_3$).

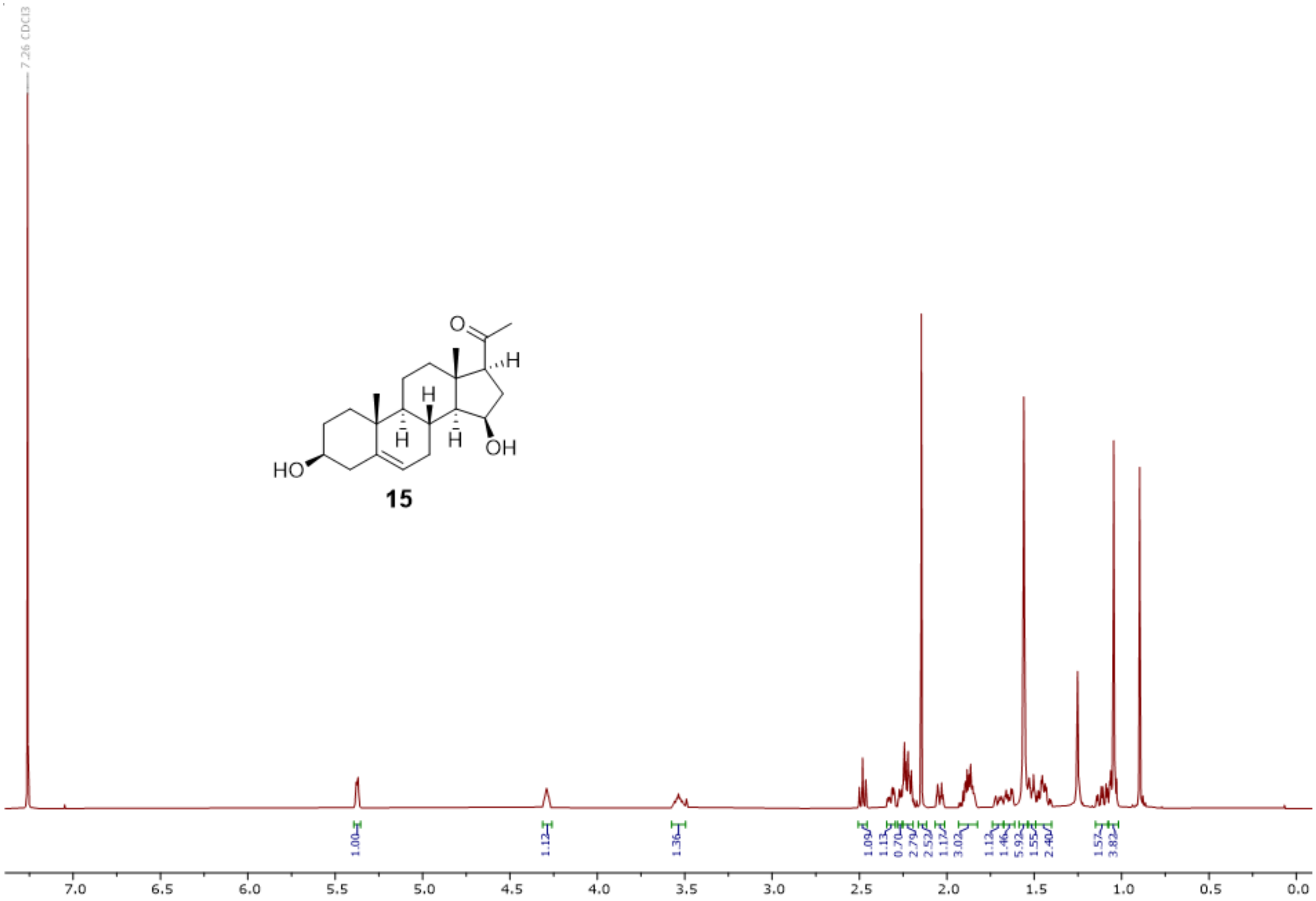


**Fig. S08** $^{1}H$-NMR spectrum of **15** (500 MHz, $CDCl_3$).

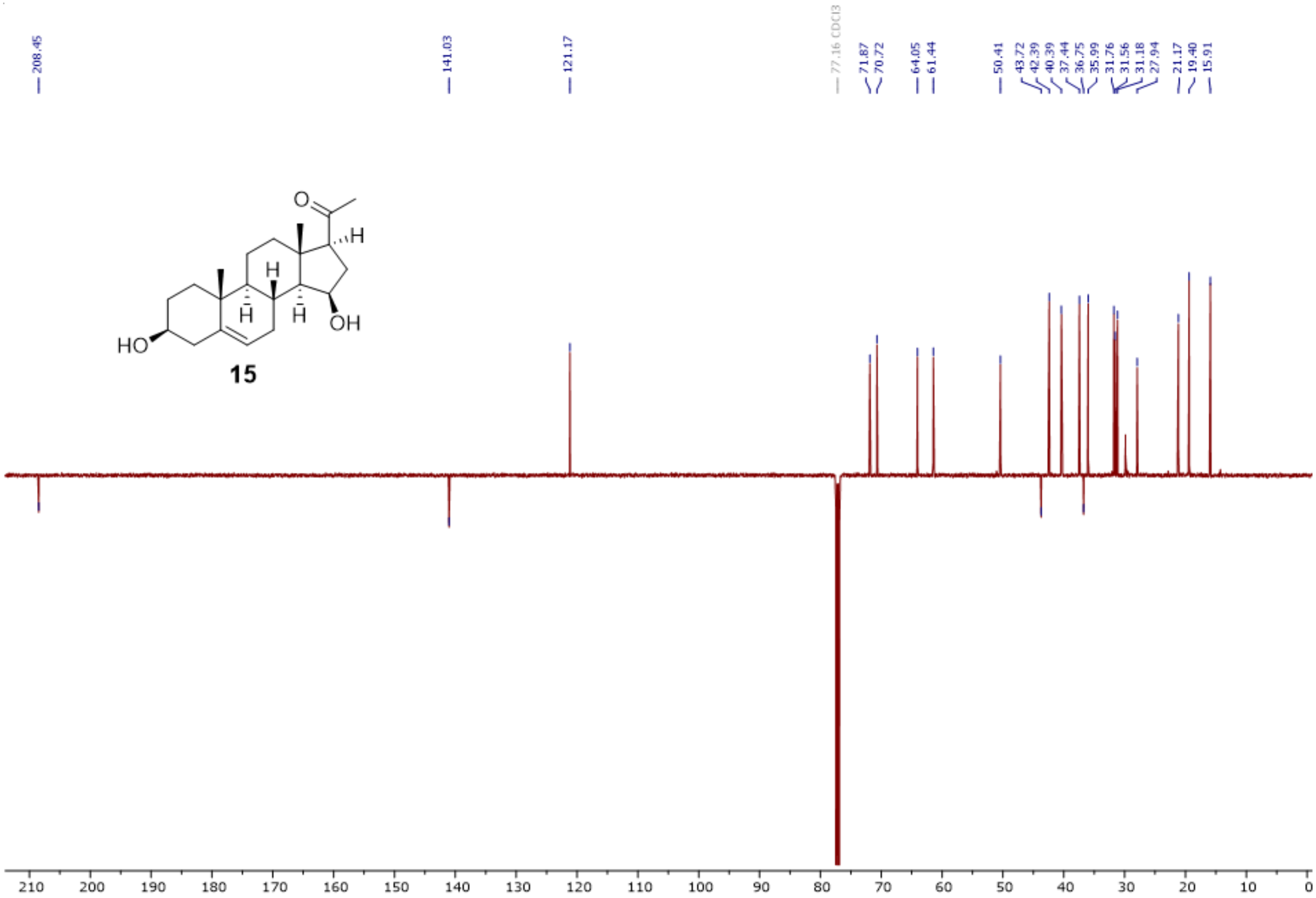


**Fig. S09**. $^{13}$C-DEPTQ spectrum of **15** (126 MHz, $CDCl_3$).

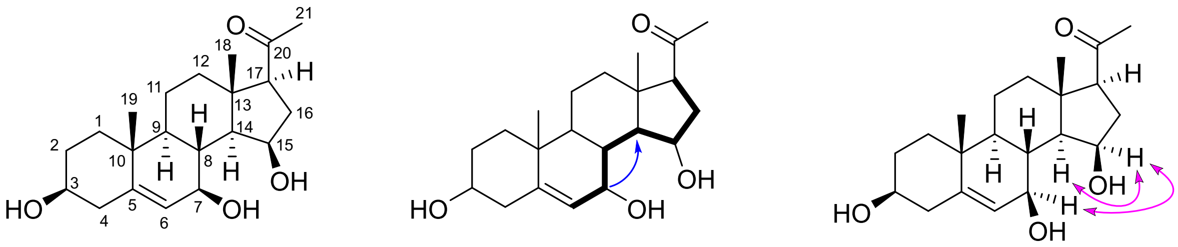

**16**

**Fig. S10**. Structure elucidation of 7β,15β-dihydroxypregnenolone (**16**). Bold: key 1H,1H-COSY, blue single-headed arrows: key HMBC, and pink double-headed arrows: key NOESY correlations.

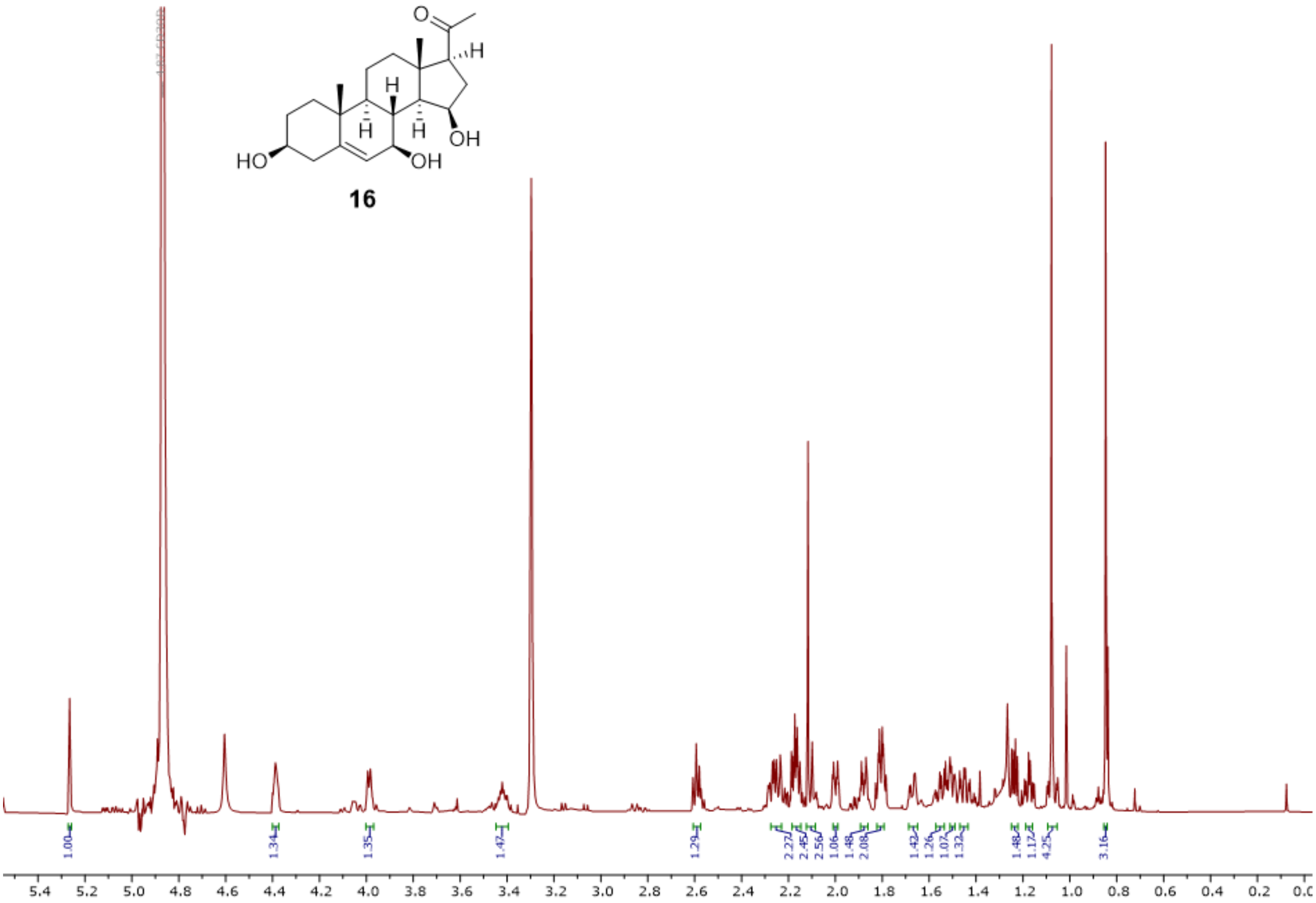


**Fig. S11**. $^{1}$H-NMR spectrum of **16** (700 MHz, MeOH-$d_3$).

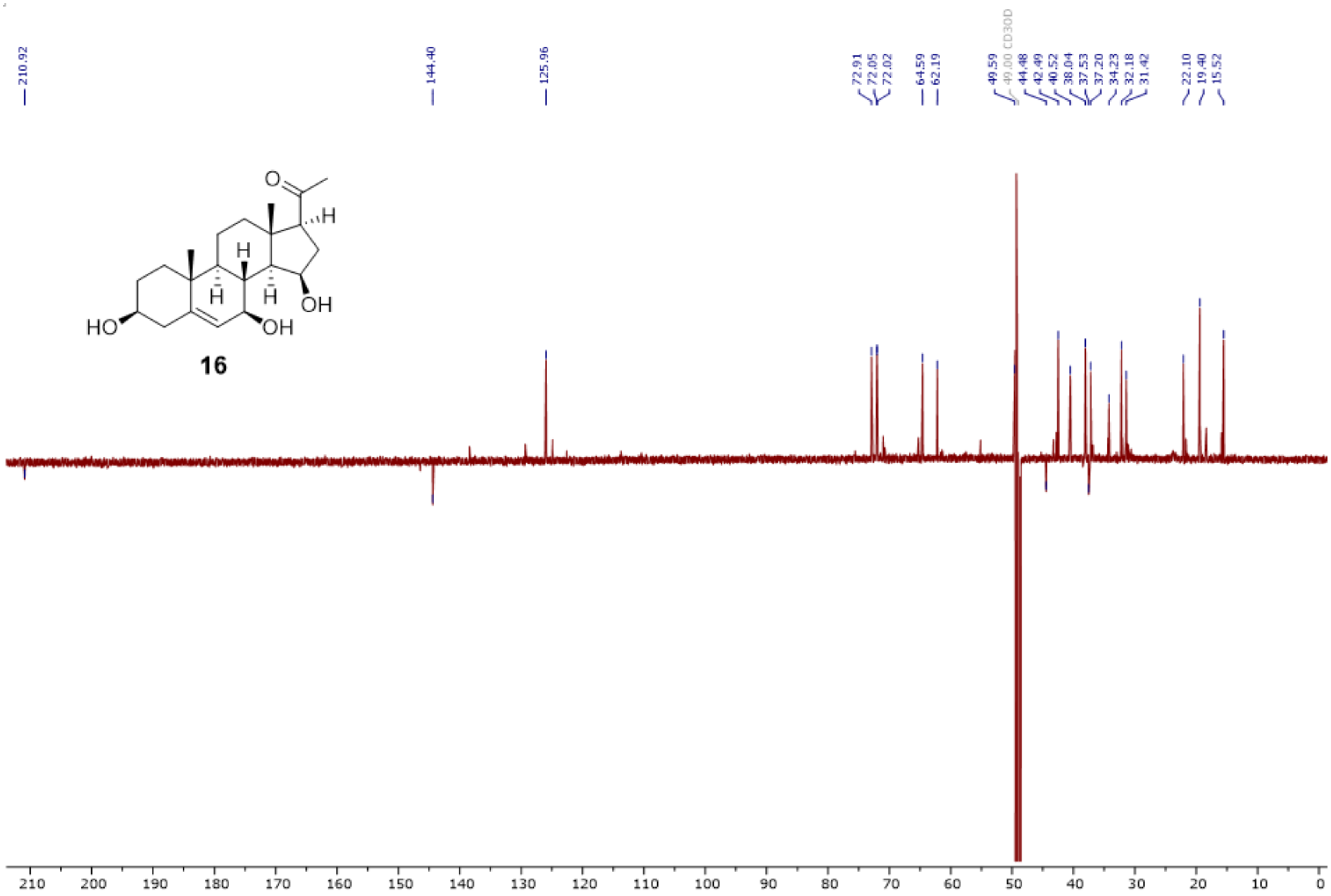


**Fig. S12**. $^{13}$C-DEPTQ spectrum of **16** (176 MHz, MeOH-$d_3$).

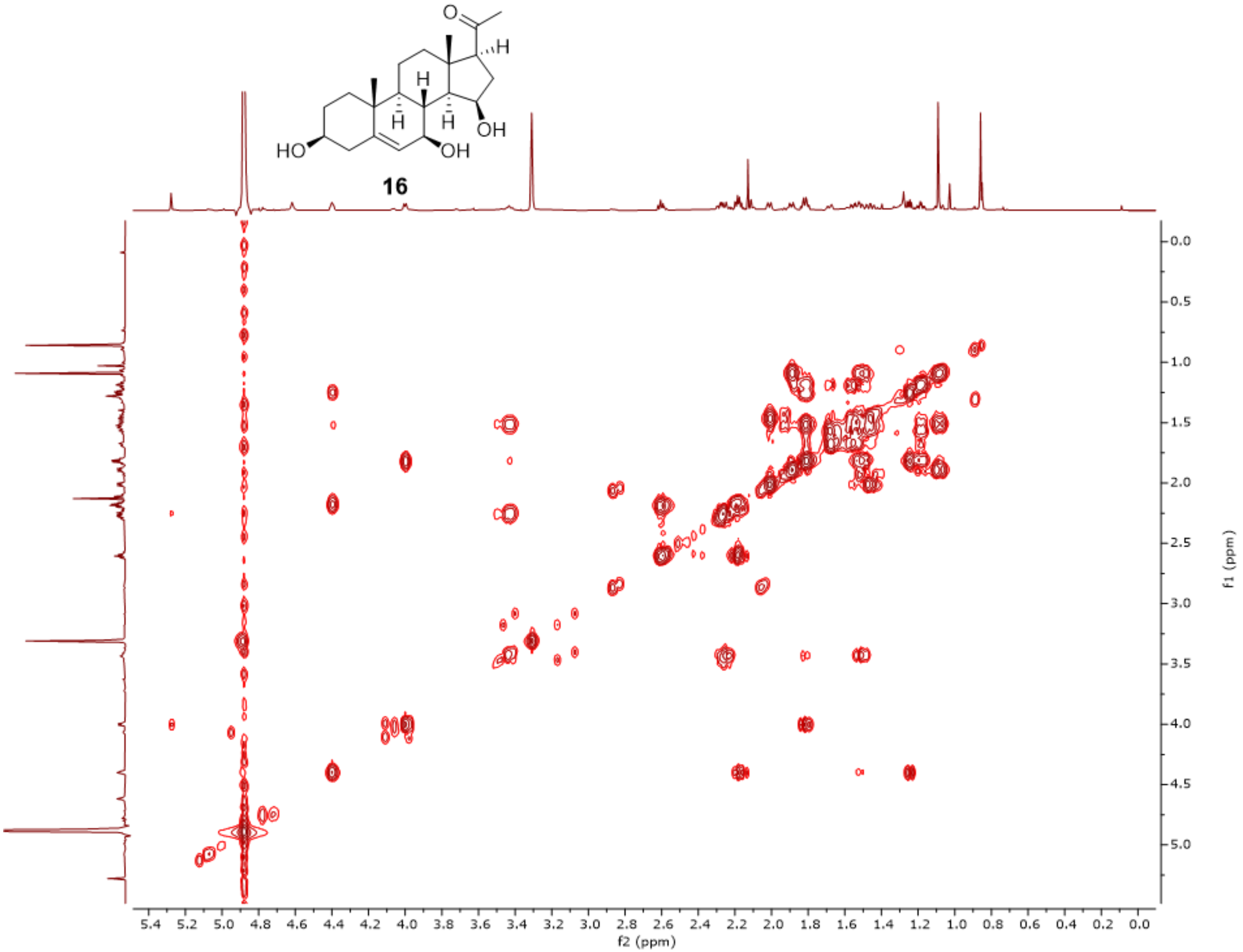


**Fig. S13**. $^{1}H$-$^{1}H$-COSY spectrum (MeOH-$d_3$) of **16**.

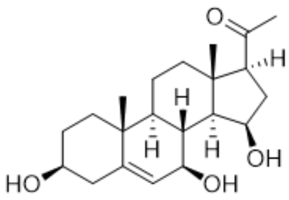


**16**

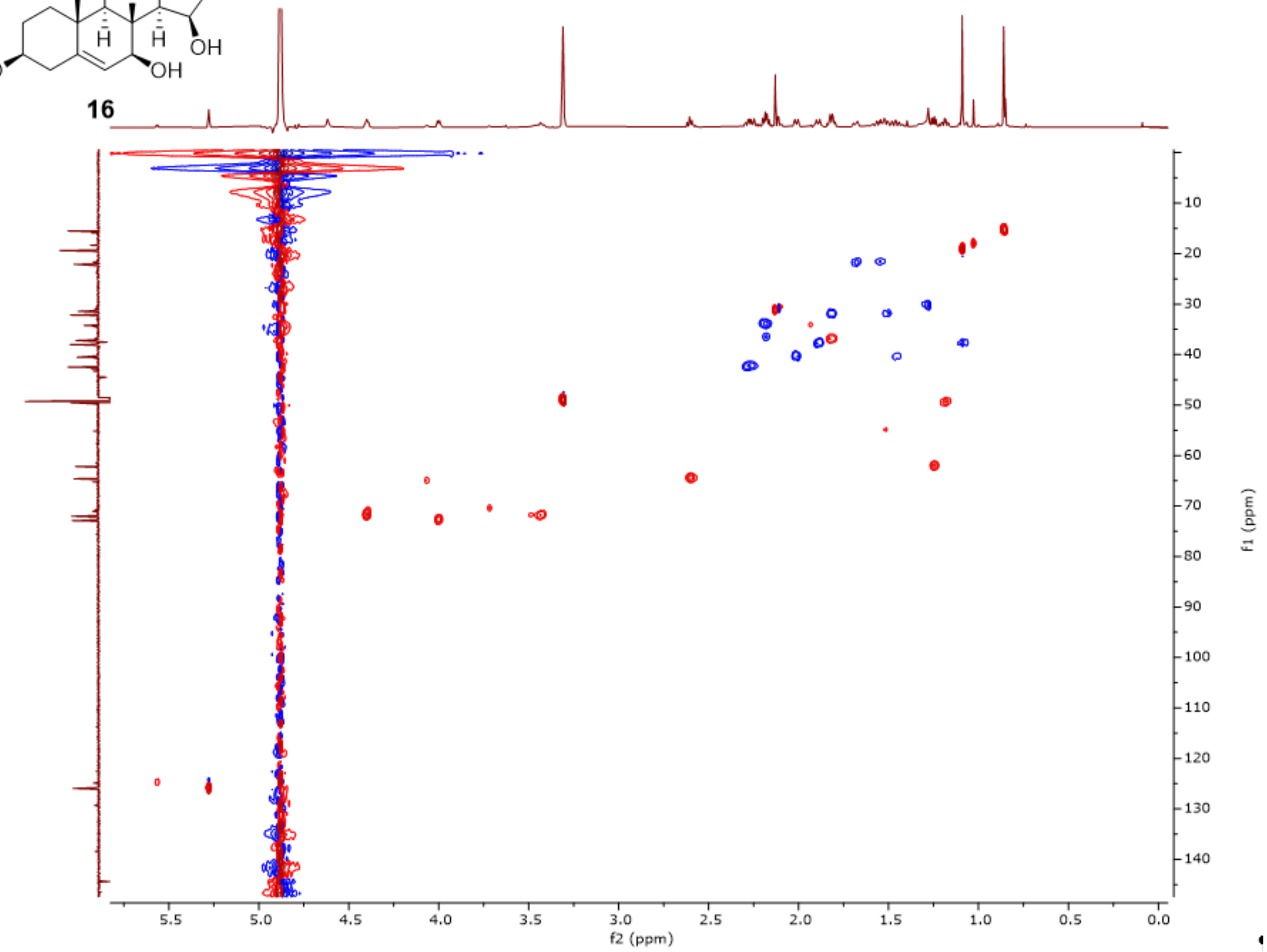


**Fig. S14**. HSQC spectrum (MeOH-$d_3$) of **16**.

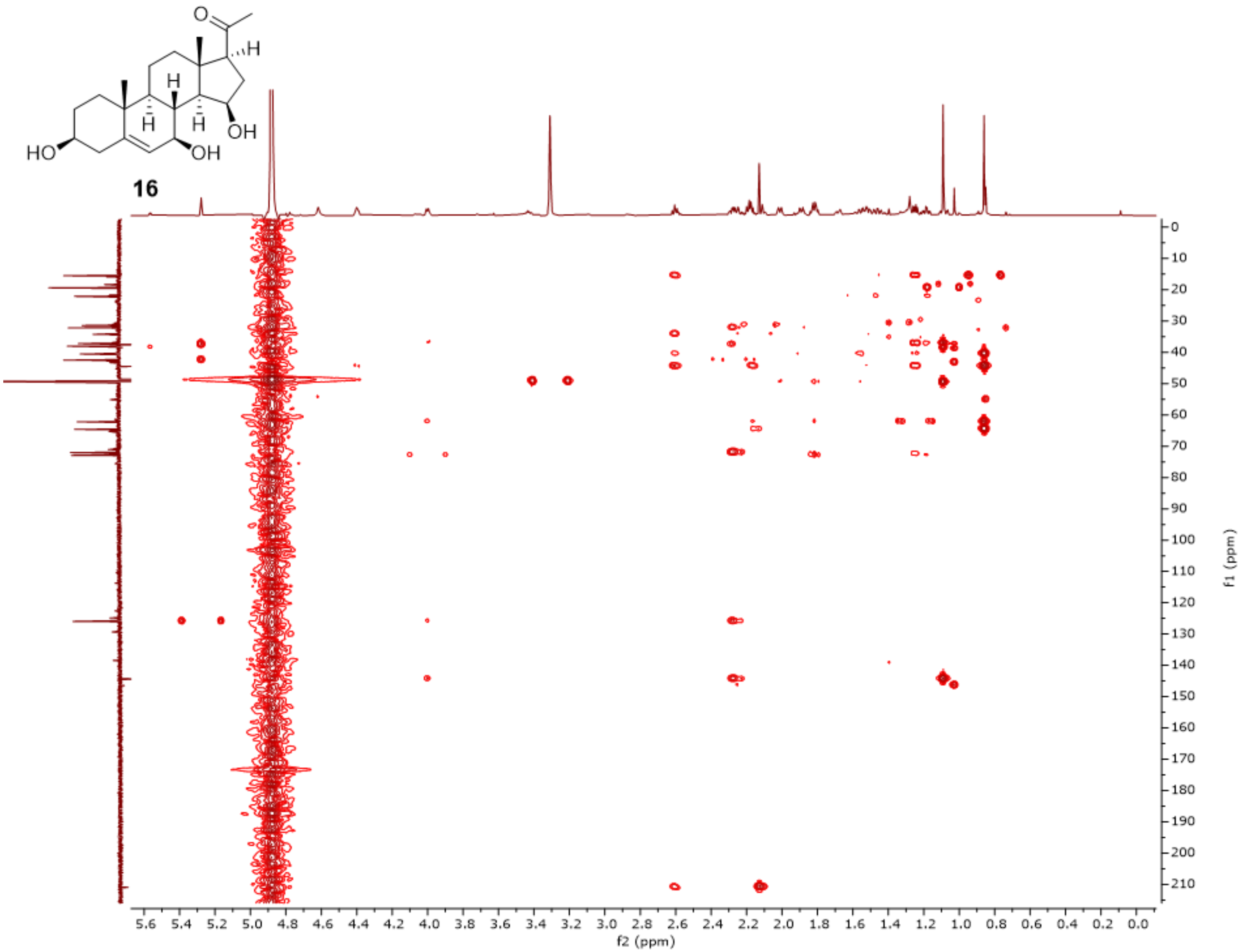


**Fig. S15**. HMBC spectrum (MeOH-$d_3$) of **16**.

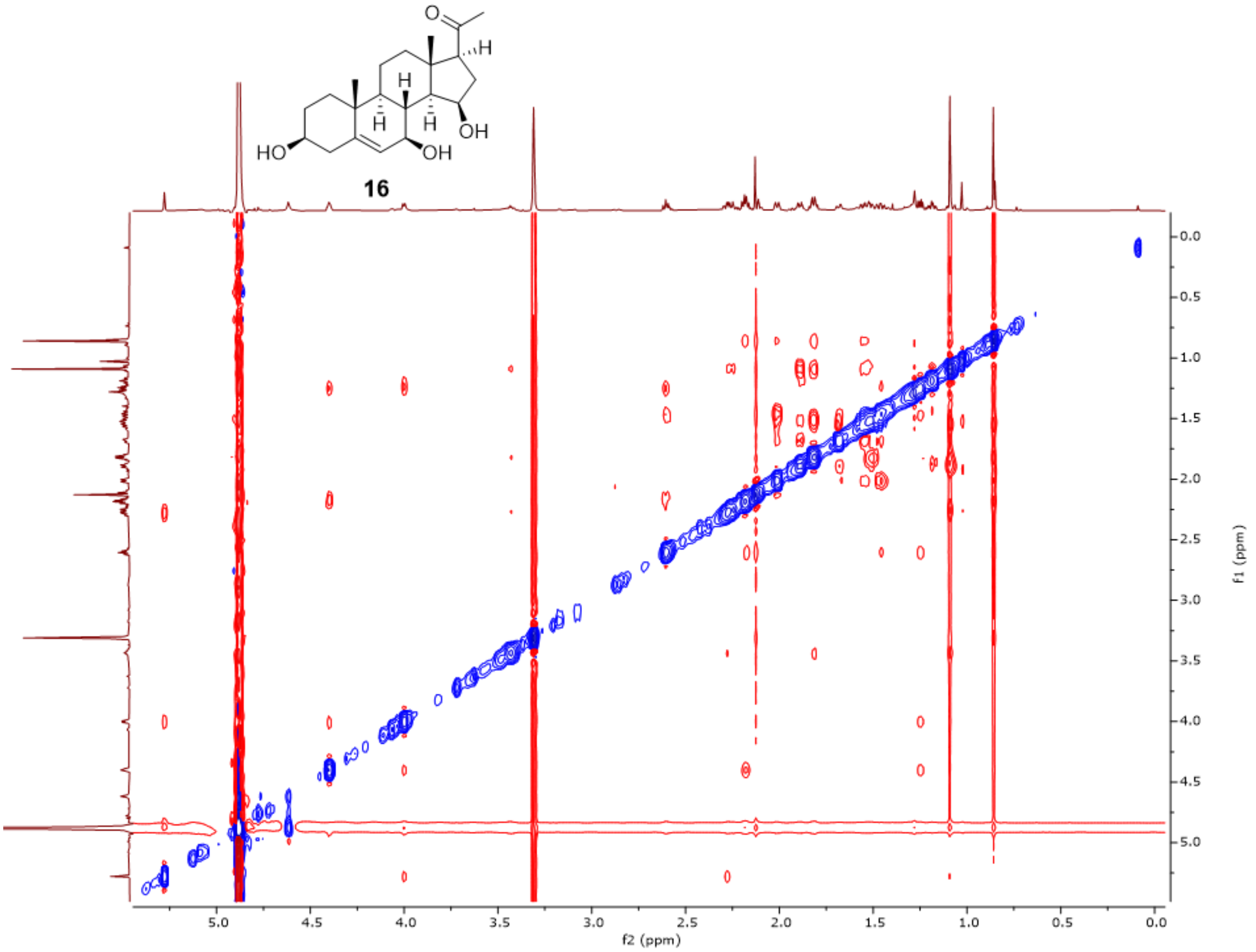


**Fig. S16**. ROESY spectrum (MeOH-$d_3$) of **16**.

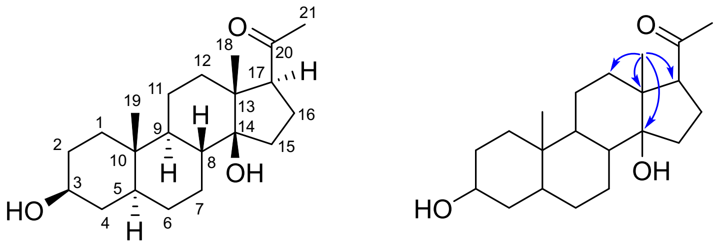

**18**

**Fig. S17**. Structure elucidation of 14β-hydroxyisopregnanolone (**18**). Blue single-headed arrows: key HMBC correlations. The C14 hydroxy group of 18 was tentatively assigned a β orientation by analogy with **10**, given the close structural similarity of their precursors and their formation by the same enzyme.

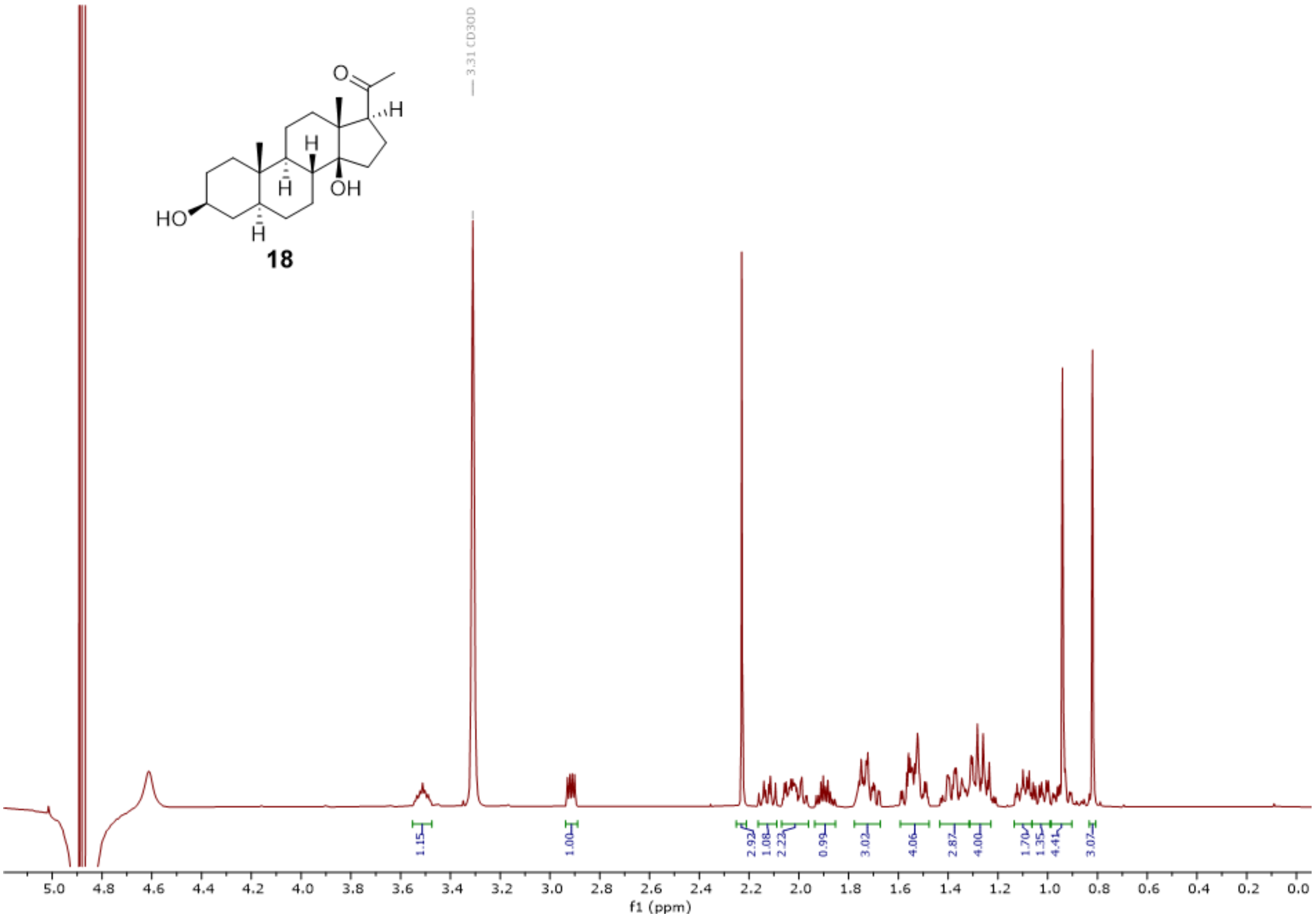


**Fig. S18**. [1]H-NMR spectrum of **18** (500 MHz, MeOH-$d_3$).

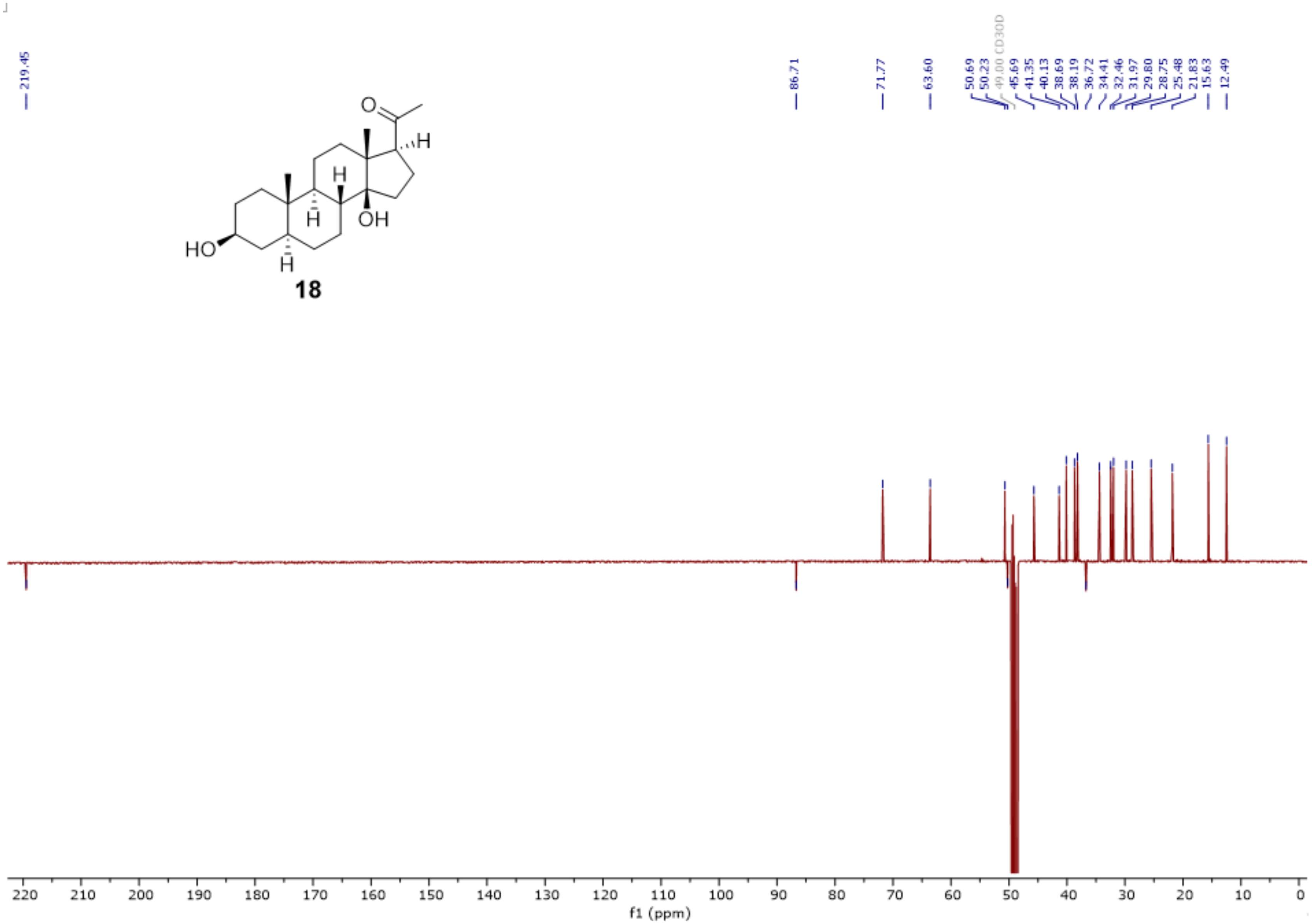


**Fig. S19**. $^{13}$C-DEPTQ spectrum of **18** (126 MHz, MeOH-$d_3$).

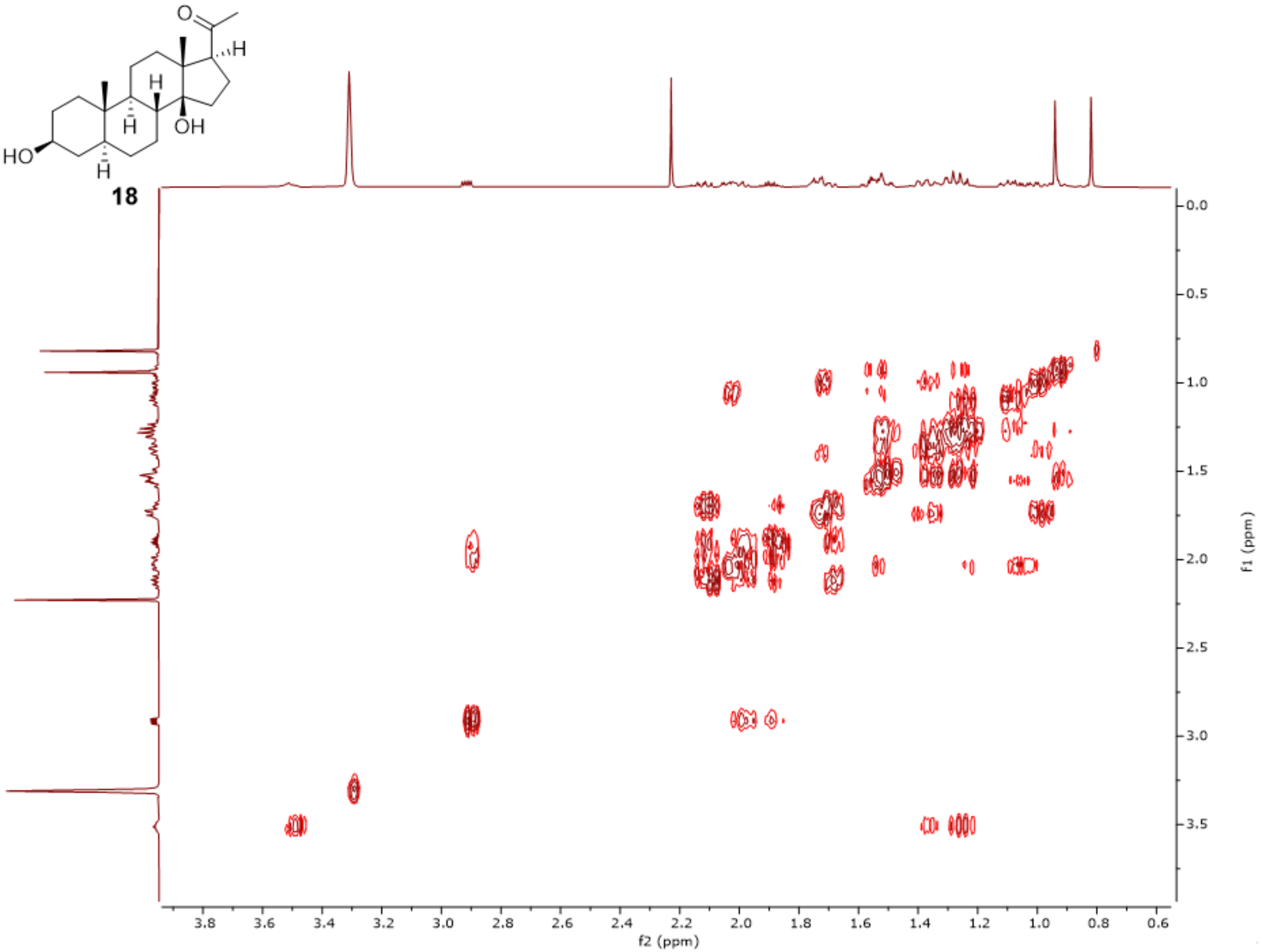


**Fig. S20**. $^{1}H$-$^{1}H$-COSY spectrum (MeOH-$d_3$) of **18**.

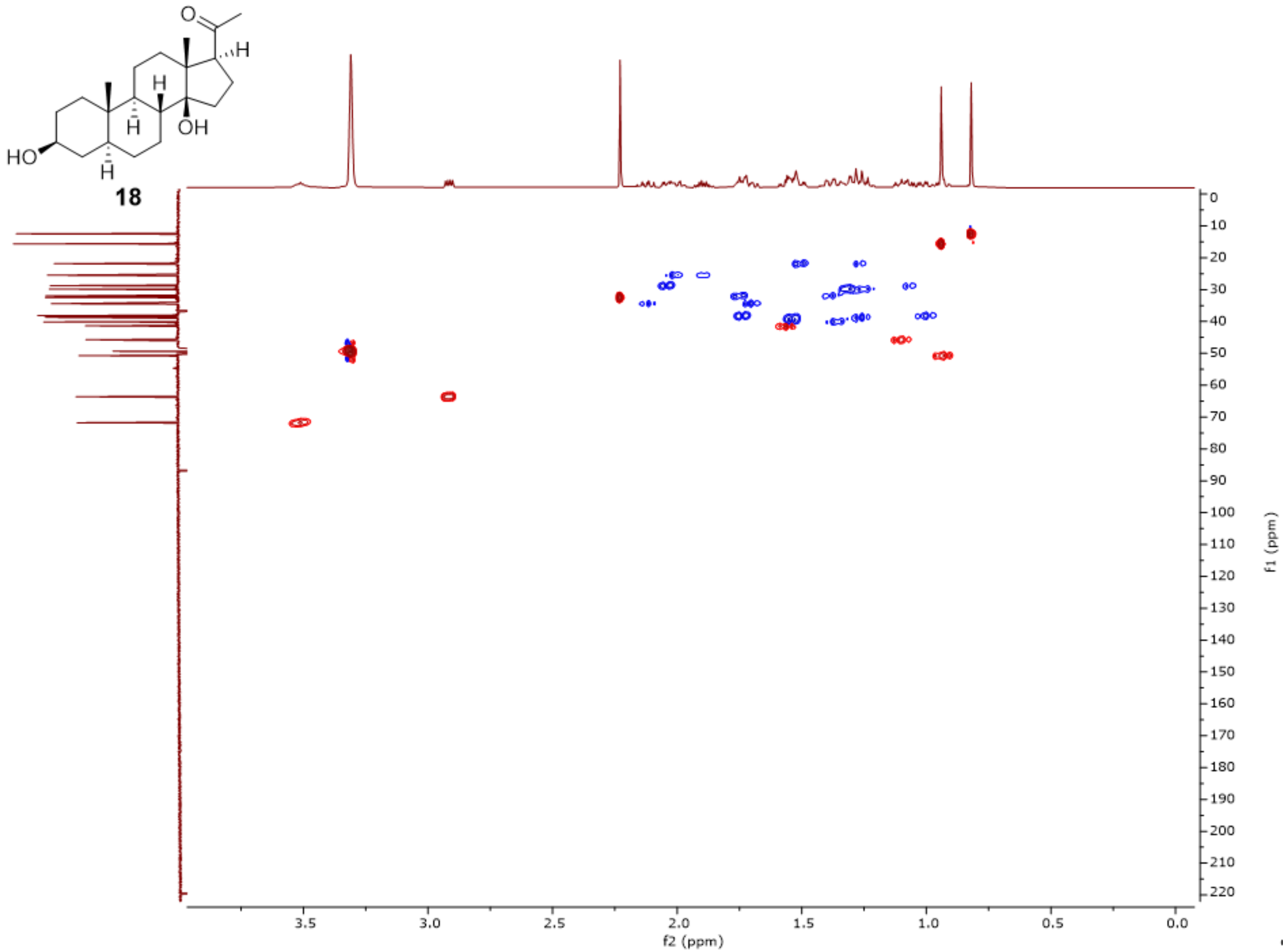


**Fig. S21**. HSQC spectrum (MeOH-$d_3$) of **18**.

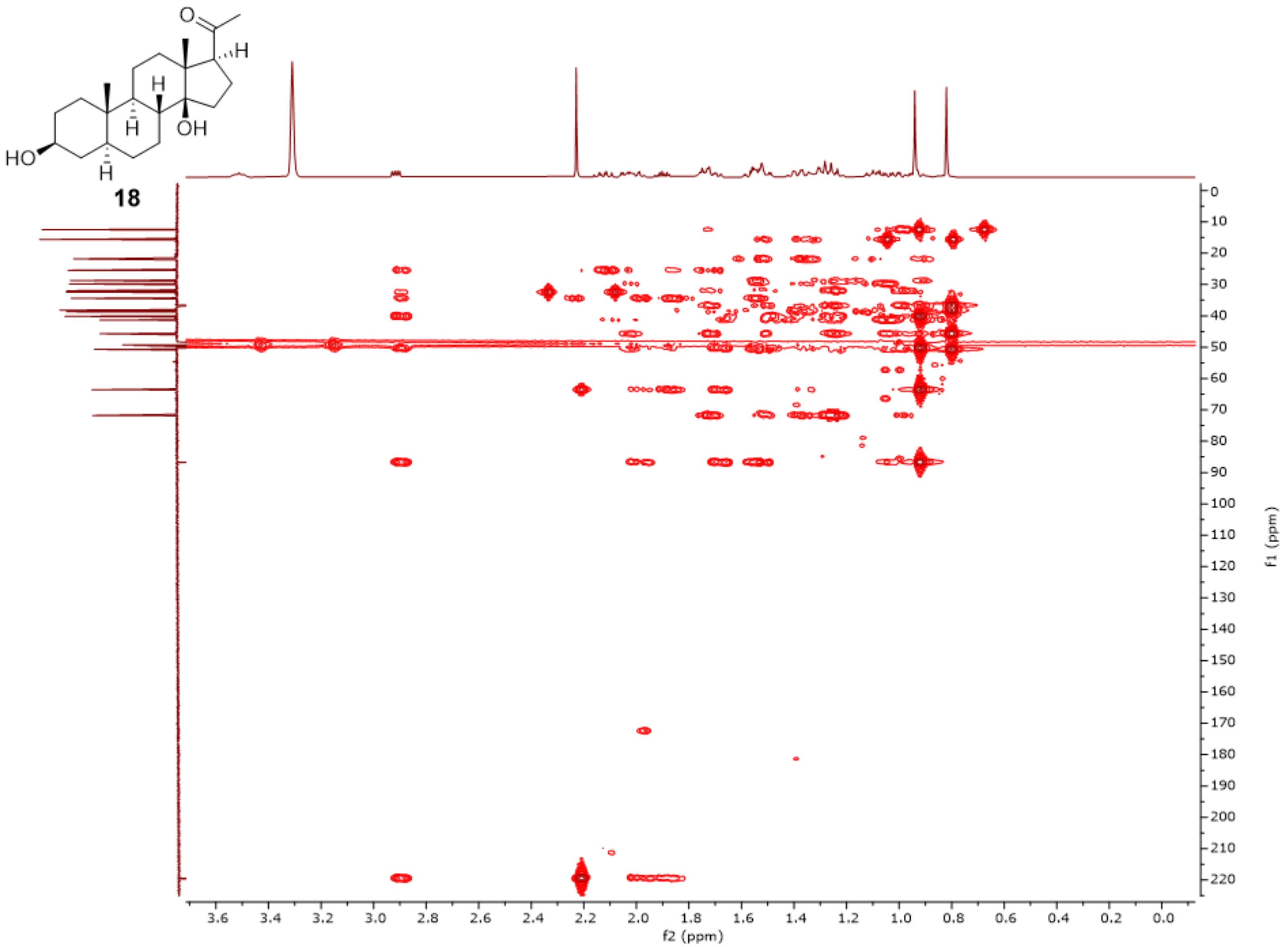


**Fig. S22**. HMBC spectrum (MeOH-$d_3$) of **18**.

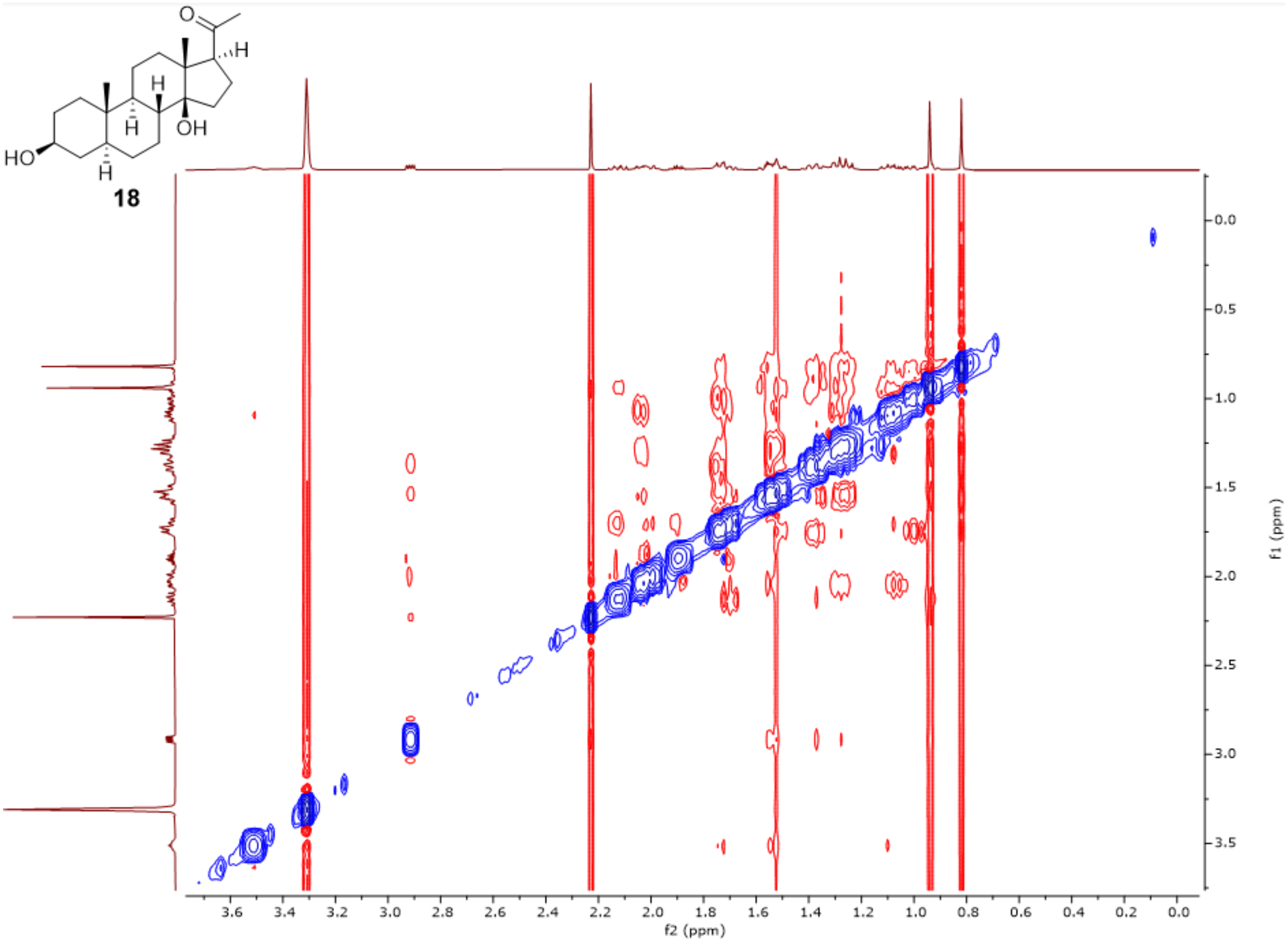


**Fig. S23**. ROESY spectrum (MeOH-*d*$_3$) of **18**.

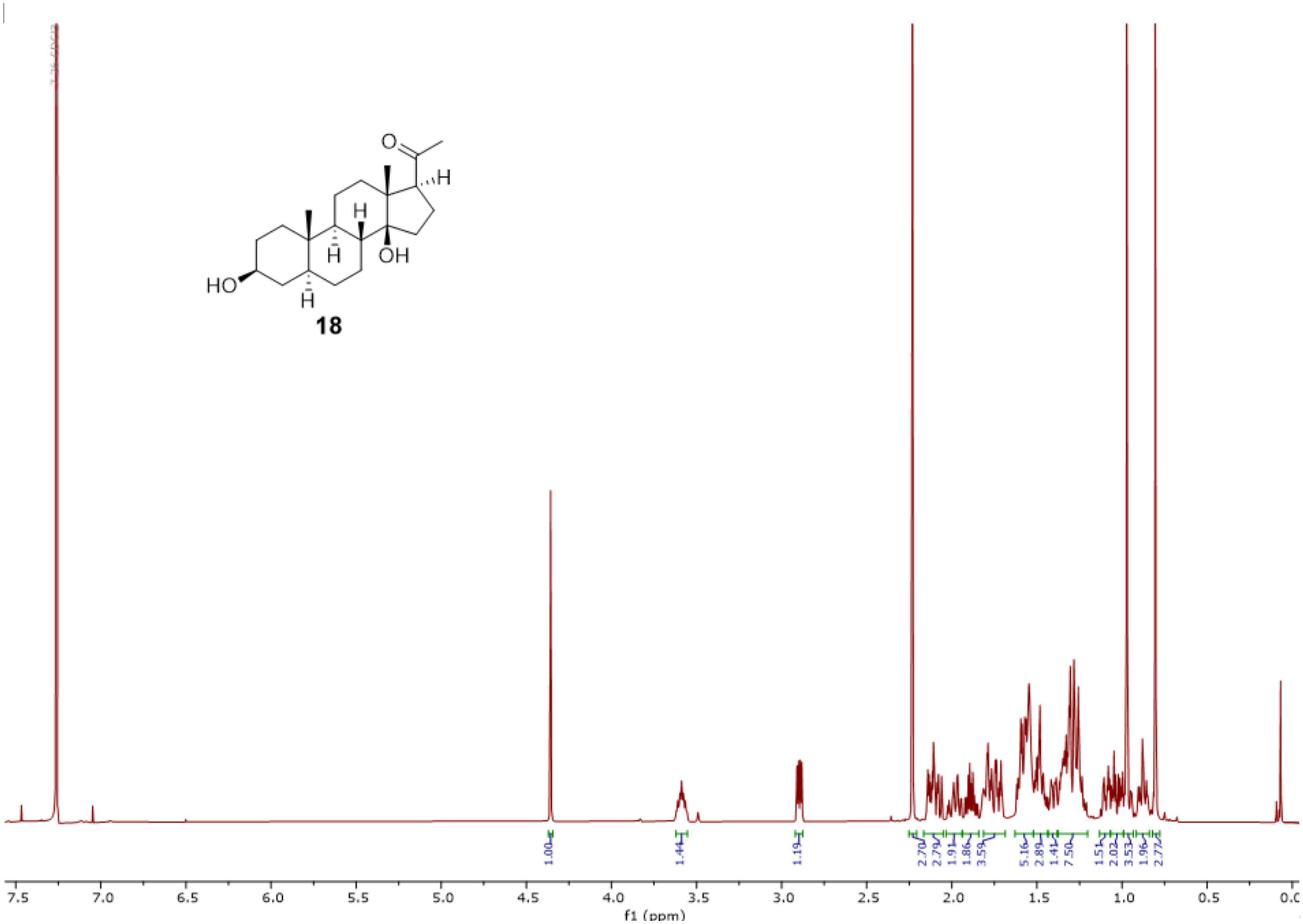


**Fig. S24**. [1]H-NMR spectrum of **18** (500 MHz, $CDCl_3$).

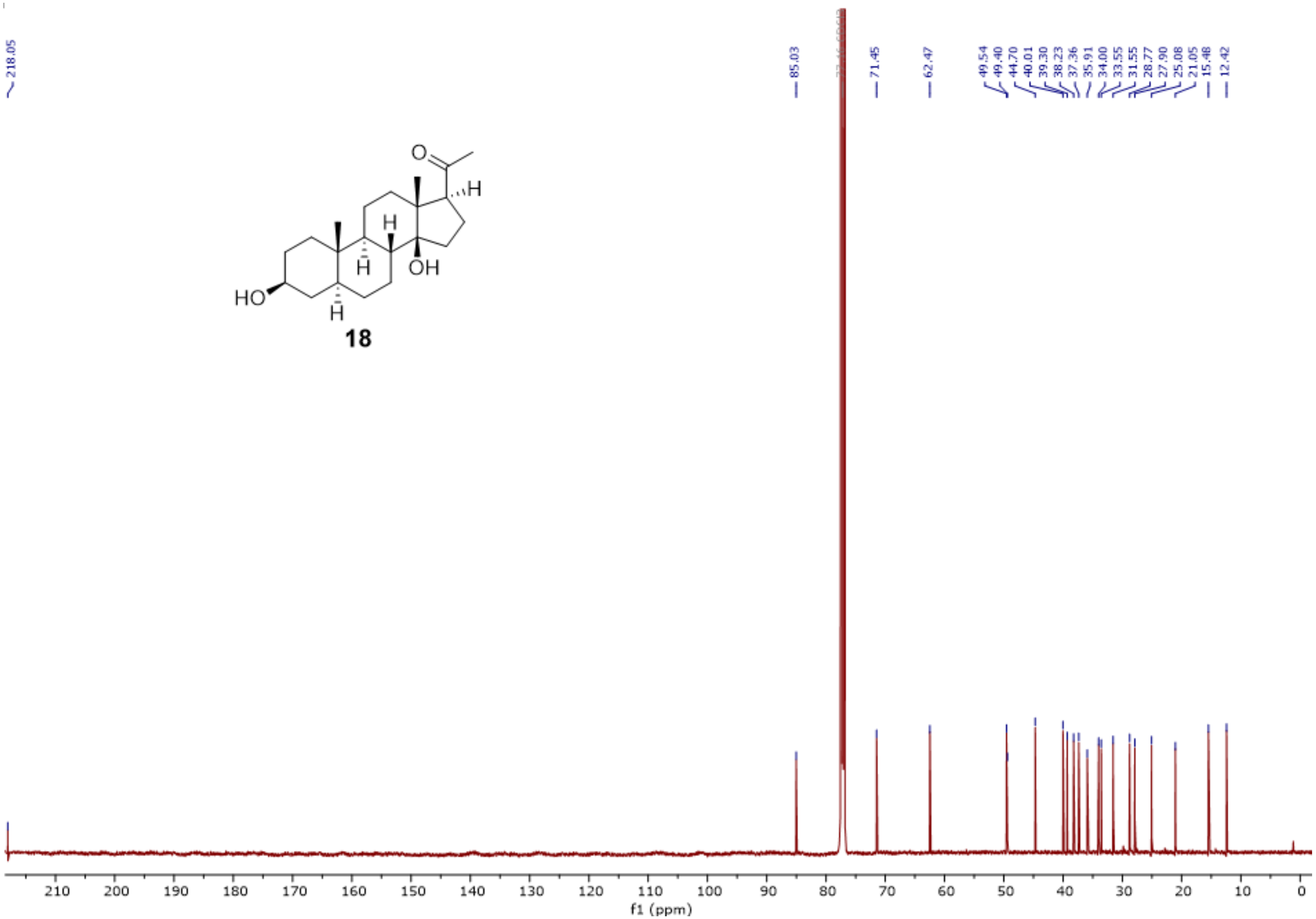


**Fig. S25**. $^{13}$C-NMR spectrum of **18** (126 MHz, $CDCl_3$).

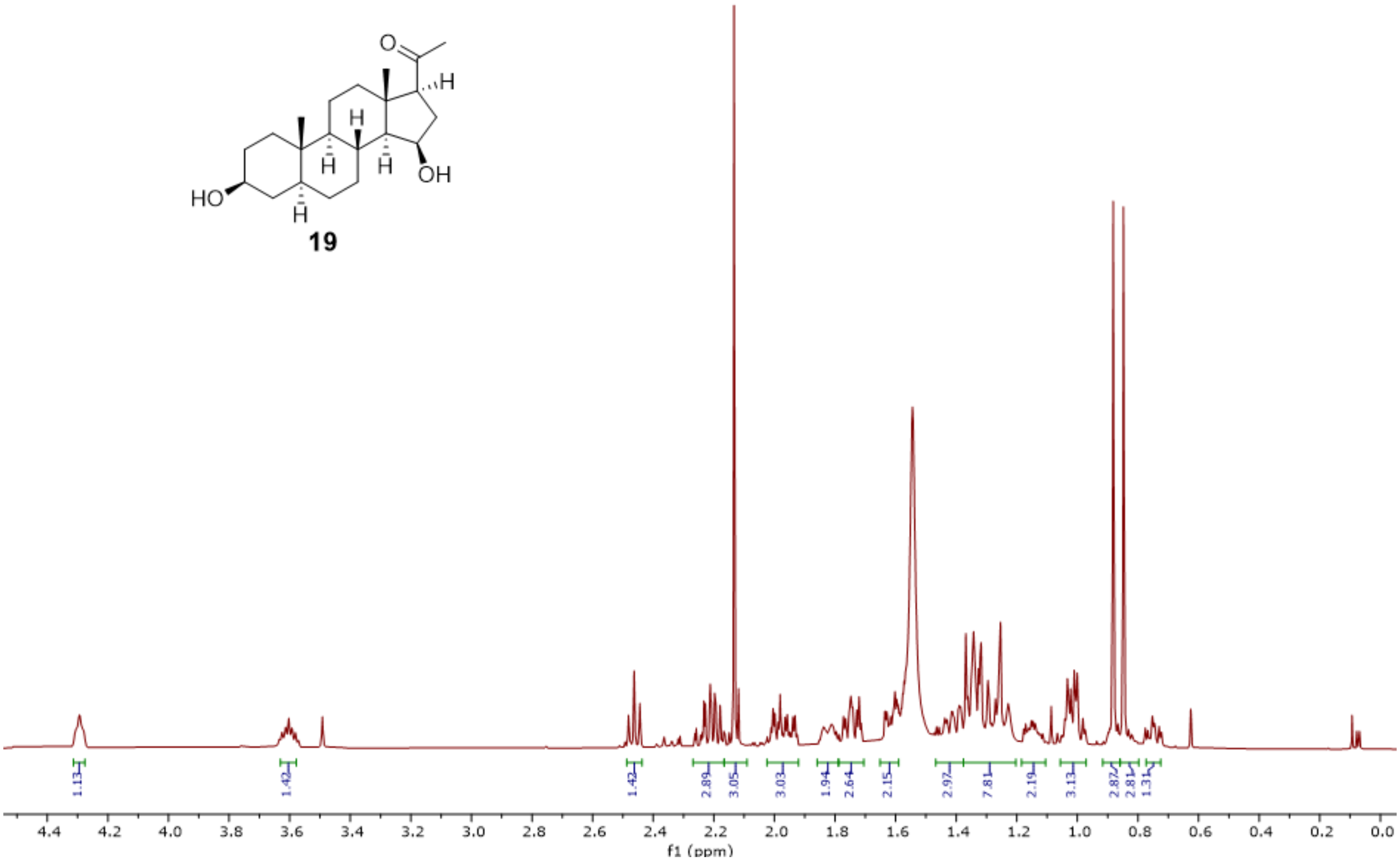


**Fig. S26**. [1]H-NMR spectrum of **19** (500 MHz, $CDCl_3$).

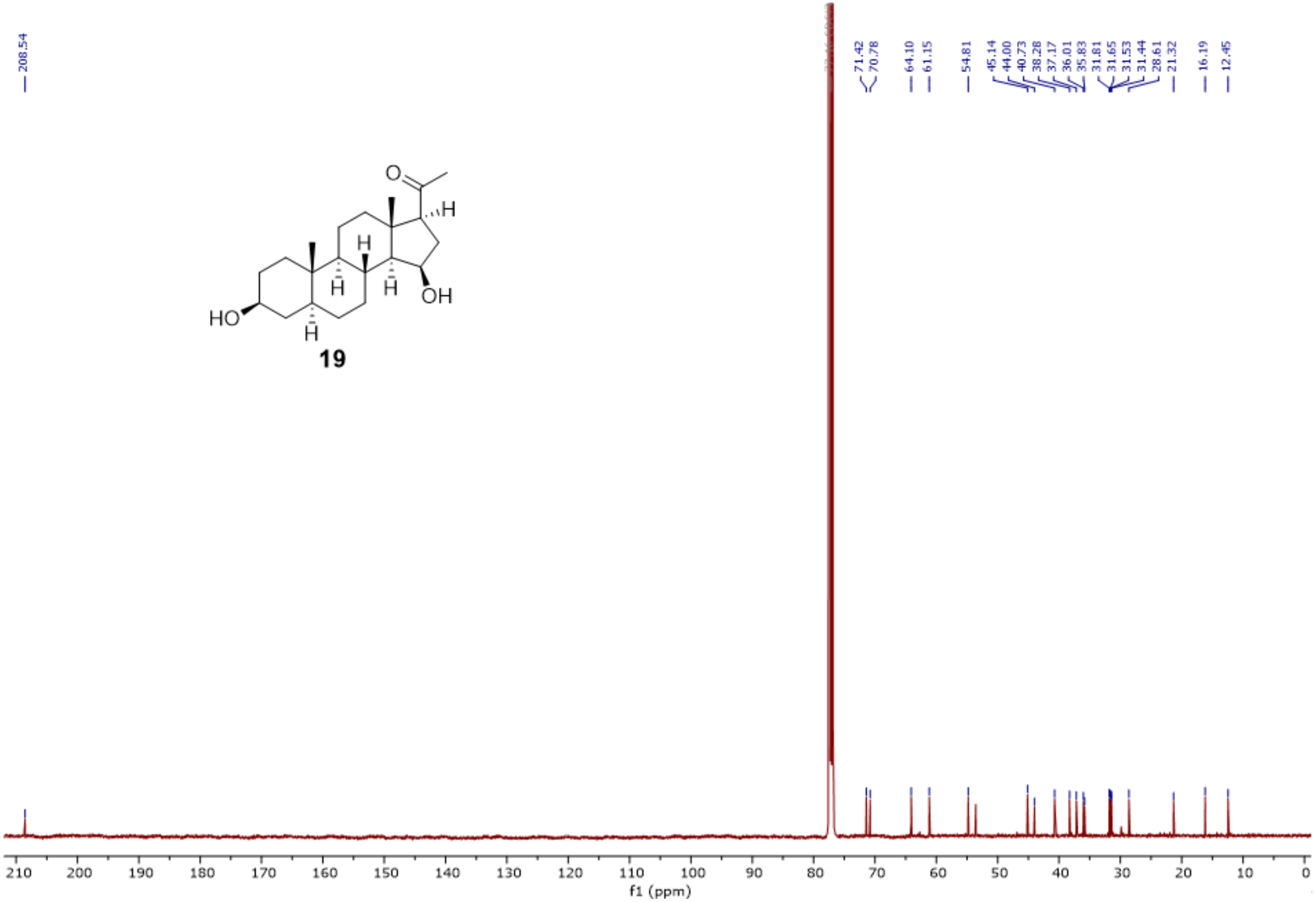


**Fig. S27**. $^{13}$C-NMR spectrum of **19** (126 MHz, $CDCl_3$).

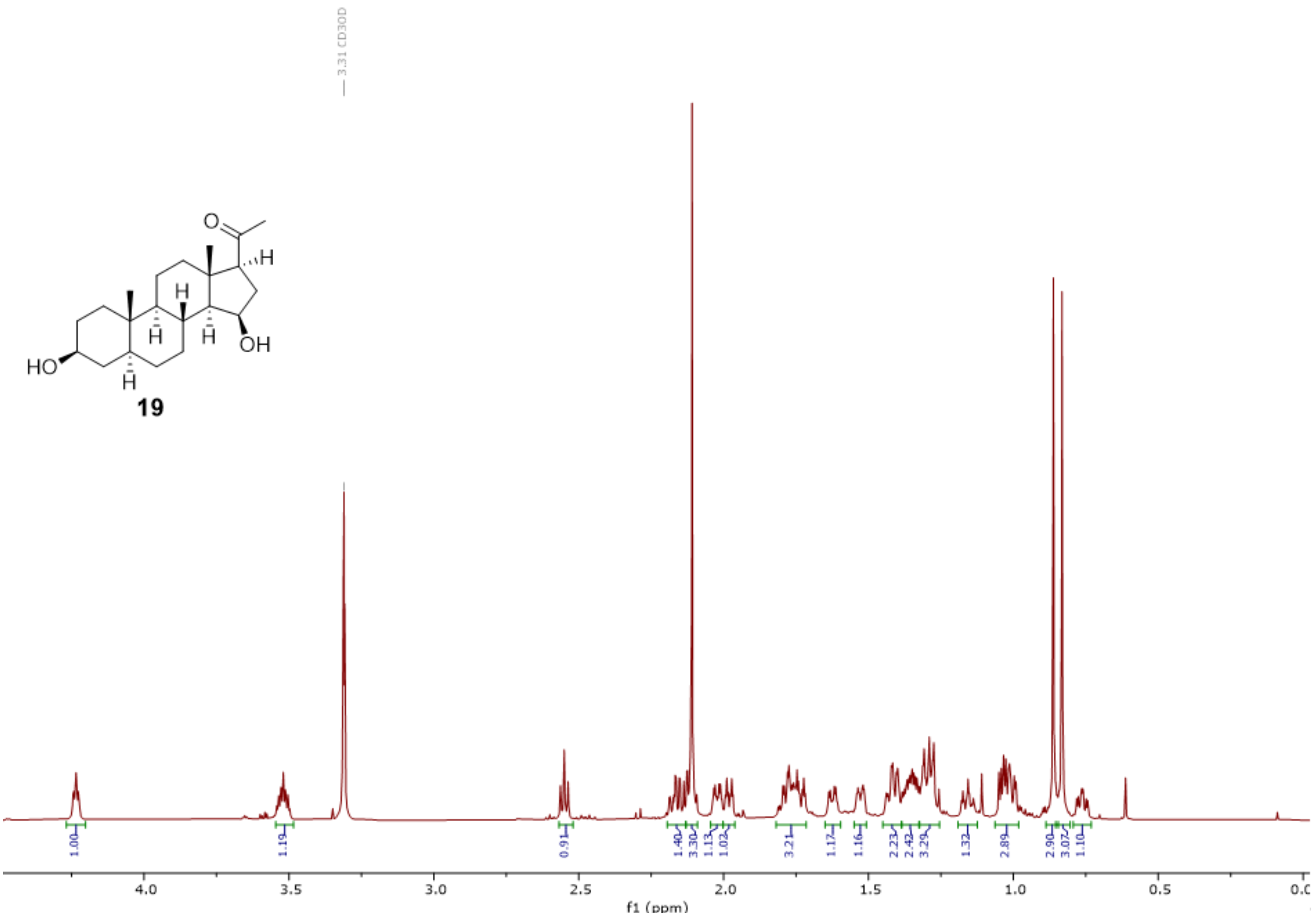


**Fig. S28**. $^1$H-NMR spectrum of **19** (700 MHz, MeOH-$d_3$).

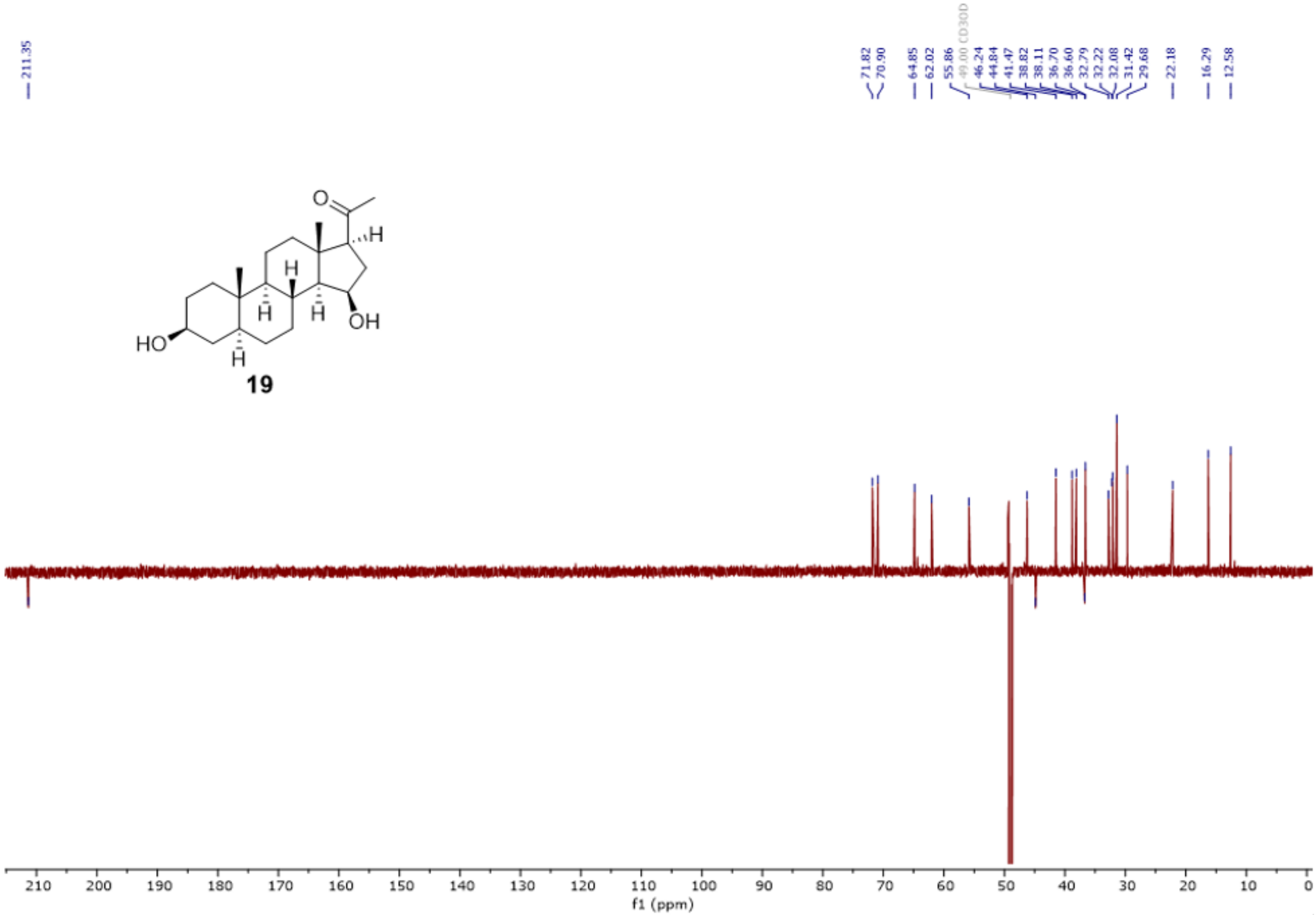


**Fig. S29**. DEPTQ spectrum of **19** (176 MHz, MeOH-$d_3$).

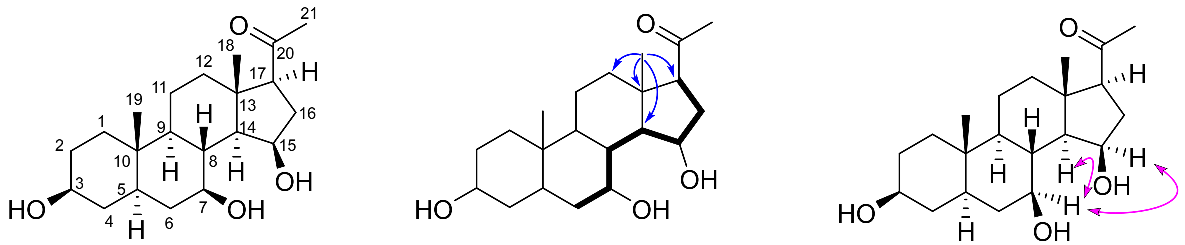


**20**

**Fig. S30**. Structure elucidation of 7β,15β-dihydroxyisopregnanolone (**20**). Bold: key $^{1}$H,$^{1}$H-COSY, blue single-headed arrows: key HMBC, and pink double-headed arrows: key NOESY correlations.

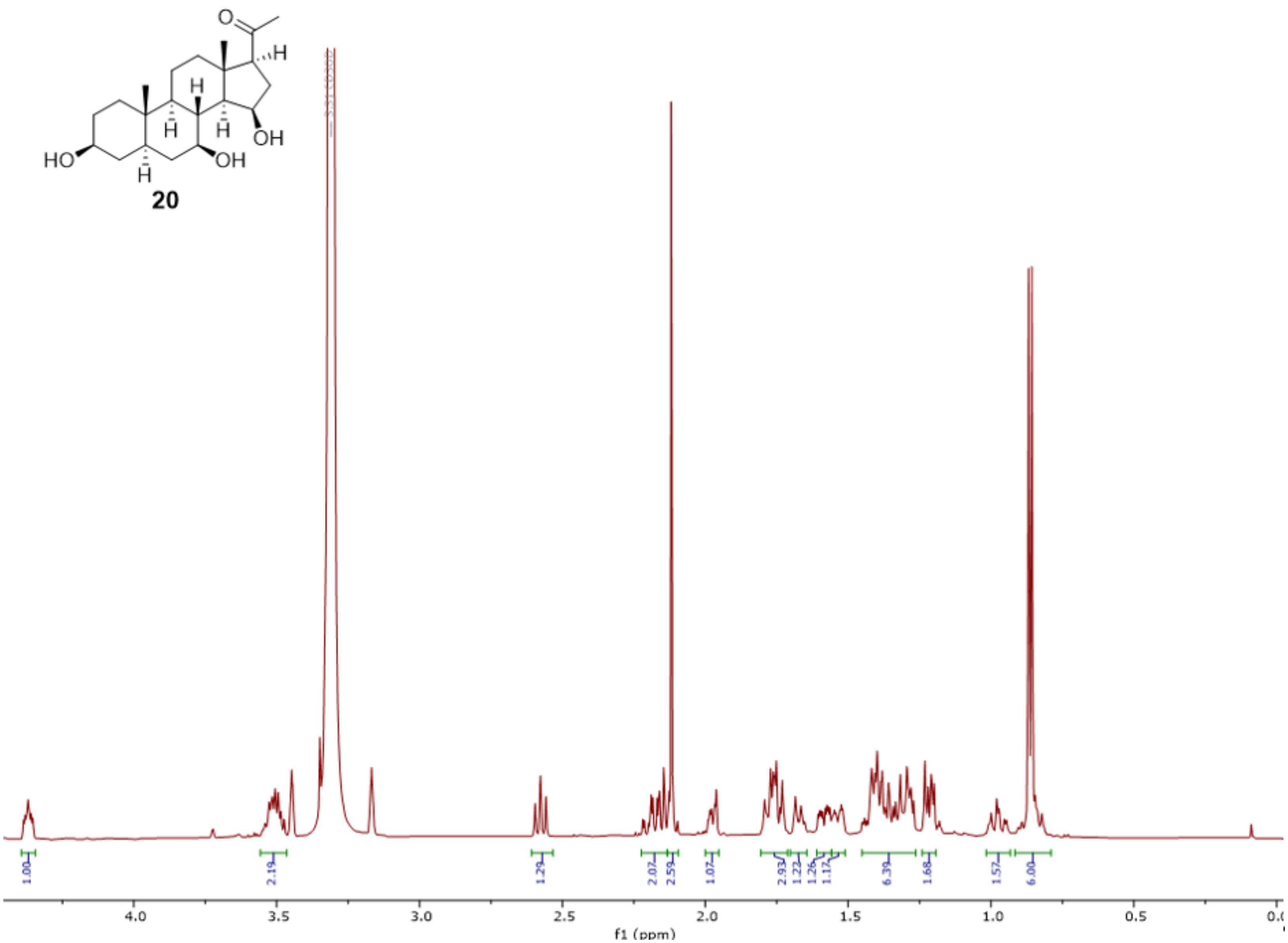


**Fig. S31**. $^{1}$H-NMR spectrum of **20** (500 MHz, MeOH-$d_3$).

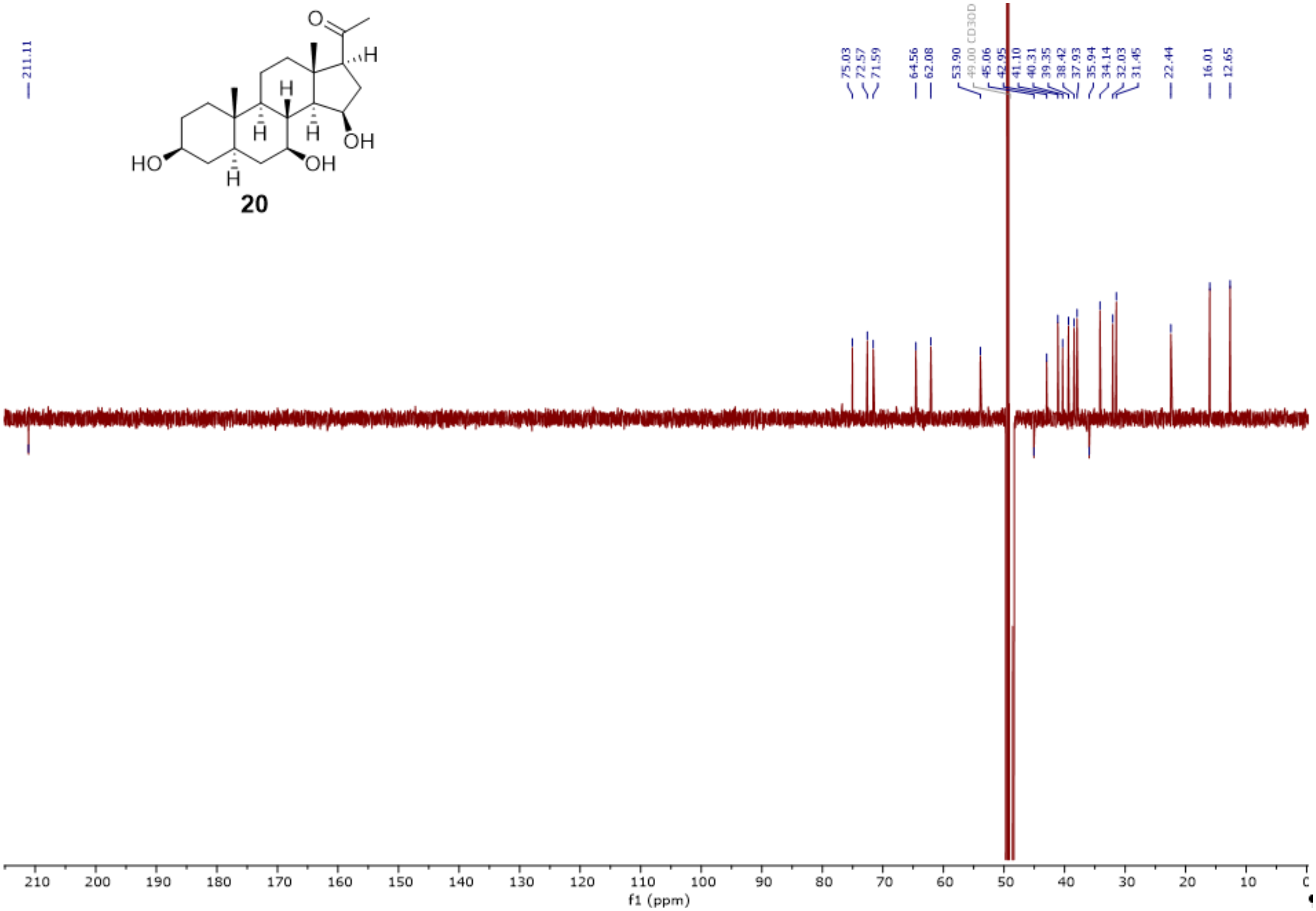


**Fig. S32**. DEPTQ spectrum of **20** (126 MHz, MeOH-$d_3$).

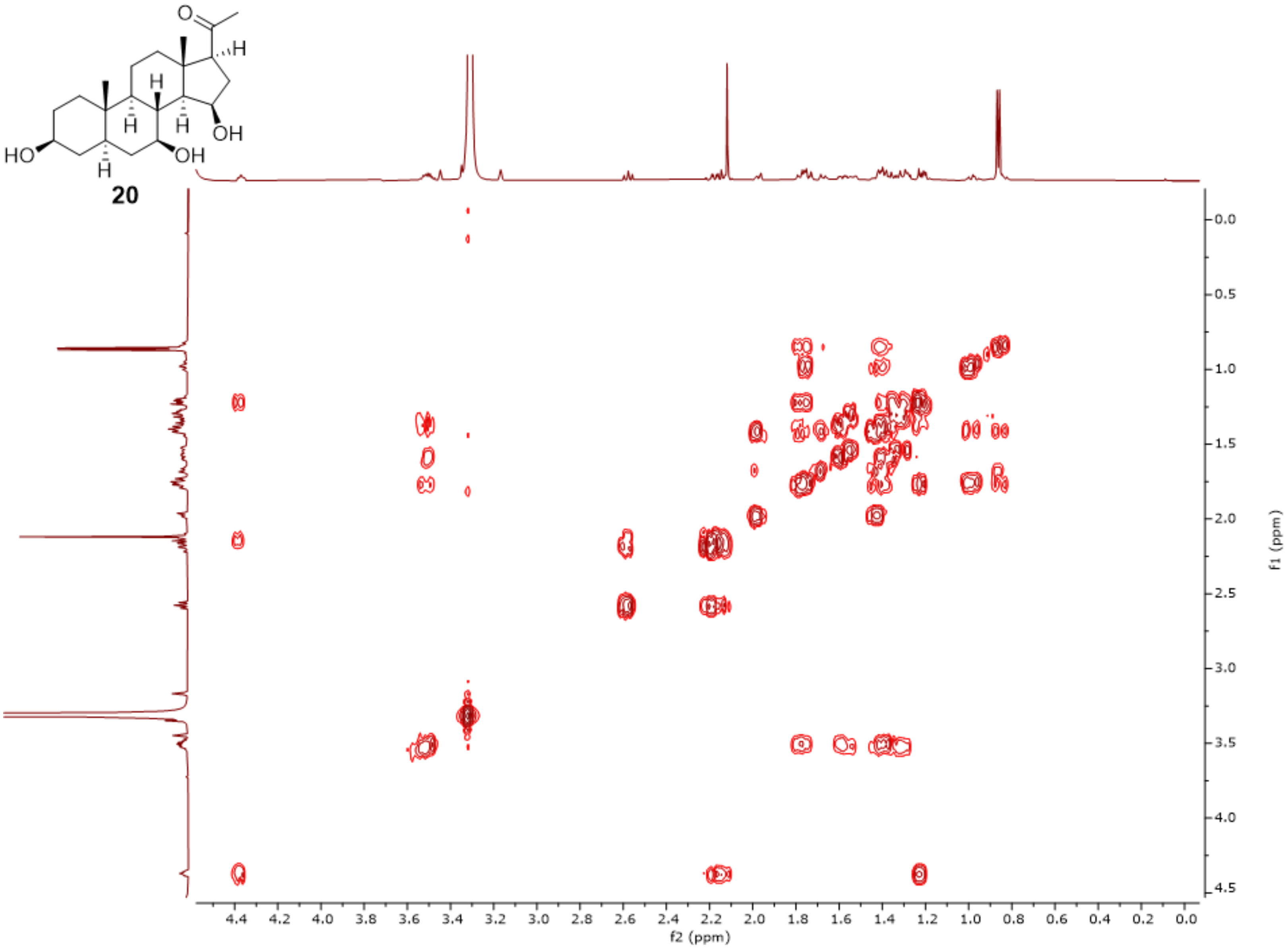


**Fig. S33**. $^{1}$H-$^{1}$H-COSY spectrum (MeOH-$d_3$) of **20**

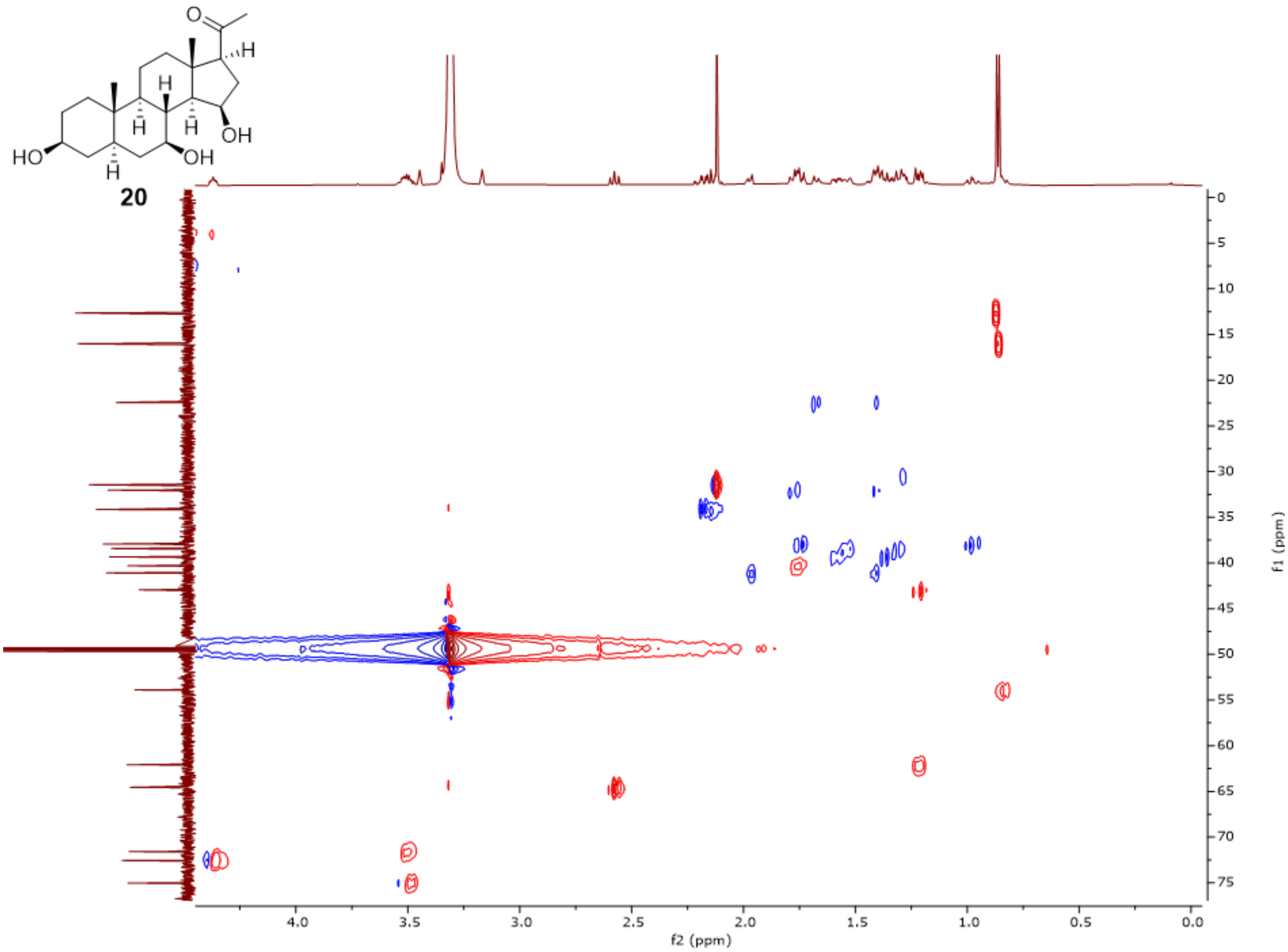


**Fig. S34**. HSQC spectrum (MeOH-$d_3$) of **20**.

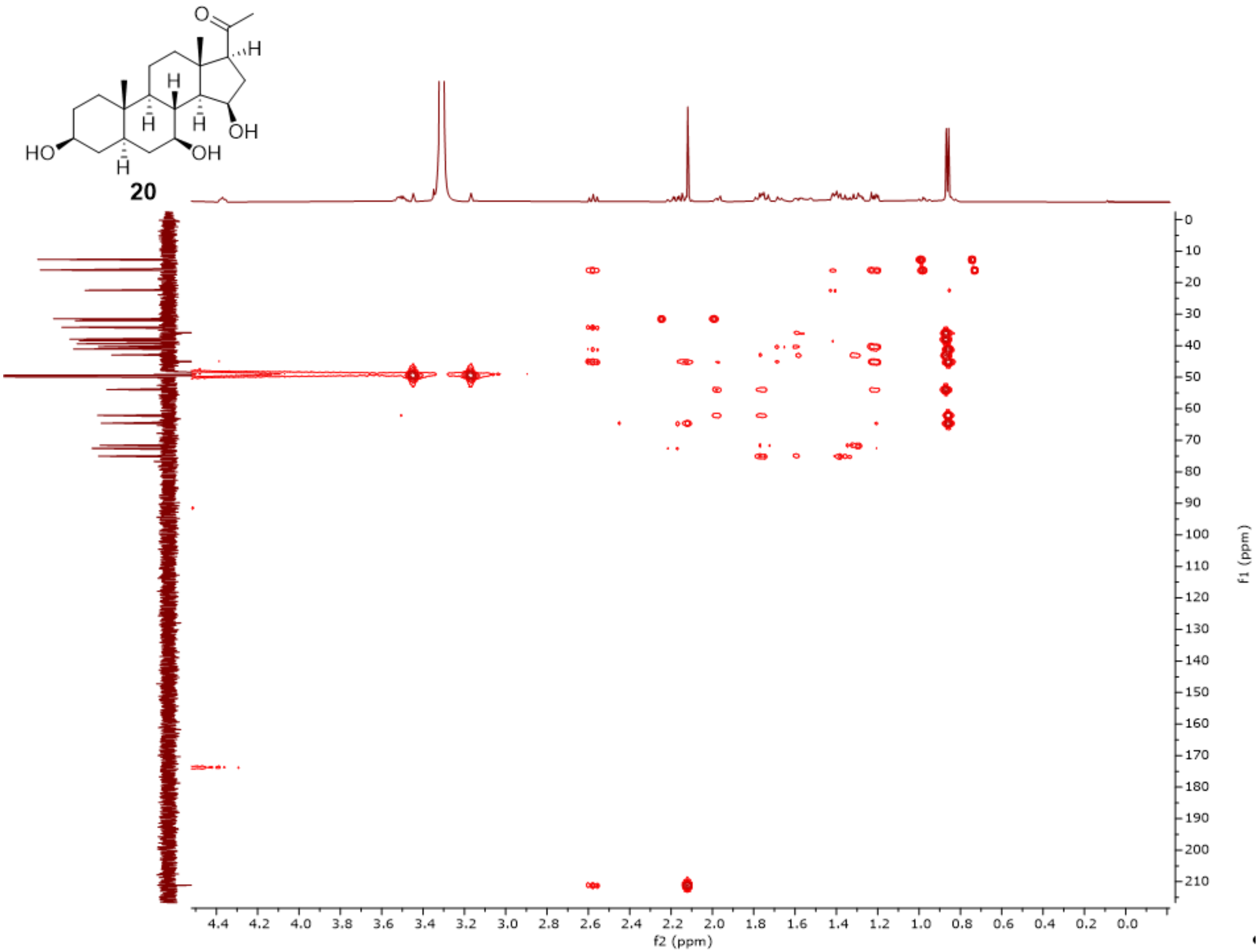


**Fig. S35**. HMBC spectrum (MeOH-$d_3$) of **20**.

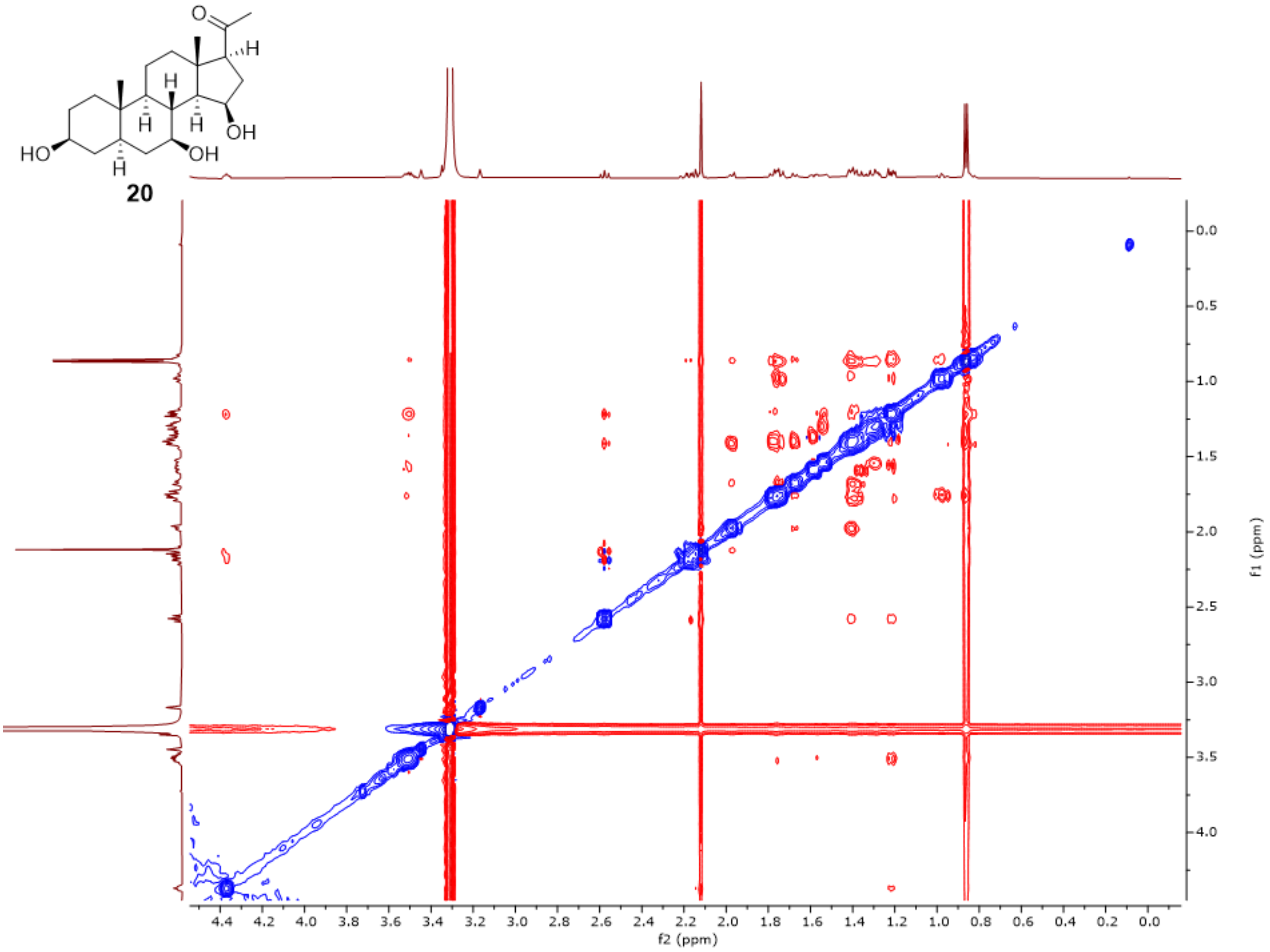


**Fig. S36**. ROESY spectrum (MeOH-$d_3$) of **20**.

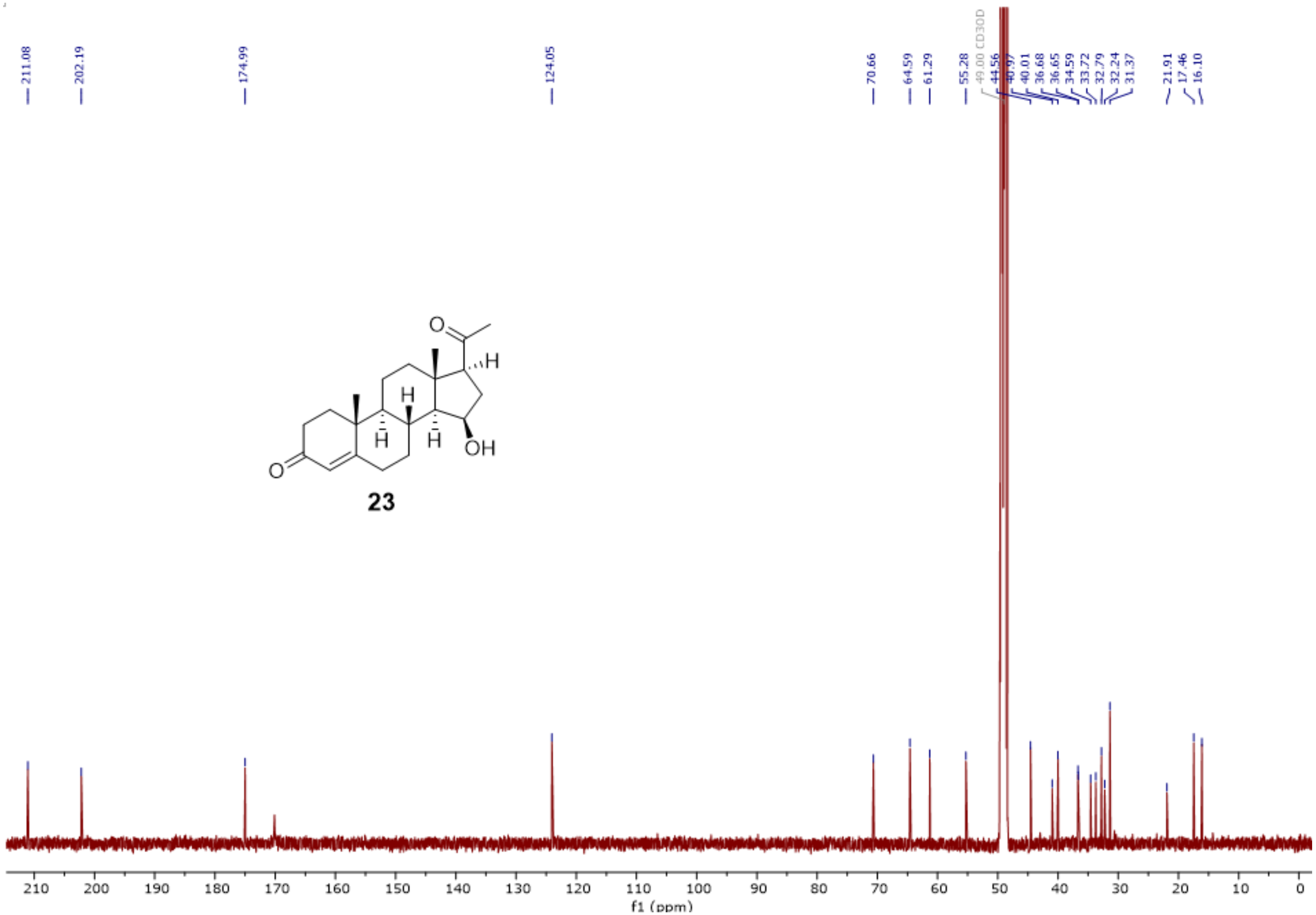


**Fig. S37**. $^{13}$C-NMR spectrum of **23** (500 MHz, MeOH-$d_3$).

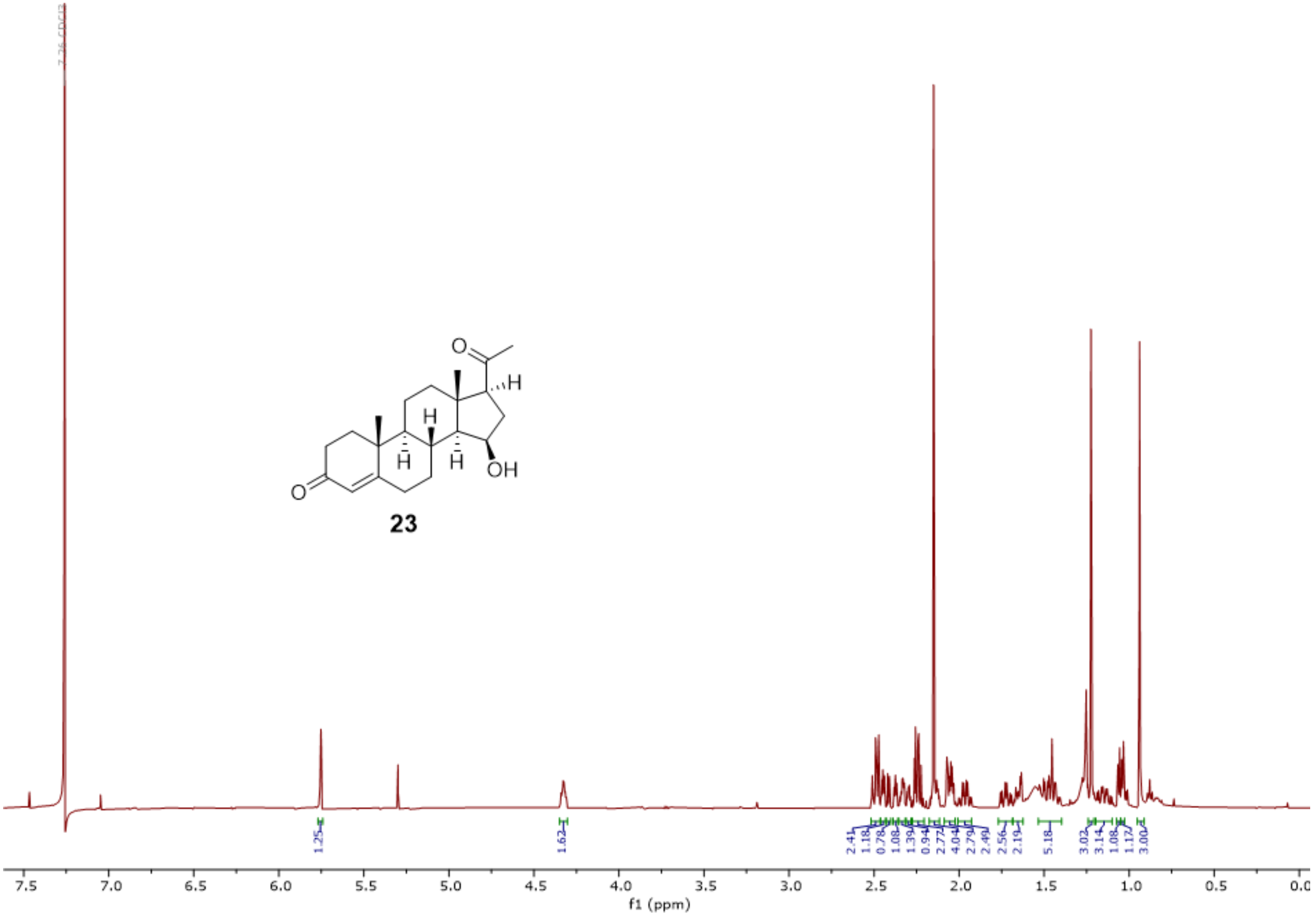

**Fig. S38**. [1]H-NMR spectrum of **23** (500 MHz, CDCl3).

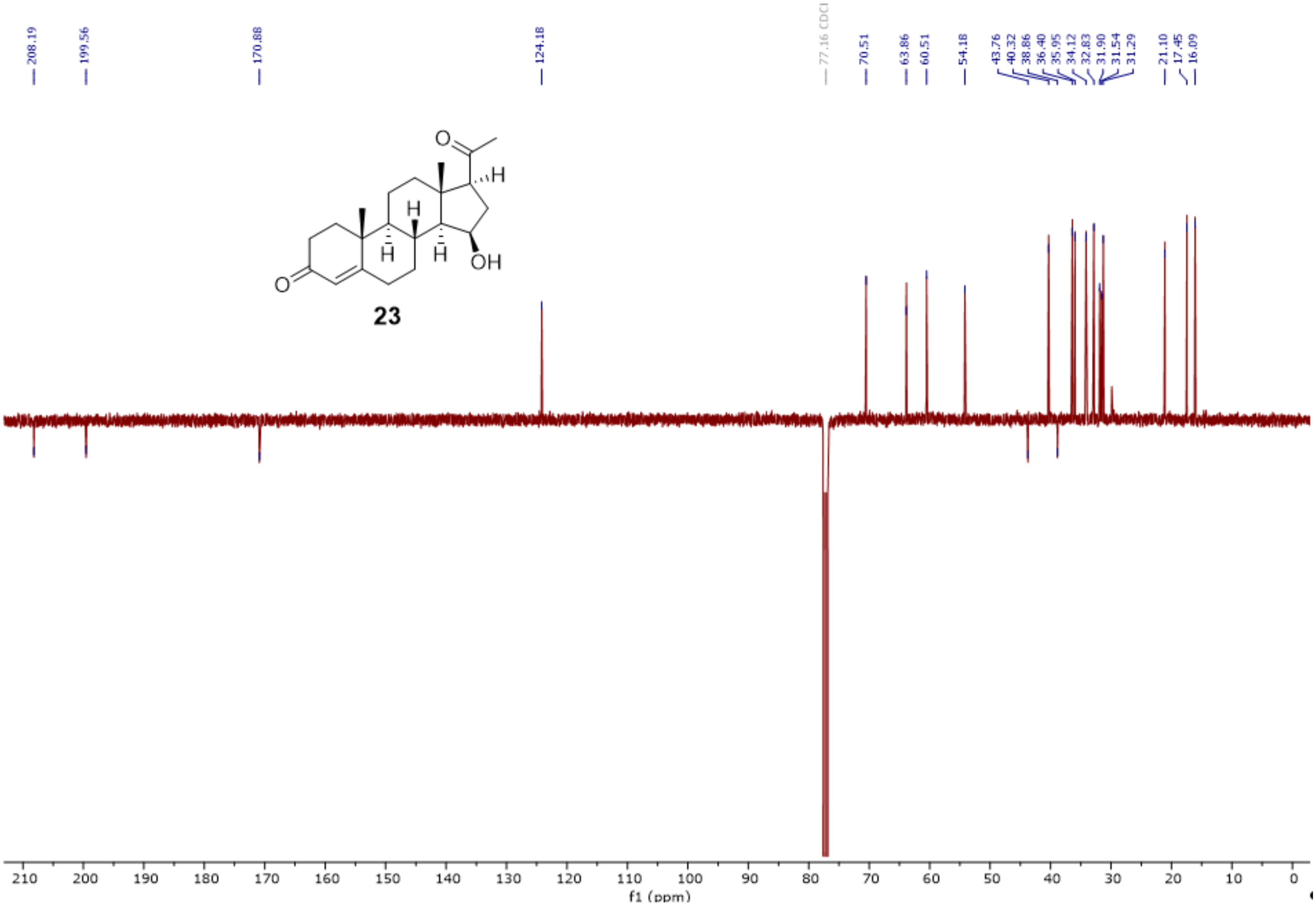


**Fig. S39**. DEPTQ spectrum of **23** (500 MHz, $CDCl_3$).

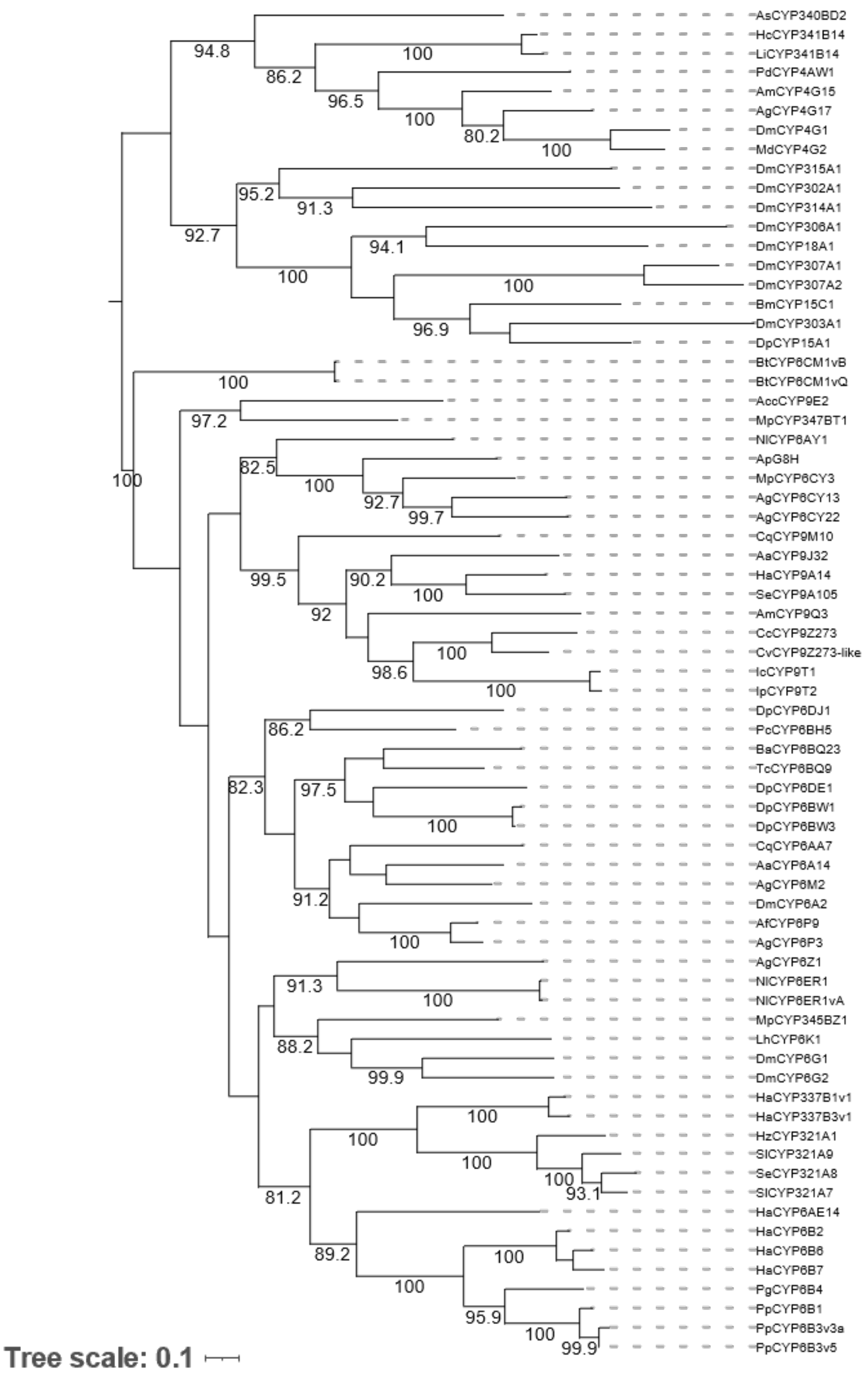


**Fig. S40**. Phylogenetic relationships of *Cc*CYP9Z273 and representative insect P450s. Maximum-likelihood phylogenetic tree of *Cc*CYP9Z273 from *C. coerulans*, its homolog *Cv*CYP9Z273-like from *C. varians*, and functionally characterized insect P450s from different clans. Bootstrap support values above 80% are shown at the corresponding nodes based on 1,000 bootstrap replicates. Branch lengths represent the estimated number of amino acid substitutions per site. Dashed lines connect terminal branches to sequence labels.

```
CcCYP9Z273    1    MPFTFFHEFEFNMWTIAVLVILTAAALYYILVRPLSYWKRLNVTQANPVILLGDYWTYIFGLESYAEMLR  70
Cv homolog    1    ------------MLTISLLVILTAVAFYHLLIRPLSYWKRKHVKQGHPVILLGDYWRYFFRMESFAELHR  58

CcCYP9Z273   71    RLYNQFPNERYIGIYQLRTPTLMLRCPELIKRVTVKDFDHFVDHRNVTPESADPLWGKNLFSLDEDRWRE  140
Cv homolog   59    RLYNQFPNERYYGVYQYLTPTLVVRDPELVKRITVKDFDHFVDHRAVMPESSDPLWAKNLFSLTGERWRE  128

CcCYP9Z273  141    MRPVLQPAFTSSKMKLMFQLVTECAEDFVAFFLEKNQDLIEVELRDTLSRFANDAIATSAYGVKVNSVRE  210
Cv homolog  129    MRPVLSPAFTGSKMRLMFGLISDCAKDFVSFFLEKNQDVLEVDFRDTFTRFTNDAIATSAYGIEVNSLRT  198

CcCYP9Z273  211    PENEFYTTSIDTFTLTPWVAFKFFASFYLSPFWLTLFNLRVFTKKNSEFFIRVIDETIKEREEKGIIRPD  280
Cv homolog  199    PDNEFYTKSIKAFTLTSAVALKFFAGLLLPTFIINLLKLRMFEKEISDFFTNIVEETIKIREEKGIVRPD  268

CcCYP9Z273  281    LIHLLMEARKEQNRKYNEKIAEREKGCPA---SELGKPIPEKVEITNVDIAAQAMVFSLGGFDSSASVMC  347
Cv homolog  269    MIHLMMEA-KKGYYKYEEDTEARDTGFAAVKESDIGKQTRRKIEITNSDIAAQALIFSLGGFDSSAALMC  337

CcCYP9Z273  348    FLAYELAIHQDLQQKLRDEINDTLAKCNGVLTYEALMEMKYIDMVVSEALRKWSNGVILDRVSTKPYDVV  417
Cv homolog  338    FVGYELAVHQDIQEKLRYEIQDTLTECNGTITYEALLRMKYMDMVLSETLRRWSNGVIADRICTKPYTIE  407

CcCYP9Z273  418    PLTPDEKPFRIPVRTVIMIPGYALQHDPKFFPDPDRFDPERFNDENKRNVDPYTNLPFGIGPRSCLGQRF  487
Cv homolog  408    PTTAEEEPLHLAVGTLVMAPSHALQHDPKYFPDPDRFDPERFSDENKGSIDPYTYLPFGIGPRNCIGSRF  477

CcCYP9Z273  488    GRMETKVLFFHLVSKFKIVPTAKSVIPMKPLLYTFNMTAEDGFKFAFERIKEDQK  542
Cv homolog  478    ALLETKLLFFHLLSKFKIIPTPKSGIPLKLSKQTLNMSAEGGFWFAFERLKEDQL  532
```

■ Identical residues    Nonidentical residues and gaps are unshaded.

**Fig. S41**. Pairwise amino acid sequence alignment of *Cc*CYP9Z273 and its closest P450 homolog from *Chrysolina varians*. The full-length amino acid sequences were aligned, revealing 64.6% sequence identity (352 identical residues across 545 aligned positions). Identical residues are highlighted in blue, whereas nonidentical residues and gaps are unshaded. Amino acid positions are indicated on both sides of the alignment.

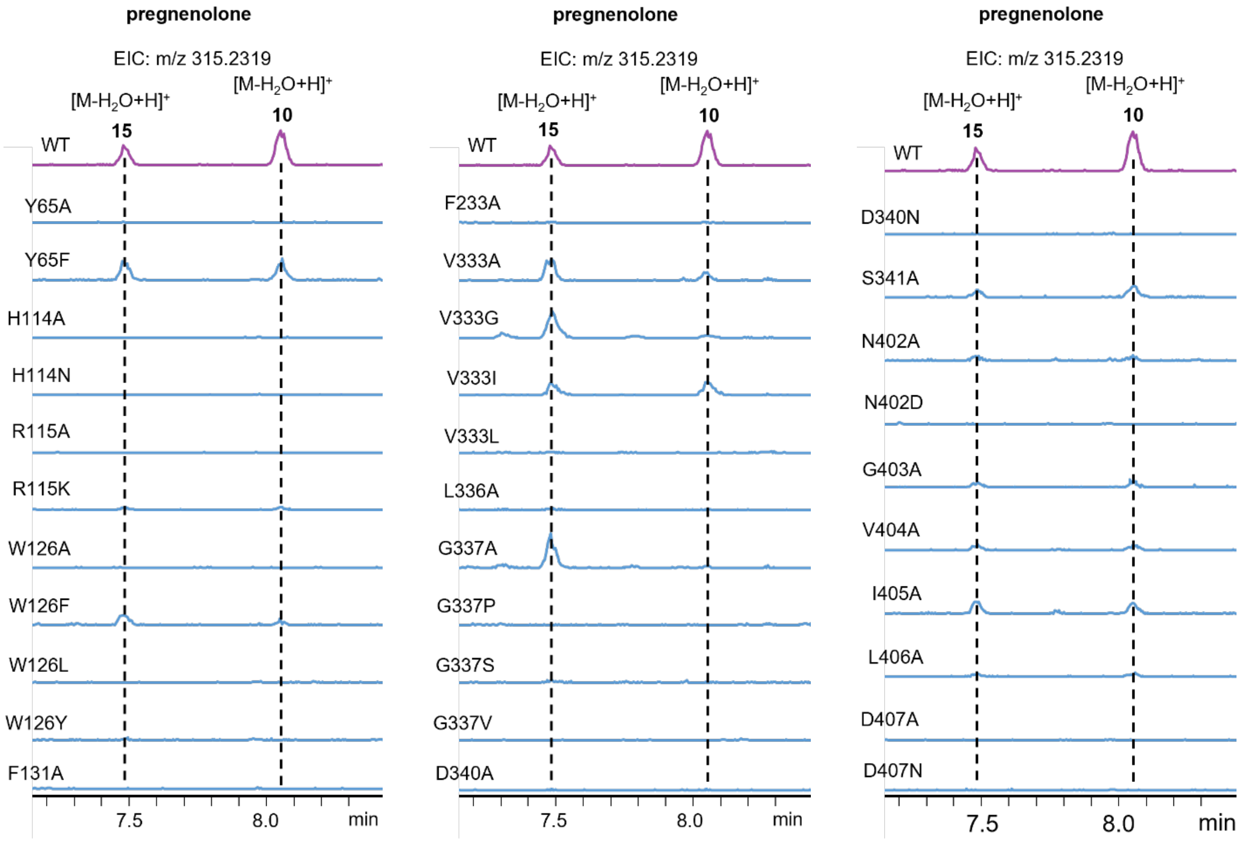


**Fig. S42**. Screening of *Cc*CYP9Z273 variants in yeast fed with pregnenolone (**9**). Extracted-ion chromatograms showing formation of **15** and **10** by yeast expressing wild-type (WT) or the indicated variants following pregnenolone (**9**) feeding. Representative chromatograms from three technical replicates (n = 3) are shown.

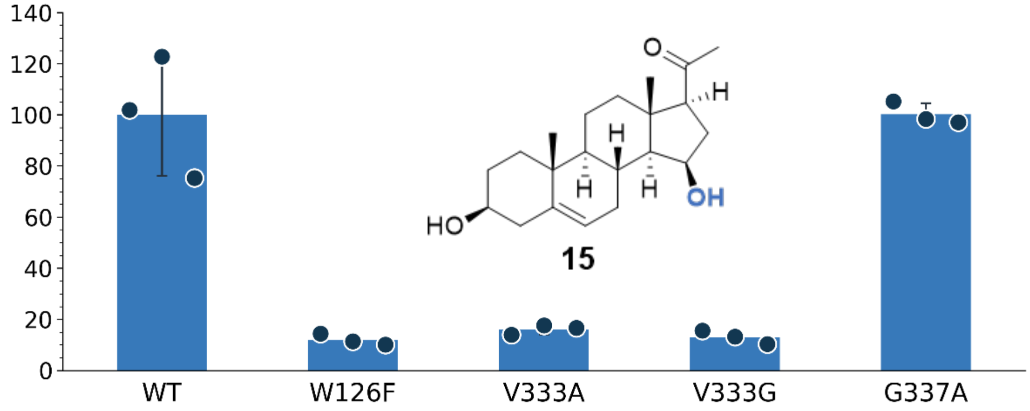


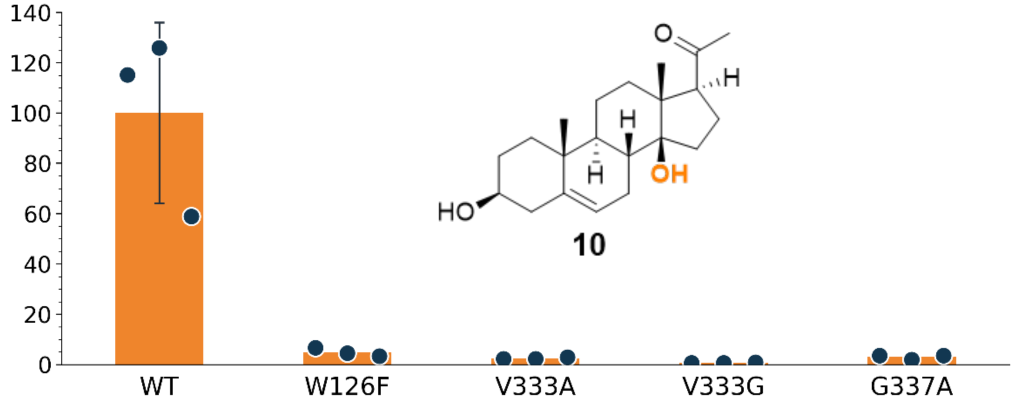


**Fig. S43**. Individual replicate data for pregnenolone (**9**) hydroxylation by *Cc*CYP9Z273 and its mutants. The y axes show the relative formation of **10** and **15** (% of WT), with formation of each product by WT normalized to 100%. Bars represent the mean ± SD of three technical replicates (n = 3), with individual data points shown as circles.

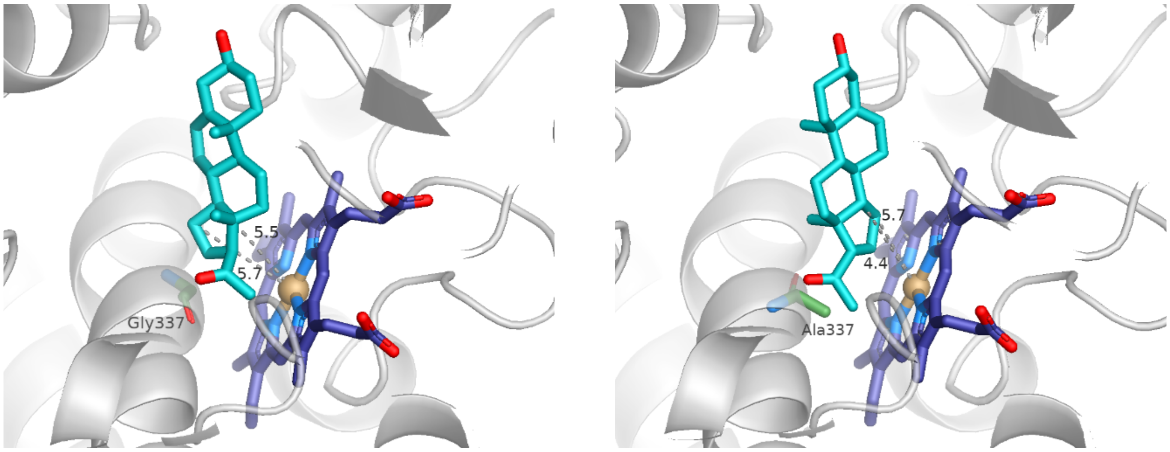


**Fig. 44**. Detailed comparison of representative productive-like docking poses of pregnenolone in WT *Cc*CYP9Z273 and the G337A variant. This figure provides a detailed view of the docking models shown in the main text. Pregnenolone is shown in teal, the heme cofactor in blue, and residue 337 in green, with the protein backbone shown as a semi-transparent gray cartoon. Dashed lines indicate the distances from C14 and C15 of pregnenolone to the heme iron. In the WT model (left), the C14–Fe and C15–Fe distances are 5.53 and 5.72 Å, respectively, whereas in the G337A model (right), the corresponding distances are 5.73 and 4.37 Å. The selected G337A pose shows a relative shift in substrate orientation, with C15 positioned closer to the heme iron. WT and G337A structures are shown in the same orientation for direct comparison.

**Table S02**. Isolation yields and NMR spectroscopic data of compounds **10**, **15**, **16**, **18**, **19**, **20** and **23**.

14β-Hydroxypregnenolone (caraumbellogenin, **10**). Yield: 1.5 mg (4.5 μmol, 7%)
$^{13}$C-NMR (126 MHz, DMSO): δ = 215.2, 140.0, 121.0, 84.1, 70.0, 62.6, 48.7, 45.5, 42.1, 38.4, 36.9, 36.6, 36.5, 33.4, 31.5, 31.4, 26.9, 23.4, 20.4, 19.3, 15.3 ppm. (**Fig. S11** and **S12**)
$^{13}$C NMR (126 MHz, $CDCl_3$): δ = 218.1, 139.2, 122.5, 85.3, 71.8, 62.7, 49.3, 46.1, 42.3, 38.8, 37.4, 37.0, 36.4, 34.6, 33.6, 31.7, 27.5, 24.7, 21.0, 19.6, 15.2 ppm. (**Fig. S13** and **S14**)

15β-Hydroxypregnenolone (**15**). Yield: 1.5 mg (4.5 μmol, 7%)
$^{13}$C-NMR (126 MHz, CDCl3): δ = 208.5, 141.0, 121.2, 71.9, 70.7, 64.0, 61.4, 50.4, 43.7, 42.4, 40.4, 37.4, 36.8, 36.0, 31.8, 31.6, 31.2, 27.9, 21.2, 19.4, 15.9 ppm. (**Fig. S15** and **S16**)

7β,15β-Dihydroxypregnenolone (**16**). Yield: 0.6 mg (1.7 μmol, 3%). (NMR data see **Table S02**, **Fig. S17–22**)

14β-Hydroxyisopregnanolone (**18**). Yield: 1.3 mg (3.9 μmol, 6%). (NMR data see **Table S03**, **Fig. S23–30**)

15β-Hydroxyisopregnanolone (**19**). Yield: 1.3 mg (3.9 μmol, 6%).
$^{13}$C NMR (126 MHz, CDCl3): δ = 208.5, 71.4, 70.8, 64.1, 61.2, 54.8, 45.1, 44.0, 40.7, 38.3, 37.2, 36.0, 35.8, 31.8, 31.6, 31.5, 31.4, 28.6, 21.3, 16.2, 12.4 ppm. (**Fig. S31** and **S32**)
$^{13}$C NMR (176 MHz, MeOH-$d_3$): δ = 211.3, 71.8, 70.9, 64.8, 62.0, 55.9, 46.2, 44.8, 41.5, 38.8, 38.1, 36.7, 36.6, 32.8, 32.2, 32.1, 31.4, 29.7, 22.2, 16.3, 12.6 ppm. (**Fig. S33** and **S34**)

7β,15β-Dihydroxyisopregnanolone (**20**). Yield: 1.3 mg (1.7 μmol, 3%). (NMR data see **Table S04**, **Fig. S35–40**)

15β-Hydroxyprogesterone (**23**). Yield: 1.2 mg (3.6 μmol, 6%).
$^{13}$C NMR (126 MHz, MeOH-$d_3$): δ = 211.1, 202.2, 175.0, 124.0, 70.7, 64.6, 61.3, 55.3, 44.6, 41.0, 40.0, 36.7, 36.7, 34.6, 33.7, 32.8, 32.2, 31.4, 21.9, 17.5, 16.1 ppm. (**Fig. S41**)
$^{13}$C NMR (126 MHz, CDCl3): δ = 208.19, 199.56, 170.88, 124.18, 70.51, 63.86, 60.51, 54.18, 43.76, 40.32, 38.86, 36.40, 35.95, 34.12, 32.83, 31.90, 31.54, 31.29, 21.10, 17.45, 16.09 ppm. (**Fig. S42** and **S43**)

**Table S03**. NMR data of 7β,15β-dihydroxypregnenolone (**16**) in MeOH-$d_3$ recorded at 298 K.

| C[a] | type | $^{13}C$[b] | $^{1}H$[b] |
|---|---|---|---|
| 1 | $CH_2$ | 38.0 | 1.88 (dt, $J$ = 13.4, 3.6 Hz, 1H) |
| | | | 1.09 (m, 1H) |
| 2 | $CH_2$ | 32.2 | 1.82 (m, 1H) |
| | | | 1.51 (m, 1H) |
| 3 | CH | 72.1 | 3.42 (m, 1H) |
| 4 | $CH_2$ | 42.5 | 2.28 (m, 2H) |
| 5 | Cq | 144.4 | – |
| 6 | CH | 126.0 | 5.26 (m, 1H) |
| 7 | CH | 73.0 | 3.99 (m, 1H) |
| 8 | CH | 37.2 | 1.82 (m, 1H) |
| 9 | CH | 49.6 | 1.17 (td, $J$ = 12.0, 4.7 Hz, 1H) |
| 10 | Cq | 37.5 | – |
| 11 | $CH_2$ | 22.1 | 1.68 (m, 1H) |
| | | | 1.55 (m, 1H) |
| 12 | $CH_2$ | 40.5 | 2.00(dt, $J$ = 12.2, 3.5 Hz, 1H) |
| 13 | Cq | 44.5 | – |
| 14 | CH | 62.2 | 1.24 (dd, $J$ = 11.2, 5.8 Hz, 1H) |
| 15 | CH | 72.0 | 4.39 (m, 1H) |
| 16 | $CH_2$ | 34.2 | 2.18 (m, 2H) |
| 17 | Cq | 64.6 | 2.59 (t, J = 9.5 Hz, 1H) |
| 18 | $CH_3$ | 15.5 | 0.85 (s, 3H) |
| 19 | $CH_3$ | 19.4 | 1.09 (s, 3H) |
| 20 | Cq | 210.9 | – |
| 21 | $CH_3$ | 31.4 | 2.13 (s, 3H) |

[a] Carbon numbering is shown in **Fig. S04**. [b] Chemical shifts δ in ppm, multiplicity: s = singlet, d = doublet, t = triplet, m = multiplet.

**Table S04**. NMR data of 14β-hydroxyisopregnanolone (**18**) in MeOH-$d_3$ recorded at 298 K.

| C[a] | type | $^{13}$C[b] | $^{1}$H[b] |
|---|---|---|---|
| 1 | $CH_2$ | 38.2 | 1.74 (m, 1H) |
| | | | 1.00 (m, 1H) |
| 2 | $CH_2$ | 38.7 | 1.54 (m ,1H) |
| | | | 1.24(m, 1H) |
| 3 | CH | 71.8 | 3.51 (tt, *J* = 10.6, 4.6 Hz, 1H) |
| 4 | $CH_2$ | 32.0 | 1.75 (m, 1H) |
| | | | 1.39 (m, 1H) |
| 5 | CH | 45.7 | 1.10 (m, 1H) |
| 6 | $CH_2$ | 29.8 | 1.31 (m, 1H) |
| | | | 1.25 (m, 1H) |
| 7 | $CH_2$ | 28.8 | 2.04 (m, 1H) |
| | | | 1.07 (m, 1H) |
| 8 | CH | 41.3 | 1.56 (m, 1H) |
| 9 | CH | 50.7 | 0.93 (m, 1H) |
| 10 | Cq | 36.7 | – |
| 11 | $CH_2$ | 21.8 | 1.51 (m ,1H) |
| | | | 1.27 (m, 1H) |
| 12 | $CH_2$ | 40.1 | 1.54 (m, 1H) |
| | | | 1.36 (m, 1H) |
| 13 | Cq | 50.2 | – |
| 14 | Cq | 86.7 | – |
| 15 | $CH_2$ | 34.4 | 2.13 (m, 1H) |
| | | | 1.70 (m, 1H) |
| 16 | $CH_2$ | 25.5 | 2.00 (m, 1H) |
| | | | 1.89 (m, 1H) |
| 17 | CH | 63.6 | 2.92 (dd, *J* = 9.6, 4.8 Hz, 1H) |
| 18 | $CH_3$ | 15.6 | 0.94 (s, 3H) |
| 19 | $CH_3$ | 12.5 | 0.82 (s, 3H) |
| 20 | Cq | 219.5 | – |
| 21 | $CH_3$ | 32.5 | 2.23 (s, 3H) |

[a] Carbon numbering is shown in **Fig. S05**. [b] Chemical shifts δ in ppm, multiplicity: s = singlet, d = doublet, t = triplet, m = multiplet.

**Table S05**. NMR data of 7β,15β-dihydroxypregnenolone (**20**) in MeOH-$d_3$ recorded at 298 K.

| C[a] | type | $^{13}$C[b] | $^{1}$H[b] |
|---|---|---|---|
| 1 | $CH_2$ | 37.9 | 1.75 (1H, m) |
| | | | 0.98 (1H, m) |
| 2 | $CH_2$ | 38.4 | 1.53 (dd, *J* = 12.3, 2.5 Hz, 1H) |
| | | | 1.30 (m, 1H) |
| 3 | CH | 71.6 | 3.50 (m, 1H) |
| 4 | $CH_2$ | 32.0 | 1.78 (1H, m) |
| | | | 1.38 (1H, m) |
| 5 | CH | 42.9 | 1.20 (m, 1H) |
| 6 | $CH_2$ | 39.3 | 1.58 (ddd, *J* = 12.6, 5.3, 2.8 Hz, 1H) |
| | | | 1.36 (m, 1H) |
| 7 | CH | 75.0 | 3.48 (m, 1H) |
| 8 | CH | 40.3 | 1.76 (1, m) |
| 9 | CH | 53.9 | 0.84 (m, 1H) |
| 10 | Cq | 35.9 | – |
| 11 | $CH_2$ | 22.4 | 1.67 (m, 1H) |
| | | | 1.40 (m, 1H) |
| 12 | $CH_2$ | 41.1 | 1.97 (m, 1H) |
| | | | 1.41 (m, 1H) |
| 13 | Cq | 45.1 | – |
| 14 | CH | 62.1 | 1.21 (m, 1H) |
| 15 | CH | 72.6 | 4.37 (ddd, *J* = 8.2, 5.6, 2.9 Hz, 1H) |
| 16 | $CH_2$ | 34.1 | 2.17 (m, 2H) |
| 17 | Cq | 64.6 | 2.58 (t, *J* = 9.5 Hz, 1H) |
| 18 | $CH_3$ | 16.0 | 0.86 (s, 3H) |
| 19 | $CH_3$ | 12.6 | 0.87 (s, 3H) |
| 20 | Cq | 211.1 | – |
| 21 | $CH_3$ | 31.4 | 2.12 (s, 3H) |

[a] Carbon numbering is shown in **Fig. S06**. [b] Chemical shifts δ in ppm, multiplicity: s = singlet, d = doublet, t = triplet, m = multiplet.

**Table S07**. LC–MS features increased following *CcCYP9Z273* gene silencing.

| RT (min) | m/z | Annotation |
|---|---|---|
| 3.14 | 275.15991 | Unassigned LC–MS feature |
| 3.21 | 245.18573 | Unassigned LC–MS feature |
| 3.22 | 222.11552 | Unassigned LC–MS feature |
| 3.44 | 265.15445 | Unassigned LC–MS feature |
| 3.52 | 107.04846 | Unassigned LC–MS feature |
| 3.53 | 245.18575 | Unassigned LC–MS feature |
| 3.56 | 518.14467 | Unassigned LC–MS feature |
| 3.73 | 245.18571 | Unassigned LC–MS feature |
| 3.77 | 304.16527 | Unassigned LC–MS feature |
| 3.77 | 237.12312 | Unassigned LC–MS feature |
| 3.79 | 338.08367 | Unassigned LC–MS feature |
| 3.82 | 243.16930 | Unassigned LC–MS feature |
| 4.02 | 279.17007 | Unassigned LC–MS feature |
| 4.16 | 245.18598 | Unassigned LC–MS feature |
| 4.18 | 318.18090 | Unassigned LC–MS feature |
| 4.23 | 279.16989 | Unassigned LC–MS feature |
| 4.37 | 272.13133 | Unassigned LC–MS feature |
| 4.39 | 274.14698 | Unassigned LC–MS feature |
| 4.60 | 181.12221 | Unassigned LC–MS feature |
| 4.64 | 259.20135 | Unassigned LC–MS feature |
| 4.68 | 267.12010 | Unassigned LC–MS feature |
| 4.73 | 259.20119 | Unassigned LC–MS feature |
| 5.11 | 149.09565 | Unassigned LC–MS feature |
| 5.11 | 107.08488 | Unassigned LC–MS feature |
| 6.59 | 181.12220 | Unassigned LC–MS feature |
| 6.71 | 237.14825 | Unassigned LC–MS feature |
| 6.84 | 331.08106 | Unassigned LC–MS feature |
| 6.89 | 271.05979 | Unassigned LC–MS feature |
| 7.07 | 301.07069 | Unassigned LC–MS feature |
| 7.94 | 263.16370 | Unassigned LC–MS feature |
| 8.16 | 318.30028 | Unassigned LC–MS feature |
| 8.29 | 123.07989 | Unassigned LC–MS feature |
| 8.29 | 151.11143 | Unassigned LC–MS feature |
| 8.77 | 191.14315 | Unassigned LC–MS feature |
| 8.88 | 359.11248 | Unassigned LC–MS feature |
| 9.99 | 299.23703 | Putative dehydrated pregnenolone ion, $[M-H_2O+H]^+$ |
| 9.99 | 317.24751 | Pregnenolone (9), $[M+H]^+$ |
| 11.01 | 205.19516 | Unassigned LC–MS feature |
| 11.21 | 219.17429 | Unassigned LC–MS feature |

“Unassigned LC–MS feature” denotes an ion for which no structural assignment was made. Accurate mass alone is insufficient to establish or exclude a steroidal origin. Features sharing an RT may represent ions derived from the same compound rather than distinct metabolites

**Table S08**. LC–MS features decreased following *CcCYP9Z273* gene silencing.

| RT (min) | m/z | Annotation |
|---|---|---|
| 4.66 | 123.05478 | Unassigned LC–MS feature |
| 5.54 | 553.30159 | Putative cardenolide hexoside, $[M+H]^+$ |
| 5.54 | 355.22665 | Putative cardenolide aglycone fragment after loss of two $H_2O$ molecules |
| 5.54 | 373.23765 | Putative cardenolide aglycone fragment after loss of $H_2O$ |
| 5.54 | 391.24846 | Putative protonated cardenolide aglycone ion |
| 6.00 | 375.25339 | Putative protonated cardenolide aglycone ion |
| 6.00 | 339.23214 | Putative cardenolide aglycone fragment after loss of two $H_2O$ molecules |
| 6.00 | 537.30685 | Putative cardenolide hexoside, $[M+H]^+$ |
| 8.93 | 242.21131 | Unassigned LC–MS feature |
| 8.94 | 228.19591 | Unassigned LC–MS feature |

"Unassigned LC–MS feature" denotes an ion for which no structural assignment was made. Accurate mass alone is insufficient to establish or exclude a steroidal origin. Putative cardenolide ion assignments are based on shared RTs and mass differences consistent with loss of a hexosyl residue (162.0528 Da) and/or $H_2O$ (18.0106 Da). These assignments do not establish aglycone stereochemistry, sugar identity or glycosylation site. Features sharing an RT may represent ions derived from the same compound rather than distinct metabolites

**Table S09**. Selected docking poses of pregnenolone in WT and G337A *Cc*CYP9Z273

| Variant | Pose | Role | Cluster/member | Score | SP-dG | C14-Fe (Å) | C15-Fe (Å) |
|---|---|---|---|---|---|---|---|
| WT | 1 | Top-ranked (non-productive) | 0/1 | -2.3943 | -8.0817 | 9.14 | 10.53 |
| WT | 190 | Productive-like | 37/1 | 14.5293 | -7.1534 | 5.53 | 5.72 |
| G337A | 1 | Top-ranked (non-productive) | 0/1 | -3.7134 | -8.0040 | 10.71 | 11.54 |
| G337A | 9 | Productive-like | 1/1 | -1.1620 | -7.7934 | 5.73 | 4.37 |

Score denotes SwissDock MEMBER_SCORE. The top-ranked docking solutions are included to illustrate that docking score alone did not identify catalytically plausible C14/C15 orientations. WT pose190 and G337A pose9 were selected as representative productive-like geometries based on substrate-heme geometry together with cluster representation, docking score and steric plausibility.
